# Metabolite interactions in the bacterial Calvin cycle and implications for flux regulation

**DOI:** 10.1101/2022.03.15.483797

**Authors:** Emil Sporre, Jan Karlsen, Karen Schriever, Johannes Asplund Samuelsson, Markus Janasch, Linnéa Strandberg, David Kotol, Luise Zeckey, Ilaria Piazza, Per-Olof Syrén, Fredrik Edfors, Elton P. Hudson

## Abstract

Metabolite-level regulation of enzyme activity is important for microbes to cope with environmental shifts. Knowledge of such regulations can also guide strain engineering to improve industrial phenotypes. Recently developed chemoproteomics workflows allow for genome-wide detection of metabolite-protein interactions that may regulate pathway activity. We applied limited proteolysis small molecule mapping (LiP-SMap) to identify and compare metabolite-protein interactions in the proteomes of two cyanobacteria and two lithoautotrophic bacteria that fix CO_2_ using the Calvin cycle. Clustering analysis of the hundreds of detected interactions showed that some metabolites interacted in a species-specific manner, such as interactions of glucose-6-phosphate in *Cupriavidus necator* and of glyoxylate in *Synechocystis sp* PCC 6803. These are interpreted in light of the different central carbon conversion pathways present. Metabolites interacting with the Calvin cycle enzymes fructose-1,6/sedoheptulose-1,7-bisphosphatase (F/SBPase) and transketolase were tested for effects on catalytic activity *in vitro*. The Calvin cycle intermediate glyceraldehyde-3-phosphate activated both *Synechocystis* and *Cupriavidus* F/SBPase, which suggests a feed-forward activation of the cycle in both photoautotrophs and chemolithoautotrophs. In contrast to the stimulating effect in reduced conditions, glyceraldehyde-3-phosphate inactivated the *Synechocystis* F/SBPase in oxidized conditions by accelerating protein aggregation. Thus, metabolite-level regulation of the Calvin cycle is more prevalent than previously appreciated and may act in addition to redox regulation.

## Main

Metabolite-protein interactions, such as allosteric inhibition or activation of enzymes, can act as feedback mechanisms for adapting metabolic flux to changing conditions [1, 2], and can also be targets for metabolic engineering [3]. Interaction proteomics techniques have been developed that can detect metabolite-protein interactions [4]. Limited proteolysis-coupled mass spectrometry (LiP-MS) detects changes in a protein’s susceptibility to digestion by proteinase K which may occur when a protein undergoes conformational change or binds to other proteins or effectors. In LiP-SMap (limited proteolysis-small molecule mapping), interactions between proteins and metabolites in cell extracts are revealed by comparison of digestion patterns with or without the added metabolite. To demonstrate and benchmark LiP-SMap, Piazza et al. treated yeast and *E. coli* extracts with metabolites [5]. Hundreds of metabolite-protein interactions were detected that were not previously known, and in many cases the altered peptides could be mapped near the enzyme active site.

An unexplored application area for interaction proteomics is the Calvin cycle, present in diverse bacteria, eukaryotic algae and plants. Redox regulation of enzyme activity in the plant Calvin cycle and the surrounding network has been studied [6–8] and some metabolite regulation of key enzymes are incorporated into models of photosynthesis [9–11]. In contrast, post-translational regulation of the Calvin cycle in bacteria is less characterized.

The bacterial Calvin cycle is of biotechnological interest as cyanobacteria and chemoautotrophic bacteria have been modified to produce biochemicals from carbon dioxide using sunlight, electricity, or hydrogen as energy sources [12–15]. The Calvin cycle is susceptible to instability at branch points where intermediates are drained, and the kinetic parameters of cycle enzymes and branching enzymes are constrained [16, 17]. Modulation of enzyme kinetic parameters (K_M_, K_i_, k_cat_, Hill coefficient) such as by allosteric or competitive effectors, could affect the rate of light-mediated activation/deactivation or cycle stability.

The Calvin cycle is present in all cyanobacteria, and in approximately 7% of non-cyanobacterial genomes [18]. Among cyanobacteria, Calvin cycle enzyme sequences have significant homology, though potential metabolite-level regulation may be different. For example, comparisons of the transcriptomic response to changes in inorganic carbon supply suggest that the cyanobacterium *Synechocystis* sp. PCC 6803 responds primarily through biochemical regulation of enzyme fluxes, while *Synechococcus elongatus* PCC 7942 responds at the level of transcription [19–21]. In chemolithoautotrophs, the Calvin cycle is frequently acquired through horizontal gene transfer and may provide a growth advantage in environments poor in organic substrates due to improved cofactor recycling, or in environments with mixed or fluctuating carbon sources [22–24]. Conservation of metabolite-level regulation in the Calvin cycle across bacterial families would imply core design principles for its operation, while differences may indicate adaptations specific to a certain microbial lifestyle or evolutionary trajectory[25].

Here, we applied the LiP-SMap technique to uncover new regulatory metabolite interactions with central carbon metabolism enzymes in four bacterial strains containing the Calvin cycle, *Synechocystis sp*. PCC 6803, *Synechococcus* PCC 7942, *Cupriavidus necator* (formerly *Ralstonia eutropha*), and *Hydrogenophaga pseudoflava. Synechocystis* is a model for studying photosynthesis, particularly because it can also metabolize glucose [26].

*Synechococcus* is an obligate photoautotroph and a model for the circadian rhythm [27]. *Cupriavidus necator* and *Hydrogenophaga pseudoflava* are chemoautotrophic betaproteobacteria in the order Burkholderiales [28–31]. LiP-SMap revealed species-specific interaction patterns for several tested metabolites, such as GAP, G6P and glyoxylate, which indicates that enzyme regulation by these metabolites may differ between autotrophic organisms. Complementary *in vitro* assays showed that GAP increases the catalytic activity of both *Synechocystis* and *Cupriavidus necator* F/SBPase in reducing conditions, suggesting a conserved feed-forward activation mechanism in the Calvin cycle.

## Results

### Assessment of LiP-SMap method and data

The LiP-SMap protocol developed by Piazza et al. and previously applied to *E. coli* was applied to four autotrophic bacteria here, with some modifications [5]. Cell cultures were harvested, and extracted proteomes were filtered to remove endogenous metabolites (reducing amounts by >90%), and resuspended in a buffer containing 1 mM MgCl_2_. The metabolite under study was added to four aliquots of the proteome extract at two different concentrations, while buffer was added to another four aliquots as negative controls.

Extracts were then digested partially by proteinase K, followed by tryptic digestion with a mixture of trypsin and LysC. Peptides were subsequently quantified using liquid chromatography-mass spectrometry (**Figure 1**). Any peptide that was differentially abundant (q < 0.01) in the metabolite-treated condition versus the control condition was assigned as a metabolite interaction. Proteins with at least one metabolite-interacting peptide were assigned as a metabolite-interacting protein

**Figure 1.**
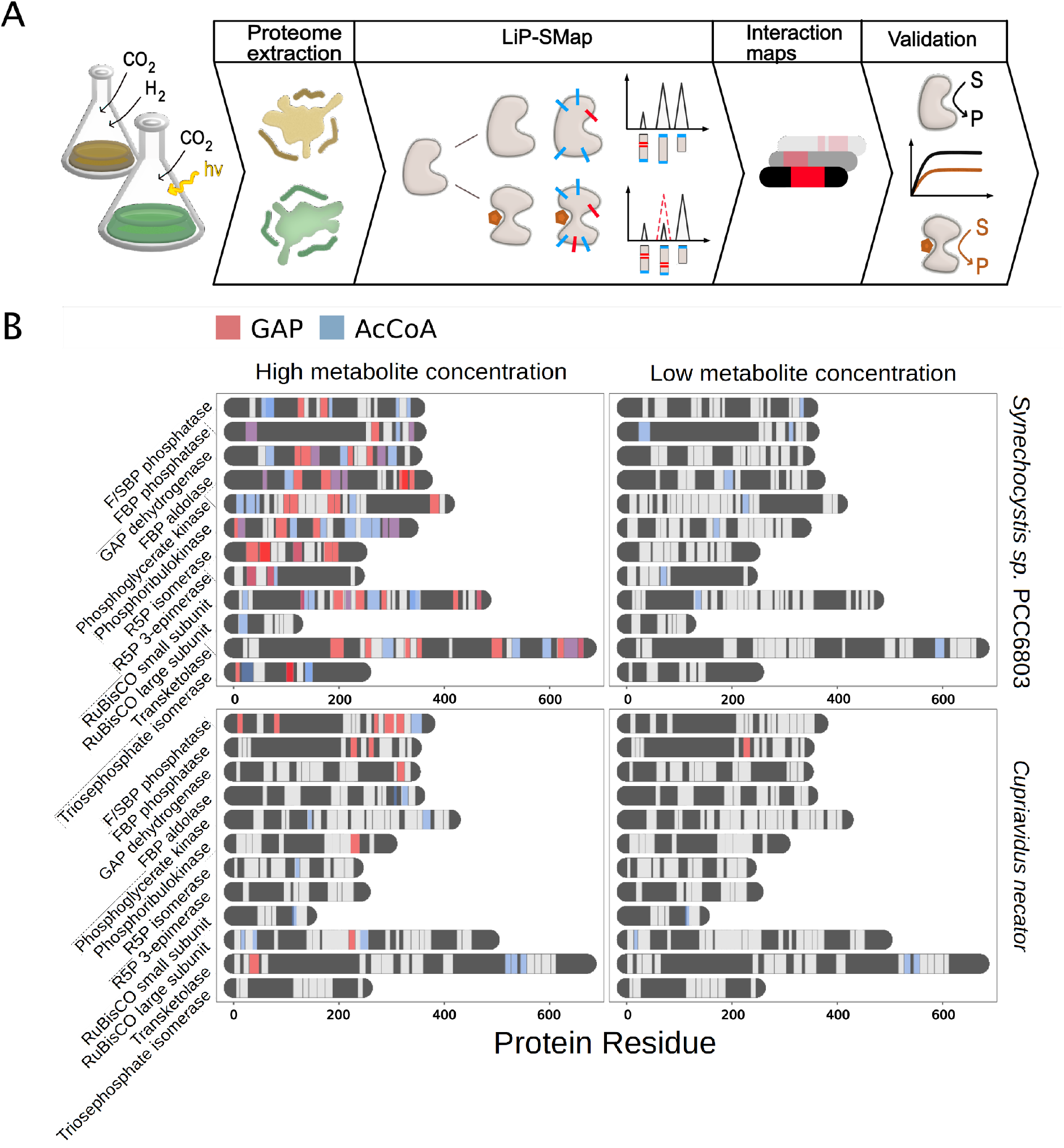
A) Workflow of interaction proteomics. Blue ticks represent digestion by trypsin/LysC and red ticks represent digestion by proteinase K. Peptides cut by proteinase K are not detected and differences in such digestion cause differentially abundant peptides. B) Peptide coverage of Calvin cycle enzymes. LiP-SMap-detected peptides from Calvin cycle enzymes of *Synechocystis* and *Cupriavidus* are colored in light gray. Regions of proteins with no peptide coverage are colored in dark gray. Significantly altered peptides when extracts were treated with GAP and AcCoA are colored in red or blue. For high concentration tests, 5 mM GAP or 10 mM AcCoA was added. For low concentration tests, 0.5 mM GAP or 1 mM AcCoA was added.

To assess the capability of LiP-SMap to detect changes in protein structure, we first tested the effect of reducing and oxidizing agents (Dithiothreitol (DTT) and 5,5′-dithiobis-2-nitrobenzoic acid (DTNB), respectively) on the extracted proteome of *Synechocystis*. Addition of DTT to 1 mM resulted in altered peptides in 21 proteins, while addition of DTNB to 50 μM altered peptides from 129 proteins, including known redox-sensitive enzymes PRK, F/SBPase, and Rubisco (**Figure S1, Dataset S1**). These results indicate that LiP-SMap can detect the changes in protein structure resulting from reducing and oxidizing agents, and that *Synechocystis* proteome extracts are partially oxidized.

A total of 8,000-15,000 peptides were detected in each LiP-SMap experiment (**Figure S2, Dataset S2**), and a higher number of peptides were detected in metabolite-interacting proteins than in non-interacting proteins (**Figure S3**). The peptide coverage of Calvin cycle enzymes was generally high, averaging 14 peptides per enzyme (minimum 4, maximum 37), with a sequence coverage of approximately 50% (**Figure 1**). To gauge the technical reproducibility of LiP-SMap, we compared the results from two consecutive LiP-SMap experiments on the same *Synechococcus* lysate treated with 10 mM glyoxylate. Out of 155 peptides significantly affected in at least one experiment, 48 were mutual. The 35-65% overlap of significant peptides among replicates is in line with what has been reported previously for MS-based proteomics [32]. While the overlap in significant peptides was only moderate, the log2FC of thosesignificant peptides was highly correlated across the two glyoxylate treatments (r = 0.88), whereas log2FC values for all peptides was lower (r = 0.49) (**Figure S4**).

### Interactions of metabolites with Calvin cycle and surrounding enzymes

Metabolites were screened for interactions with proteome extracts from *Synechocystis* (24 metabolites tested), *Synechococcus* (21), *Cupriavidus* (23), *and Hydrogenophaga* (8). These bacteria use the Calvin cycle to fix CO_2_ but differ in terms of phylogeny, energy source, substrate utilization, and natural habitat. Metabolites were chosen based on their potential to act as a regulatory signal or represent the energy, redox or metabolic status of the cell (e.g. metabolites located at end points or branch points of metabolic pathways). For each metabolite, we chose two concentrations, typically 1 mM and 10 mM (**Tables S1-S2**). The high metabolite concentrations were intended to mimic spikes in metabolite levels that occur during environmental shifts and metabolic perturbations, which may require rapid regulation of enzyme activity [33–36]. There were more detected interactions at the high concentrations than at the low concentrations. The majority (typically > 90%) of interactions from the low-concentration treatment were also observed in the high-concentration treatment (**Figure S5**).

To compare metabolite-protein interactions between species, we extracted a list of all proteins affected by any metabolite for each strain and grouped them according to KEGG orthology groups (KOGs), including only orthology groups present in all four strains (**Figure S6, Dataset S3**). Principal component analysis (PCA) was used to cluster and compare metabolite-KOG interaction patterns between the species. Differences in interaction partners between species were observed at the high metabolite concentrations, as evidenced by separation among species on PCA plots for each metabolite (**Figure 2**). For example, in the case of GAP and acetyl-CoA, metabolites with more than 200 KOG interactions in all four species, interactions in the photoautotrophs clustered apart from those in the chemoautotrophs. G6P, an entry metabolite of the pentose phosphate (PP) and the Entner-Doudoroff (ED) pathway, showed a high number of interactions in *Cupriavidus* that clustered apart from those in other species. In contrast, some metabolite-KOG interactions were similar in all species. For example, similarity of interactions with metabolites in lower glycolysis and in the reductive branch of the tricarboxylic acid cycle (2OG, PEP and citrate) may indicate conserved regulatory mechanisms in all species. As fewer interactions were detected at the low metabolite concentrations, there was a weaker separation of the species in the PCA analysis (**Figure S7**).

**Figure 2.**
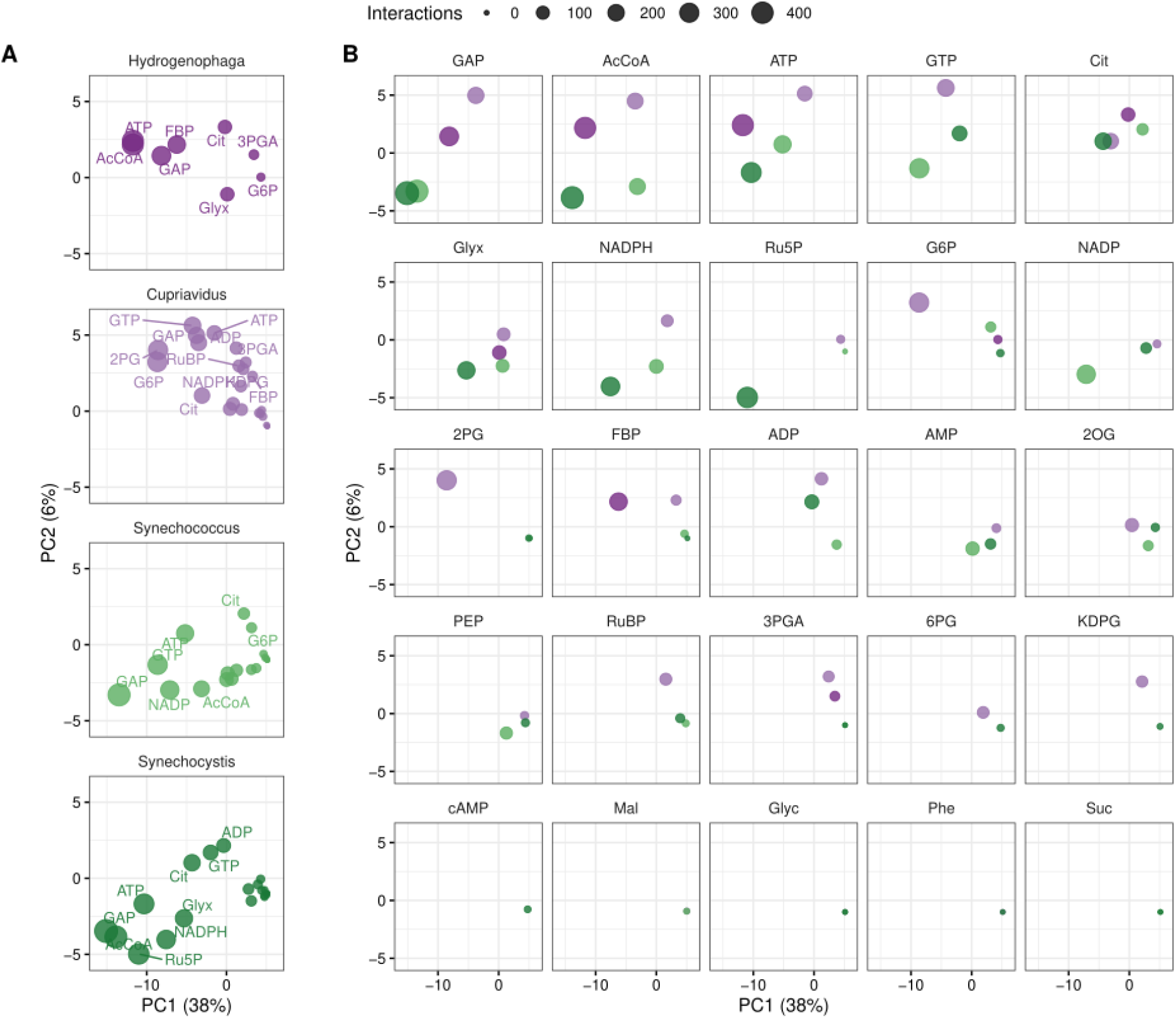
Similarity of ortholog interaction patterns, high added metabolite concentration. Principal components were calculated from the presence or absence of interaction with each of 477 orthologs (see Materials and Methods). All data points shown here are from the same principal component analysis, but split per organism (A) or metabolite (B) to reduce overplotting. Percentages indicate the fraction of the total variance captured by the principal components.

Interactions with AcCoA, ATP, citrate, and GTP were widespread in all four microbes. These metabolites have strong Mg^2+^ chelating properties [37, 38] and may sequester Mg^2+^ from proteins in extracts. Indeed, the number of ATP interactions in *Synechocystis* proteome extracts was reduced from 172 to 19 when Mg^2+^ in the LiP buffer was increased from 1 mM to 3 mM (**Figure S8**). The signaling metabolite cAMP is representative of a metabolite with few interactions, it resulted in LiP peptides from 10 proteins in *Synechocystis:* the cAMP receptor protein (SyCRP1 *sll1371* p_adj_=0.06), a regulatory protein implicated in cyclic electron flow (*slr1658* [39]), glutathione-related enzymes (Gsh *slr1238* and GlxII *sll1019* [40]), proteoloysis-related chaperones (ClpC *sll0020*, ClpB *slr0156*, GroEL *slr2076*) and an antenna core component (ApcE *slr0335*). Taken together, these interactions suggest cAMP is involved in mediating light or oxidative stress response, either by direct interaction or by inducing protein aggregation, in addition to its other diverse roles [41, 42]. However, the known cAMP-binding regulator SbtB (*slr1513*) did not show a LiP interaction. Such “false negatives,” from LiP-SMap can have several possible sources: poor coverage of the protein (e.g. SbtB had only 7 detected peptides), residual metabolite remaining in the untreated sample mixture after gel filtration, or that binding site of the metabolite is not near a proteinase K cleavage site.

Next, we specifically examined interactions of metabolites with enzymes in the Calvin cycle and enzymes in pathways that siphon carbon out of the cycle (**Figure 3, Figure S9**). Calvin cycle enzymes in these microbes are phylogenetically diverse, though the cyanobacteria enzymes are more closely related to each other than to the chemoautotroph orthologs (**Dataset S4**). Some metabolites interacted more with the Calvin cycle enzymes of certain species. For instance, the photorespiratory intermediate glyoxylate showed extensive interactions in *Synechocystis*, even at low concentrations (**Figure S9**). *Cupriavidus* Calvin cycle enzymes were particularly sensitive to intermediates of the pentose phosphate and Entner-Doudoroff pathways, such as 6PG, G6P, and KDPG, as well as 2OG and RuBP. In summary, while there were some interactions observed in all species, primarily AcCoA, ATP and GAP, most metabolites showed species-specific interactions with Calvin cycle enzymes.

**Figure 3.**
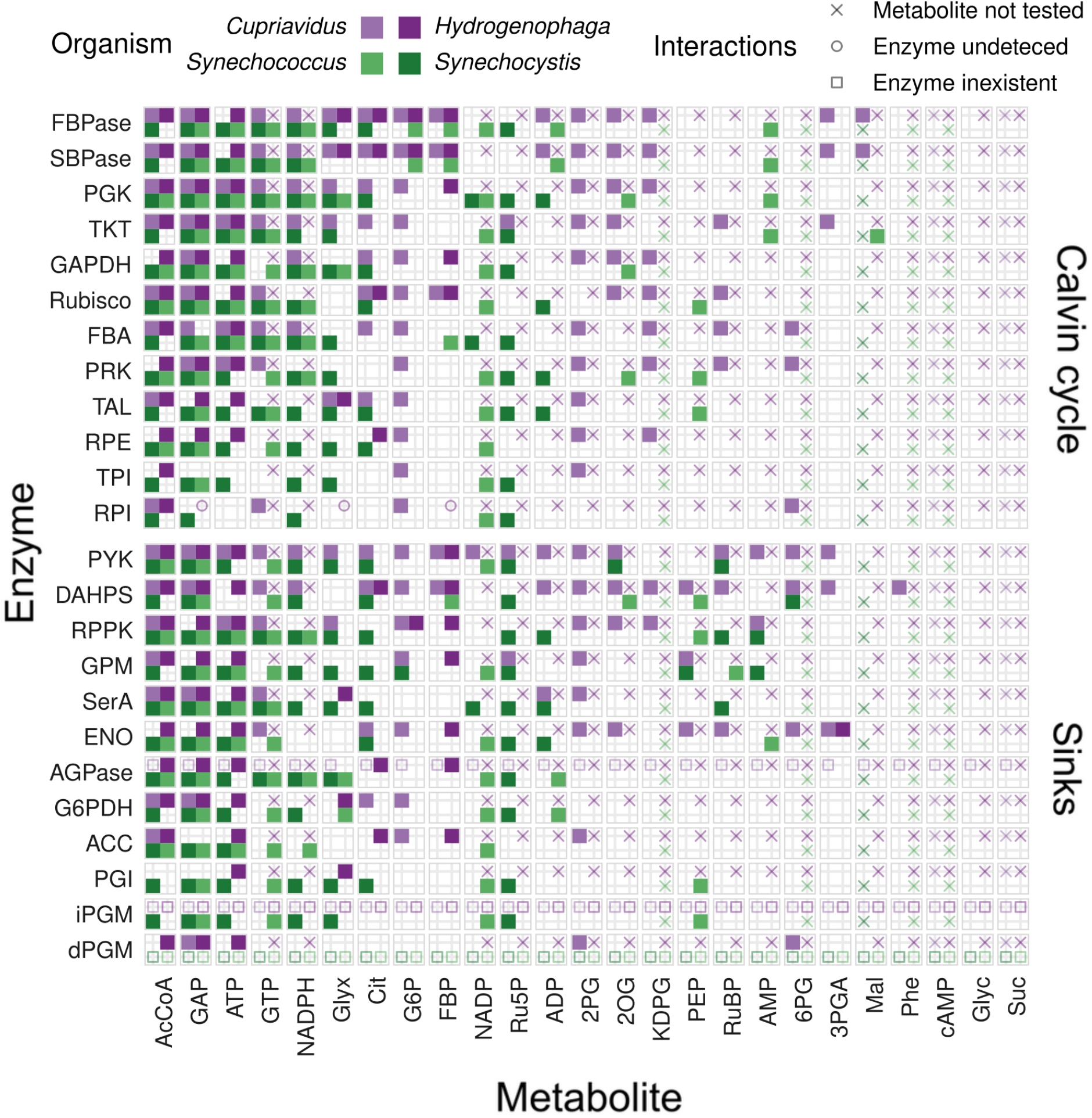
Interactions of Calvin cycle enzymes and selected central carbon metabolism enzymes with metabolites, high added metabolite concentration. Interactions between metabolites (columns) at high concentration and enzymes (rows) identified by KEGG EC number annotation are shown for each organism by tiles filled with the corresponding color. A blank tile indicates that the interaction was not detected, while missing protein data is explained by a symbol. A cross indicates that the particular condition was not measured, a circle indicates that the protein was not detected, and a square indicates that there was no such enzyme in the corresponding genome. See **Figure S9** for interactions at low metabolite concentrations.

### Validation of metabolite interactions and effect on enzyme activity

An interaction detected by LiP-SMap does not necessarily mean that the metabolite interacts directly with this protein, or that catalytic activity is modulated by this interaction. To test if metabolite-enzyme interactions identified by LiP-SMap tended to affect catalytic activity, we purified and screened two Calvin cycle enzymes from *Synechocystis* and *C. necator in vitro* that had multiple metabolite interactions in the Lip data: F/SBPase *(fbpI* and *fbp3*) and transketolase (*tktA* and *cbbTP*). F/SBPase catalyzes irreversible steps in the Calvin cycle (FBP to F6P and SBP to S7P) and F/SBPase activity has a significant effect on the rate of CO_2_ fixation in cyanobacteria in some conditions [16, 43] [44–46] Reactions far from equilibrium are more likely to be post-translationally regulated [47]. In contrast, the transketolase step of the Calvin cycle in photosynthetic microbes operates reversibly and close to equilibrium [36], though some studies have predicted that transketolase activity could have significant control over overall CO_2_ fixation rate [48] (**Figure S10)**. Therefore, metabolite-induced changes in activity of these two enzymes could be relevant for CO_2_ fixation.

Metabolites that altered the LiP pattern of *Synechocystis* F/SBPase (syn-F/SBPase) or *Cupriavidus* F/SBPase (cn-F/SBPase) were screened for their effect on kinetics using a Malachite Green assay. Additionally, the thermal unfolding of these enzymes was monitored in absence and presence of metabolites to assess any thermal shifts in melting temperature (T_m_) as indication for metabolite-induced conformational changes **Table 1, Table S3**. The enzyme was assayed in the presence of DTT, to approximate reduced conditions. Several interactions for metabolites that showed the strongest effects are discussed in more detail below. The addition of GAP at 0.5 mM stimulated syn-F/SBPase and cn-F/SBPase activity 50-70%, by reducing K_M_ (**Figure 4**). In contrast, the GAP isomer dihydroxyacetone phosphate (DHAP) did not have an effect (**Figure S11**). GAP also caused a small thermal shift of syn-F/SBPase and cn-F/SBPase. The thermal shift of syn-F/SBPase was seen at multiple Mg^2+^ concentrations, indicating that altered LiP and kinetics were caused by a direct conformational change mediated by GAP (**Figure S12-S13**). NADPH inhibited both syn-F/SBPase and cn-F/SBPase. NADPH increased the T_m_ of cn-F/SBPase, but did not strongly affect the T_m_ of syn-F/SBPase (**Figure S13-14**). The similar kinetic effects of NADPH and GAP on both F/SBPase enzymes suggest evolutionary convergence, as syn*-*F/SBPase (class II) and class I F/SBPases (for which cn-F/SBPase is a representative) have a similar monomeric fold but little sequence similarity [49, 50]. In contrast to GAP and NADPH that affected both enzymes, the addition of G6P stimulated the cn-F/SBPase up to 100%, but had no effect on the syn-F/SBPase. The specificity of the G6P effect to the cn-F/SBPase is in agreement with LiP-SMap data (**Figure S15**).

**Table 1.** Screening of LiP-SMap metabolites for effects on F/SBPase *in vitro*. Changes in kinetic parameters were determined by enzyme kinetic assays. The kinetic effect of a metabolite was considered significant for p < 0.05 (comparing kinetic parameters) and a maximum change in rate >20%. Changes in melting temperature (T_m_) greater than 2 °C were considered significant.

| Effector (mM) | LiP-SMap interaction | Kinetics change | $T_m$ change |
| --- | --- | --- | --- |
| syn-F/SBPase |  |  |  |
| AcCoA (2) | Yes | Not significant | Not significant |
| AMP (0.25) | No | Inhibit | Increase |
| Citrate (5) | Yes | Inhibit | Decrease |
| GAP, +DTT (0.5) | Not tested | Stimulate | Increase |
| GAP, -DTT (0.5) | Yes | Inhibit | Not significant |
| NADPH (3) | Yes | Inhibit | Not significant |
| G6P (2) | No | Not significant | Not tested |
| cn-F/SBPase |  |  |  |
| AMP (0.25) | No | Inhibit | Increase |
| GAP, +DTT (0.5) | Not tested | Stimulate | Not significant |
| GAP, -DTT (0.5) | Yes | Not tested | Not significant |
| NADPH (3) | Yes | Inhibit | Increase |
| G6P (2) | Yes | Stimulate | Not significant |

**Figure 4.**
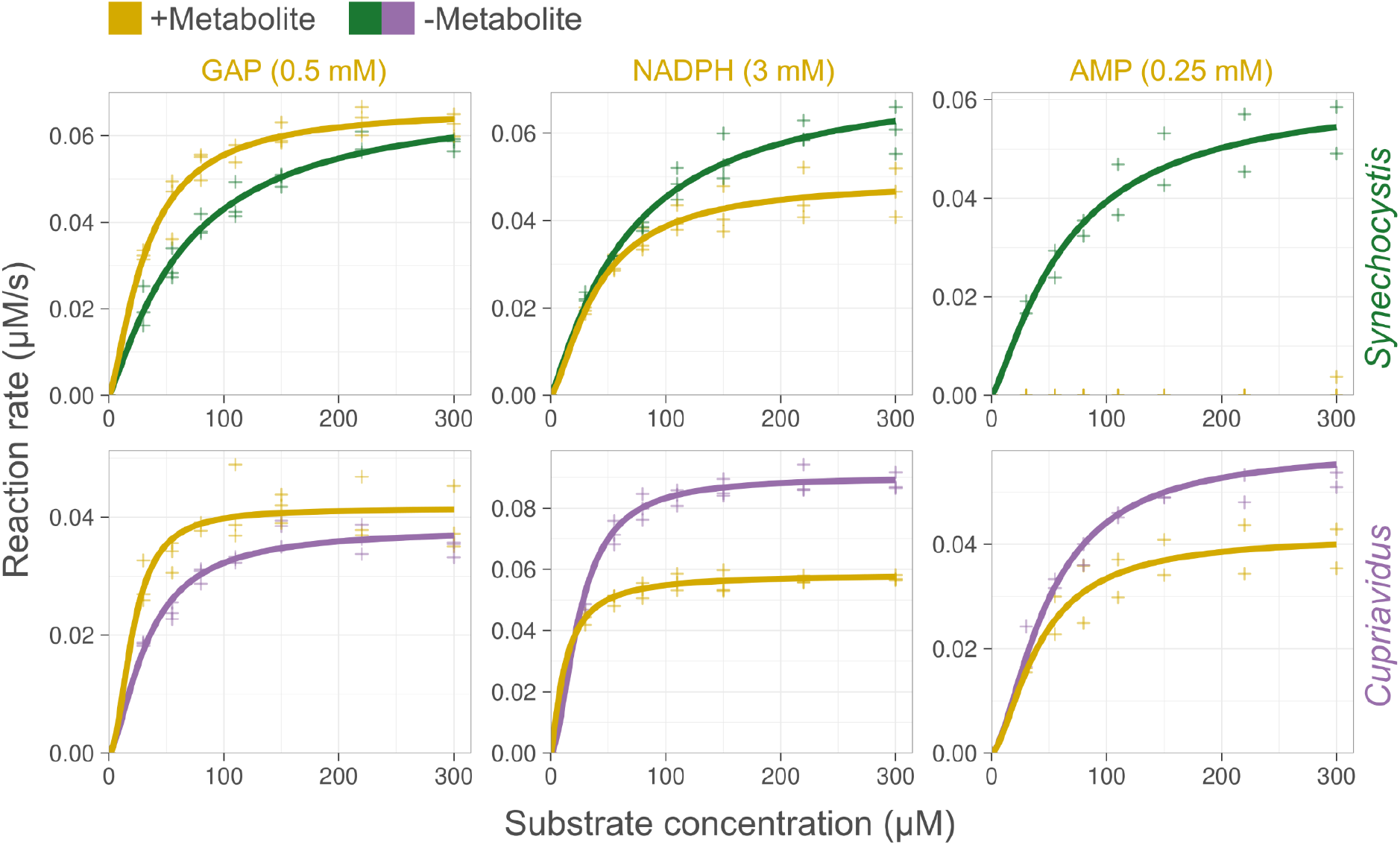
Effect of selected LiP metabolites on the activity of *Synechocystis* and *Cupriavidus* F/SBPase *in vitro*. Initial rates were measured at different substrate concentrations (FBP) in duplicates or triplicates (crosses). Lines represent data fit to the enzymatic Hill equation. The enzyme concentration was 0.42 ng/μL. No catalytic activity was detected for syn-F/SBPase treated with AMP. See also Table S3 for more metabolites.

The syn-F/SBPase is redox regulated in response to light conditions. Since GAP was the only metabolite tested that stimulated syn-F/SBPase activity, we were interested in probing whether the reductive environment (DTT in the reaction buffer) and the GAP stimulation were synergistic. In contrast to GAP’s stimulating effect in reductive conditions, we observed that GAP reduced syn-F/SBPase activity in absence of DTT (**Figure S16**). These results were verified by analyzing the same reaction mixture by Malachite Green assay and LC-MS to exclude any potential assay-based artifacts (**Figure S17**).

The syn-F/SBPase was reported to switch from a non-active initial state into the active R state upon addition of DTT [50]. We noticed that in absence of DTT a secondary syn-F/SBPase population could be observed as a peak shoulder in scattering data, which were recorded simultaneously to melting curves in the nanoDSF datasets (**Figure S16**). Simultaneously, the main peak shifted to a higher temperature. While it is not possible to identify whether the shoulder represents aggregates or a different protein state (such as different conformation or oligomeric state), it is noticeable that this population gradually disappears when increasing the concentration of DTT, suggesting that it may represent the non-active dimeric state. Indeed, the gradual decrease in the peak shoulder and shift of the main peak to the right match kinetic DTT activation data presented by Feng et al [50]. It appears that already smaller visible fractions of this shoulder correlate with a strong decrease in activity. Moreover, the fraction of this shoulder in absence of DTT increased when pre-incubating the sample at 30 C for different durations, which may either indicate aggregation or the dissociation of the active enzyme tetramer into the inactive state over time. This process was further amplified by addition of GAP during pre-incubation.

Considering that GAP contains a reactive aldehyde group and showed interactions with many proteins in Lip-SMap, an unspecific effect such as inducing protein aggregation in oxidative conditions appears plausible, explaining why GAP reduces enzyme activity in absence of DTT. The addition of GAP to enzyme treated with DTT however did not cause a peak shift in the light scattering data, indicating that the stimulating effect of GAP observed in presence of DTT likely operates by a different, more specific mechanism (**Figure S16**).

The transketolases from *Synechocystis* and *Cupriavidus* were also screened *in vitro* against all metabolites that showed a LiP-SMap interaction in any of the four species. The most prominent effects on kinetics were observed from AMP and DHAP, which specifically reduced the activity of either *syn*-TKT and *cn*-TKT, respectively (**Fig. S18-19**). While ATP and ADP inhibition of transketolases has been reported [51], inhibition by AMP has not. Overall, approximately half of the TKT-interacting metabolites detected by LiP-SMap altered TKT catalytic activity *in vitro* (8/13 for *syn-*TKT and 4/10 for *cn-*TKT), though only a few affected kinetic parameters by more than 20% or significantly affected T_m_ in thermal shift assays (**Figure S18-20**).

LiP-SMap provides peptide-level information on where metabolites interact with a protein. For syn-FBPase, both GAP and NADPH affected peptides originating from near the active site, a region distinct from the known AMP allosteric site (**Figure 5**). To confirm that GAP and NADPH regulation was separate from well-characterized AMP regulation, we created a single amino acid exchange variant (R194H) of a residue located in a β-sheet that connects the substrate-binding site to the AMP-binding site. This mutant lost AMP sensitivity, but retained sensitivity to both GAP activation and NADPH inhibition (**Figure S21**).

**Figure 5.**
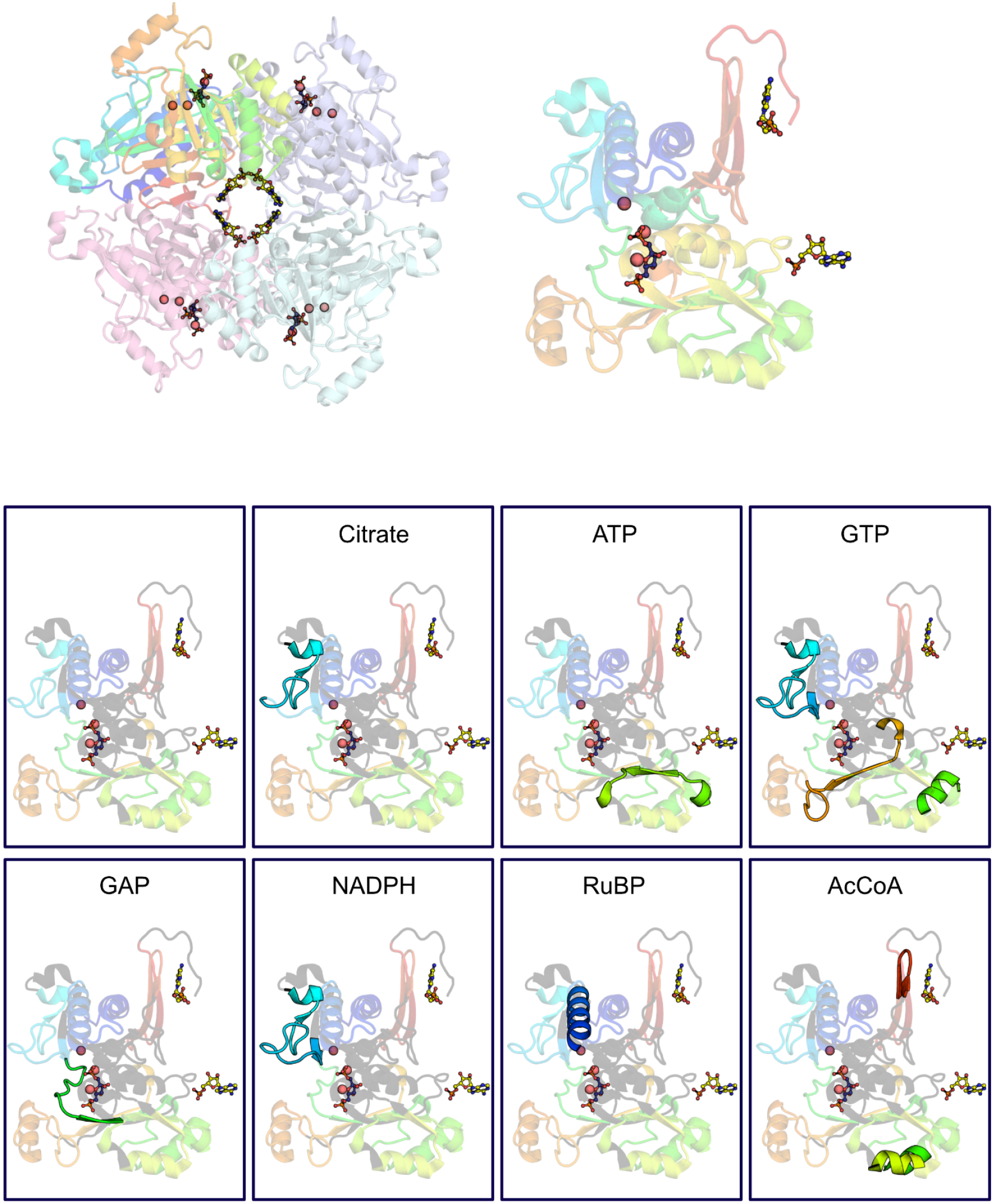
*S*tructure of *Synechocystis* F/SBPase showing peptide coverage and affected peptides from LiP-SMap. Top left: F/SBPase assembles as a homotetramer (PDB-ID: 3RPL [50]). The substrate FBP (shown as blue sticks) is coordinated by active site residues as well as Mg^2+^ ions (red spheres). AMP allosteric inhibitor molecules are located in the central interface of the tetramer (yellow sticks). Top right: monomeric view colored according to different structural elements, showing interaction with FBP and the AMP molecules at two adjacent interfaces with other monomers. Bottom: Monomeric view as shown in top right panel with peptides that were not detected in any condition shown as dark-gray ribbons. Peptides affected by the indicated metabolites (high added concentration) are outlined as opaque ribbons in individual panels.

### Predicted effects of enzyme-metabolite interactions on flux control in *Synechocystis*

We next evaluated the effects of the regulatory interactions of GAP and NADPH on F/SBPase on flux control in the Calvin cycle and metabolic stability, using ensemble modeling [16] (**Figure 6**). Two Calvin cycle models were considered, a base model with no F/SBPase regulations, and a model with NADPH inhibition and GAP activation added to F/SBPase reaction (models “Base” and “FSBPase”, respectively). For each model variant, a large set of possible metabolic states was generated from randomly sampled metabolite concentrations and enzyme kinetic parameters (V_max_, K_M_, K_i_, K_a_), each satisfying the same steady-state flux distribution (parsimonious flux balance analysis solution). The two resulting model ensembles (∼3 million models each for “Base,” and “F/SBPase”) were then assessed for system stability, which refers to the ability of the system to dynamically return to its metabolic state upon an infinitesimal small perturbation of the metabolite concentrations. The addition of regulation on F/SBPase did not alter stability significantly, with a median stability over all parameter sets of 91% and 89% for the Base and FSBPase model, respectively. Furthermore, added F/SBPase regulation did not significantly alter the metabolite concentrations at which the system tends to be more or less robust.

**Figure 6.**
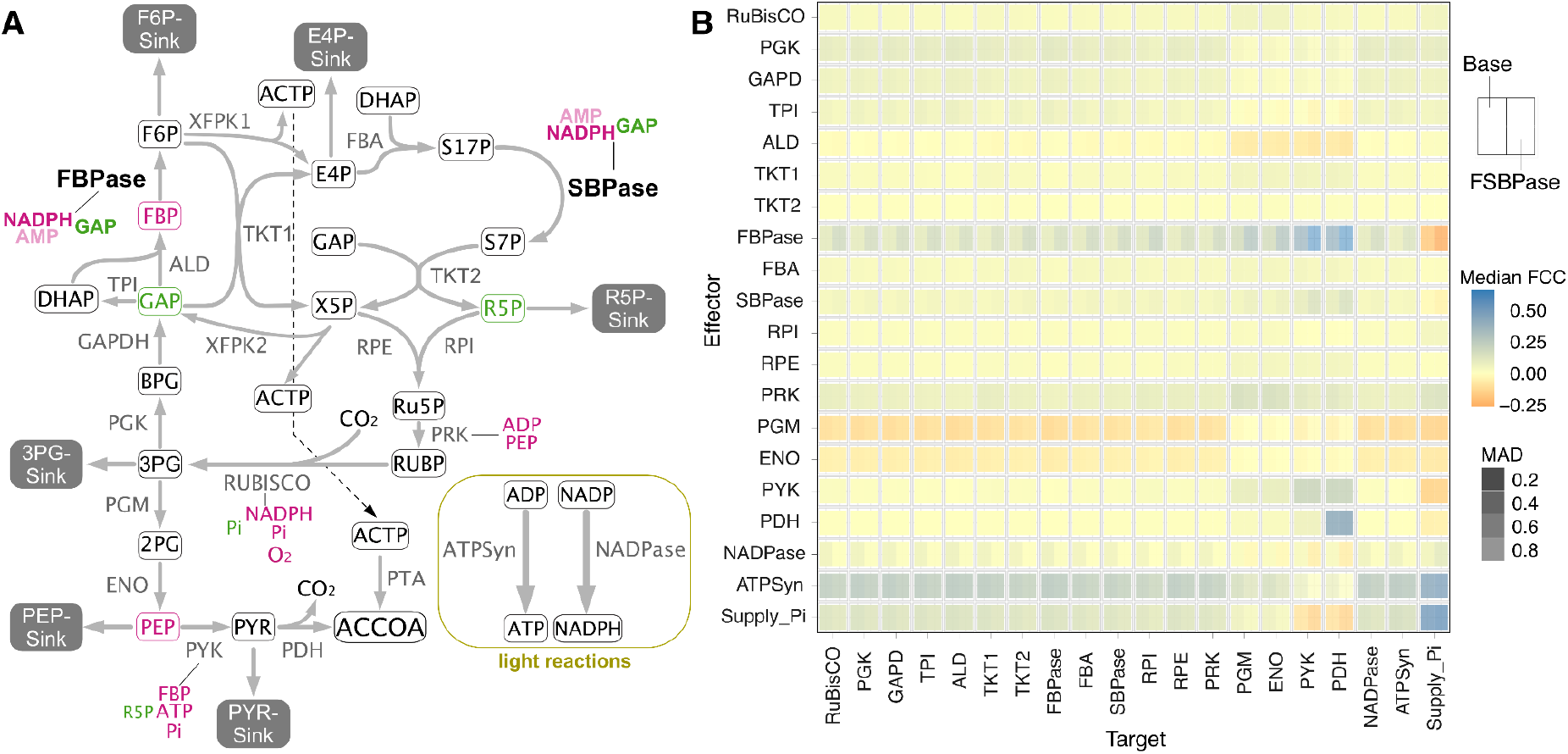
Addition of regulation on F/SBPase activity alters flux control in the Calvin cycle. A) Schematic overview of the modeled metabolism showing all included biochemical regulations. Bold: Interactions identified from LiP-SMap and verified as modulating enzyme activity. Red: inhibition, Green: activation. AMP inhibition (pink) was omitted from the model. B) Median flux control coefficients (FCCs) and median absolute deviation (MAD) over the entire model ensembles for both model variants, “Base”: no added regulation to F/SBPase, and “F/SBPase”: with GAP and NADPH regulation added.

The fully parameterized ensemble of kinetic models enables the quantification of flux control using metabolic control analysis (MCA), resulting in flux control coefficients (FCCs) for each reaction. In the Base model, the reactions supplying ATP and NADPH (e.g. light reactions in photosynthetic microbes), and supply of phosphate had positive FCCs over many other reactions, emphasizing their importance in autotrophic metabolism (**Figure S22-23** for all FCCs). While Rubisco showed positive flux control only over reactions downstream of the Calvin cycle, F/SBPase, PGK, GAPDH and PRK have flux control throughout the cycle. The F/SBPase model variant generated FCCs are similar to the Base model, but with some distinctions. Most prominently, the flux control exerted by F/SBPase over other reactions increased, signifying higher sensitivity of these reactions to F/SBPase flux. The added regulations amplify F/SBPase’s role in controlling CO_2_ fixation, as other reactions have reduced FCCs (PGK, GAPD,TPI, PRK). Flux control exerted by ATP and NADPH supply was reduced in the F/SBPase model, rendering the system with added regulation less sensitive and thereby more stable towards potential perturbations in ATP and NADPH supply. The changes in FCCs in the F/SBPase model are a result of both the direct effects of GAP and NADPH on F/SBPase, as well as the response of the whole system and the interactions between all participating entities.

## Discussion

The recently developed chemoproteomic workflow LiP-SMap was applied to reveal metabolite-level regulation of enzymes within the Calvin cycle and central carbon metabolism, comparing four autotrophic bacteria. We found that some tested metabolites interacted extensively in all organisms, such as ATP, GTP, GAP, AcCoA and citrate. Furthermore, many metabolites showed species-specific interaction patterns at high concentrations, as determined by PCA analysis of interacting orthologs, which could indicate the importance of these as metabolic signals and regulators for a species’ metabolic capacity or evolutionary adaptation. The extent of interactions at low added metabolite concentrations (0.5-1 mM) was significantly less than at high concentrations (5-10 mM). Metabolite control of enzyme activity is therefore likely not prevalent at steady state *in vivo* concentrations (< 1 mM), but rather when metabolite levels fluctuate, such as during environmental shifts or, relevant for synthetic biology, when new metabolic pathways are installed [52].

The LiP-SMap technique likely detects many interactions that do not alter the conformation of a protein, as the correlation between interaction and melting temperature shifts for F/SBPase and TKT enzymes was low. However, regulation does not always involve a conformational change. In addition to competitive inhibition or activation, it has been proposed that metabolites can affect enzyme activity through alteration of the redox potential of thiol groups, so that the degree of activation by thioredoxin (or DTT *in vitro*) is altered [11].

The Calvin cycle intermediate GAP interacted with many proteins in all species. Clustering analysis showed that a subset of GAP interactions in the photoautotrophs were different compared to those in the chemoautotrophs. GAP intersects several central pathways, such as EMP, ED, PP and the Calvin cycle, and different utilization of these pathways between species may require specific regulation by GAP. For example, variability in the abundance of GAP could regulate various “anaplerotic,” fluxes into the Calvin cycle from the ED, PP and PGI shunts [53]. Feed-forward activation of F/SBPase by GAP may work to prevent excessive GAP accumulation and increase Calvin cycle flux in response to up-shifts in energy and CO_2_ levels in the growth environment. Feed-forward regulation in glycolysis and gluconeogenesis has been reported. For example,n *E. coli* and red blood cells, pyruvate kinase is stimulated by the glycolysis intermediate FBP [54, 55]. The type I FBPase in *E. coli* is stimulated by the gluconeogenesis substrate PEP [2, 56]. In photosynthetic microbes, intracellular levels of GAP, DHAP, FBP and SBP fluctuate with light [34–36, 57, 58]

In the absence of DTT, GAP inhibits F/SBPase in a mechanism that likely involves aggregation or dissociation into an inactive state. In the cellular context, this inactivation, in addition to disulfide formation and AMP formation, could work to prevent a futile cycle forming between PFK and F/SBPase, though PFK flux in the dark is likely minimal [58, 59]. Moreover, inactivation could prevent depletion of glycolytic intermediates that are needed for resuming photosynthesis when illumination returns.

Rapid post-translational regulation of glycogen metabolism appears to be an important feature of photoautotrophic metabolism [60][61]. ADP-glucose pyrophosphorylase, which catalyzes the first committed step in starch synthesis, interacted with more metabolites in *Synechocystis* (7 metabolites) and *Synechococcus* (7) compared to the chemoautotrophs *Cupriavidus* (1) and *Hydrogenophaga* (5). As *Cupriavidus* lacks AGPase enzyme, UDP-glucose pyrophosphorylase was compared [62]. The photorespiration metabolite glyoxylate interacted with phosphoglucomutase and AGPase in the two cyanobacteria, which suggests that elevated levels of glyoxylate in response to inorganic carbon limitation may participate in the associated activation of glycogen degradation [63, 64], though we did not test this.

The enrichment of interactions of G6P and KDGP with *Cupriavidus* proteins may be related to the preferred usage of the ED pathway for sugar catabolism in this microbe,[65] as these ED intermediates may signal sugar availability. The stimulation of F/SBPase activity by G6P in *Cupriavidus* but not in *Synechocystis* can also be interpreted in light of their different glycolysis pathways. In *Cupriavidus*, the Calvin cycle operates simultaneously and parallel to the ED pathway; G6P derived from glucose does not enter the cycle [24, 66]. Stimulation of F/SBPase by G6P could accelerate re-assimilation of CO_2_ emitted during glycolysis. In *E. coli*, where EMP glycolysis and gluconeogenesis do not operate simultaneously due to overlap in metabolic pathways, the FBPase is inhibited by G6P, a regulation that effectively turns off gluconeogenesis during growth on glucose *[67]*. In *Synechocystis*, G6P derived from glucose enters the Calvin cycle *via* the PGI shunt as F6P, downstream of F/SBPase, and fluxes through EMP or ED glycolysis are negligible [68]. In this case, regulation of F/SBPase activity by G6P may not be beneficial for cell fitness.

Interactions detected by LiP-SMap are not always direct, but can arise from secondary effects of metabolite addition, such as Mg^2+^ chelation. Metabolite chelation of Mg^2+^ ions likely explains the high number of interactions observed for e.g. ATP, GTP, and citrate. The extensive interactions detected for GAP could be due to its reactivity as an aldehyde.

Glyceraldehyde was shown to form Schiff-base adducts with lysine residues on carbonmonoxyhemoglobin *in vitro [69]*. Furthermore, GAP is a source of methylglyoxal, which was shown to react with lysine, arginine, and cysteine residues on proteins, leading to advanced glycation end products [70, 71]. Indirect effects may also occur from enzymatic conversion of the metabolite, although the removal of endogenous cofactors from the proteome extracts by filtration is presumed to limit such effects; the size exclusion chromatography reduces native metabolites by >90% [5]. Even though secondary effects could be confounding, they may still be relevant for metabolism *in vivo*. For example, excessive accumulation of citrate and ATP, which could occur during nitrogen depletion, may inactivate ribosomes and anabolic processes that depend on Mg^2+^, which is particularly relevant for photosynthesis [72, 73].

Previously, inference of regulation in microbial metabolism has been done through analysis of time-resolved metabolite and proteomics datasets [2, 33], fitting of multi-omics steady state data [74, 75], or through coelution of proteins and metabolites from a chromatography column [76, 77]. By quantification of individual peptides, the LiP-SMap method can provide insight into which area of the protein changes in presence of metabolite, which provides an extra level of information compared to inference methods. However, the method is also limited by the somewhat sporadic nature of peptide detection from complex mixtures by MS; even replicate tests performed in parallel do not fully overlap with respect to which peptides are detected [32]. In this respect, LiP-SMap may work best in tandem with other interaction-proteomics techniques, such as thermal proteome profiling, which relies on quantification of proteins, not individual peptides [78]. LiP-SMAP would benefit from a more accurate peptide quantification and a wider peptide coverage for improved specificity and sensitivity, respectively.

## Materials and methods

### Cultivations and harvest

*Cupriavidus necator* strain DSMZ 428 was grown in Ralstonia Minimal Media (RMM) with 100 mM HEPES pH 7.5 under chemostat conditions in a Photon Systems Instruments Multi-Cultivator MC-1000 OD. Each reactor tube was set up to a volume of 55 mL, OD_600_ 0.05 and 3.5 g/L fructose. Once growth ceased, an inlet feed of 0.01 - 0.05 mL/min of 8 g/L formic acid in RMM with 100 mM HEPES pH 7.5 was initiated. Cultivations were kept running until a stable OD_600_ had been observed for at least 5 doubling times.

*Hydrogenophaga pseudoflava* strain DSMZ 1084 was grown at 30 °C and 200 RPM in sealed flasks of ∼135 mL containing ∼25 mL DSMZ media 133 and ∼110 mL of gas (70% H_2_, 15% CO_2_ and 15% O_2_) at 1 bar overpressure. Cultivations were started from overnight pre-cultures grown on 1.5 g/L acetate and harvested during exponential growth at OD_600_ ∼1.0.

*Synechocystis sp*. PCC 6803 (gift from Klaas Hellingwerf, University Amsterdam) & *Synechococcus elongatus* PCC7942 (from Pasteur Culture Collection, France) were grown in BG-11 media at 1% CO_2_ and a light intensity of ∼70 μmol/s·m^2^ in 500 mL flasks containing 100 mL liquid until an OD_730_ of ∼1.0.

For each microbe, four biological replicate cultivations were performed, and immediately before harvest the replicates were pooled. Cells were harvested by centrifugation and washed three times with cold lysis buffer before being resuspended in a small amount of lysis buffer, snap frozen in liquid nitrogen, and stored as aliquots in -80 °C. The cyanobacteria were exposed to light at ∼ 400 μmol·s^-1^·m^-2^ for 5 minutes prior to snap freezing.

### Proteome extraction

Frozen aliquots were thawed on ice and lysed mechanically through bead beating by a FastPrep-24 5G lysis machine over six cycles of 45 seconds at 6.5 m/s with 30 seconds on ice between cycles. The lysate was spun down and the supernatant was run through a Zeba Spin Desalting Column (size exclusion chromatography). Protein concentration in the desalted lysate was evaluated using a Bradford assay. The samples were kept at 4 °C throughout the procedure.

### Limited proteolysis

For every experiment three sample groups were created, one with no added metabolite and two with different concentrations of metabolite specified in **Table S1**. Each sample group was prepared as four technical replicates with 1 μg/μL extracted protein. Proteinase K was simultaneously added to all samples at a 1:100 protease to protein ratio and incubated at 25 °C for exactly 10 minutes before immediate denaturation. All sample groups originated from the same cell extract, were treated in parallel with the same reagent aliquots and run on the LC-MS on the same plate.

### Complete digestion

The protein mix was incubated at 96 °C for 3 min prior to treatment with 5% sodium deoxycholate and 10 mM DTT and another 10 min at 96 °C after. The samples were then alkylated by 10 mM iodoacetamide at RT for 30 min in the dark, after which proteases LysC and trypsin were applied at a 1:100 protease to protein ratio and incubated at 37 °C and 400 RPM in a thermocycler for 3 and 16 hours, respectively. Digestion was halted by addition of formic acid to reduce pH below 2 which caused sodium deoxycholate to precipitate. Samples were then centrifuged at 14,000g for 10 min after which the supernatant was removed and stored at -20 °C.

### Peptide purification

Pipette tips packed with six layers of C18 matrix discs (20-200 μL; Empore SPE Discs) were activated with acetonitrile and equilibrated with 0.1% formic acid prior to being loaded with thawed peptide mixes. The matrix was then washed twice with one loading volume of 0.1% formic acid before being eluted with a mixture of 4:1 ratio of acetonitrile to 0.1% formic acid. The eluate was stored at -20 °C until analysis by LC-MS.

### LC-MS analysis

Analysis was performed on a Q-exactive HF Hybrid Quadrupole-Orbitrap Mass Spectrometer coupled with an UltiMate 3000 RSLCnano System with an EASY-Spray ion source. 2 μL of each sample was loaded onto a C18 Acclaim PepMap 100 trap column (75 μm x 2 cm, 3 μm, 100 Å) with a flow rate of 7 μL per min, using 3% acetonitrile, 0.1% formic acid and 96.9% water as solvent. The samples were then separated on ES802 EASY-Spray PepMap RSLC C18 Column (75 μm x 25 cm, 2 μm, 100Å) with a flow rate of 3.6 μL per minute for 40 minutes using a linear gradient from 1% to 32% with 95% acetonitrile, 0.1% formic acid and 4.9% water as secondary solvent. After separation MS analysis was performed using one full scan (resolution 30,000 at 200 m/z, mass range 300 – 1200 m/z) followed by 30 MS2 DIA scans (resolution 30,000 at 200 m/z, mass range 350 – 1000 m/z) with an isolation window of 10 m/z. Precursor ion fragmentation was performed with high-energy collision-induced dissociation at an NCE of 26. The maximum injection times for the MS1 and MS2 were 105 ms and 55 ms respectively, and the automatic gain control was set to 3·10^6^ and 1·10^6^ respectively. The EncyclopeDIA and Prosit workflows were used to generate a predicted library from a fasta file of the appropriate organisms UniProt proteome set (*C. necator*: UP000008210, *Synechocystis sp. PCC 6803*: UP000001425, *Synechococcus elongatus sp. PCC 7942*: UP000002717, *H. pseudoflava*: UP000293912) against which an EncyclopeDIA search was performed to generate a list of detected peptides.

### Data analysis

Peptides detected in at least three replicates in every sample group were tested for differential peptide abundance using the MSstats package (version 3.18.5) in R (version 3.6.3). For every peptide in each metabolite concentration comparison MSstats estimated fold changes and p-values adjusted for multiple hypothesis testing (Benjamini-Hochberg method) with a significance threshold of 0.01. A protein was considered to interact with a metabolite supplied at low or high concentration if at least one peptide showed significant change. General data and quality assessment statistics and visualizations were generated by the pipeline available at https://github.com/Asplund-Samuelsson/lipsmap, implemented in R version 4.1.1 with Tidyverse version 1.3.1.

### Ortholog annotations

In order to compare metabolite-protein interaction patterns between organisms, it was necessary to determine orthologous genes. Ortholog labels from the eggNOG database were downloaded from UniProt (https://www.uniprot.org/) on 14 June 2021 for each protein in the four organisms. Version 5.0 of eggNOG was used except for proteins Q31NB2 (ENOG4108VFZ), Q31RK3 (ENOG4105KVS), and Q31RK2 (ENOG4105HKE) in *Synechococcus*, which were annotated with eggNOG version 4.1. Only the 481 orthologs found in all organisms were considered. The number of interacting proteins were counted for each ortholog and metabolite concentration, in each organism. Furthermore, ortholog counts were summarized into the 20 functional categories each represented by a single letter, *e*.*g*. “A” for “RNA processing and modification.”

### Principal component analysis of interactions with orthologs

The metabolite-protein interaction patterns of orthologs were compared between metabolites and organisms using R. The interaction per ortholog was first classified binarily so that the interaction was 1 (one) if there was at least one interaction for the ortholog in a particular combination of organism, metabolite, and concentration. Otherwise the interaction was classified as 0 (zero). Orthologs without interactions were filtered out. A matrix with rows representing organism and metabolite, and columns containing the binary interaction classification of each ortholog, was subjected to principal component analysis (PCA; function *prcomp*). The first two principal components were then plotted in order to visualize how similar different organisms and metabolites were in terms of interaction with the full set of orthologous genes. The PCA was performed separately for low and high metabolite concentrations.

### Clustered heatmap of interactions with orthologs

The metabolite-protein interaction patterns of orthologs, summarized per ortholog functional category, were further inspected through visualization with a heatmap with clustered rows and columns. The ortholog interaction counts were normalized to indicate the fraction of interacting orthologs within each combination of functional category, organism, metabolite, and concentration. These fractions were then used to calculate Euclidean distance (function *vegdist* from library *vegan*) followed by clustering (*ward*.*D2* method in function *hclust*), which determined the order of functional categories (heatmap rows), and metabolites and concentrations (heatmap columns). Organisms contributed both to row and column clustering. Finally, the ortholog interaction fractions were plotted as heatmaps, using row and column orders as described, with dendrograms clarifying the clustering (function *ggtree* from library *ggtree*).

### Phylogenetic analysis

Sequences for Calvin cycle KEGG orthologs (KO) in module M00165, supplemented with transaldolase (K00616 and K13810), triose-phosphate isomerase (K01803), and ribulose-phosphate epimerase (K01783), were downloaded from UniProt on 14 October 2021. Each set of KO sequences were reduced in number with cd-hit version 4.8.1 [79, 80] by selecting the highest percent identity setting between 50% (-c 0.5) and 100% (-c 1), in 5% steps, that resulted in fewer than 1 000 representative sequences. For each KO set, we added any missing corresponding protein sequences in the four organisms studied here. Sequences were aligned using mafft version 7.453 at default settings [81]. The alignments were then used to construct phylogenetic trees with FastTree version 2.1.11 Double precision at default settings [82]. NCBI taxonomy data downloaded on 8 October 2021 was used to identify organism groups. Trees were plotted using *phytools* and *ggtree* in R in order to visualize the phylogenetic distribution of sequences and metabolite interactions for the four organisms under study. Ortholog analysis code is available at https://github.com/Asplund-Samuelsson/lipsmap.

### Cloning and transformation

The *glpX* (*slr2094*) gene of *Synechocystis sp*. PCC 6803 and the *fpb3* (cbbFp) gene in *C. necator* were synthesized by Twist Biosciences. The genes were cloned into pET-28a(+) using Gibson assembly. The products were verified by sequencing and transformed into *E. coli* BL21 by heat shock.

The *tktA* gene in *Synechocystis sp*. PCC 6803 and the *cbbTP* gene in *C. necator* were PCR amplified using the primer pairs tktAF+tktAR and cbbTpF+cbbTpR, respectively. The backbone pET-28a(+) was linearized using the primer pair pETF+peTR after which the constructs were assembled through Gibson assembly. The products were verified by sequencing and transformed into *E. coli* BL21 by heat shock.

tktAF: 5′-CCATTTGCTGTCCACCAGACAGTGAGGAGTTTTAAGCTTGG-3′

tktAR: 5′-CCGCGCGGCAGCCATATGAACATTATGGTCGTTGCTACCC-3′

cbbTpF: 5′-CCATTTGCTGTCCACCAGATCAAGCGTCCTCCAGCAG-3′

cbbTpR: 5′-CCGCGCGGCAGCCATATGGAGATGAACGCACCCGAACG-3′

pETF: 5′-CATATGGCTGCCGCGCGG-3′

pETR: 5′-CTGGTGGACAGCAAATGGGTCG-3′

### Production and purification of recombinant F/SBPase and TKT enzymes

The mutants were cultivated in 2YT media at 37 °C and 200 RPM until OD 0.4-0.6, after which overexpression was induced by addition of 0.5 mM IPTG. The *tktA* gene as well as FBPase genes were incubated at 37 °C for 8h after induction, whereas the *cbbTP* gene was incubated at 18 °C for 24 hours. Cells were harvested by centrifugation at 4 °C and cell pellets were stored at -20 °C. Frozen pellets were resuspended in 3-5 mL of B-PER™ Complete Bacterial Protein Extraction Reagent (ThermoFischer Scientific) and incubated on a rocking table for ∼30 min before centrifugation at 4,000 g. The soluble fraction was loaded onto an HisTrap Fast Flow Cytiva column (1 mL) using an ÄKTA start protein purification system and washed with 15 column volumes of wash buffer (50 mM Tris-HCl, 500 mM NaCl, 20 mM imidazole, pH 7.5) prior to elution with a stepwise gradient of elution buffer (50 mM Tris-HCl, 500 mM NaCl, 300 mM imidazole, pH 7.5). Fractions containing protein were combined and the buffer was exchanged to storage buffer (50 mM Tris-HCl, pH 7.5(TKT), pH 8.0 (F/SBPase) using a PD-10 Cytiva desalting column. The purified protein was quantified by Bradford assay and stored at -80 °C in aliquots.

### *In vitro* kinetic assays of TKT with metabolite effectors

Transketolase was characterized following a previously published protocol [83]. The conversion of D-ribose-5-phosphate and L-erythrulose to sedoheptulose-7-phosphate and glycolaldehyde was measured through the consumption of NADH by alcohol dehydrogenase when reducing glycolaldehyde to ethylene glycol. Initially, the WT kinetics were calculated from measurements of initial rates at 12 substrate concentrations (0, 100, 200, 300, 400, 500, 600, 800, 1000, 2000, 4000, 8000 μM) in quadruplicate in absence of any metabolite. Subsequently, relative comparisons of enzyme kinetics were made as calculated from 8 different substrate concentrations (0, 100, 200, 500, 750, 1000, 2000 and 4000 μM) with and without 1 mM added metabolite. The tested metabolites were 2OG, 2PG, ATP, AMP, G6P, Citrate, Glyoxylate, Malate, NADP and DHAP. The reaction mix contained 100 mM glycylglycine buffer pH 7.5, 5 mM MgCl, 2 mM thiamine pyrophosphate, 0.5 mM NADH, 100 mM L-erythrulose, 10 U ADH, 2.875 μg/mL transketolase and D-ribose-5-phosphate to a final volume of 100 μL. Absorption was measured at 340 nm twice per minute over 30 minutes starting immediately after addition of D-ribose-5-phosphate.

### *In vitro* kinetic assays of F/SBPase with metabolite effectors

*In vitro* enzyme activity assays were conducted to validate the kinetic effect of F/SBPase metabolite interactions detected by LiP-SMap. To determine metabolite-induced changes in enzyme kinetic parameters, initial rates were measured at eight different substrate concentrations (0, 30, 55, 80, 110, 150, 220, 300 μM) in the presence and absence of a metabolite (+M and -M). Tested metabolites were GAP, NADPH, AMP, AcCoA, and citrate (0.5, 3, 0.25, 2, and 5 mM, respectively). The conversion rate of fructose-1-6-bisphosphate to fructose-6-phosphate was determined from the accumulation of inorganic phosphate over time, using a Malachite Green (MG) assay adapted from a previously published protocol [84]. MG dye stock (1.55 g/L Malachite Green oxalate salt, 3 M H_2_SO_4_) was used to prepare a fresh phosphate colorimetric development solution prior to each experiment (400 μL MG dye stock, 125 μL ammonium molybdate (60 mM), 10 μL Tween-20 (11% v/v)). The development solution was filtered through a 0.2 μm syringe filter and kept in the dark. Development plates were prepared by mixing 36 μL development solution with 100 μL reaction buffer (50 mM Tris-Hcl, 15 mM MgCl_2_, 10 mM DTT) lacking DTT. Enzyme solutions for +M and -M conditions were prepared in separate 8-tube PCR strips (VWR #732-1521 or low-protein binding) by mixing 25 μL reaction buffer (+M/-M) with 25 μL purified enzyme constituted in -M reaction buffer. The two strips were pre-incubated at 30 ºC for 12 minutes in a thermocycler together with two additional PCR strips which contained substrate at eight different concentrations in -M reaction buffer. Reactions were initiated by quickly mixing 50 μL substrate with the enzyme mixture in one of the reaction strips, using a multipipette ([F/SBPase]_Final_ = 0.42 ng/μL). A sample of 20 μL was immediately transferred to a development plate before incubating the reaction strip at 30 °C, which quenches the reaction. The initiation procedure was repeated for the second reaction strip with a two minute delay. Samples were collected after 10, 20, and 30 minutes. Each sampling event was followed by an addition of 7.5 μL sodium citrate (34% w/v) to stabilize the color of the development solution. Triplicate series of phosphate standards (0-100 μM) were added to the development plate as a reference. The plate was incubated for 20 minutes in the dark before measuring the absorbance at 620 nm in a plate reader. The experiment was replicated at least twice. To quantify the amount of phosphate, the background absorbance measured at time zero was first subtracted from raw absorbance measurements. Phosphate standard series were then used to convert absorbances to phosphate concentrations. Outliers and phosphate concentrations that were lower than 10 μM (sensitivity threshold), or that exceeded 60% substrate conversions (10-minute time points were always kept nonetheless), were removed. Reaction rates were calculated as the change in phosphate concentration over time using linear regression. To determine kinetic parameters, reaction rates and substrate concentrations were fit to the Hill equation using non-linear regression. A parameter change was considered statistically significant for p < 0.05 (Student’s t-test).

In addition, one experiment was conducted where 75 μL reaction mixture (endpoint measurement rather than rate measurement) was transferred to an equal volume of methanol and analyzed on a TSQ Altis Triple Quad mass spectrometer coupled to a Vanquish UHPLC with a HESI ion source. 10 μL of sample was loaded onto a Accucore-150-amide-HILIC column (50 mm x 2.1 mm, 2.6 μm) with a flow rate of 0.4 mL per min. The samples were separated for 5 minutes using a linear gradient from 90 % to 0 % acetonitrile with 10 mM ammonium carbonate and 0.2% ammonium hydroxide in water as a secondary solvent. The mass spec was run in negative mode with a voltage of 2500V and searched for the transitions indicated in **Table S4**.

### Thermal shift assay and scattering experiments

Thermal unfolding of proteins was measured in absence and presence of metabolites by nano differential scanning fluorimetry (F350/F330) using a Prometheus NT.48 (NanoTemper) at 95% excitation power over a temperature gradient from 20 °C to 95 °C at an increase of 1 °C per minute. Transketolase samples were prepared in 50 mM Tris-HCl pH 7.5 with 5 mM MgCl_2_, 2 mM TPP, 200 ng/μL enzyme and 1 mM of metabolite. In addition, samples with and without 2 mM TPP and 10 mM DTT were also run to assay the effect of the cofactor and reductive power on protein stability. F/SBPase samples were prepared in 50 mM Tris-HCl pH 8 with 200 ng/μL enzyme, 15 mM MgCl_2_, 10 mM DTT and varying concentrations of metabolite (**Table 1**). The effect of AcCoA, GAP and citrate (2, 0.5, and 5 mM, respectively) on syn-F/SBPase was analyzed at different MgCl_2_ concentrations to test whether T_m_ changes were caused by magnesium chelation. T_m_ changes greater than 2 °C were considered significant.

Scattering data were recorded simultaneously to fluorescence measurements for each dataset. For the analysis of protein states the first derivative of scattering data was used. syn-F/SBPase samples (final assay concentration of 150 ng/uL) were prepared as described above in absence and presence of 10 mM DTT and 0.5 mM GAP, respectively. The protein was pre-incubated with the respective buffers for different durations, as indicated. All pre-incubated samples were analyzed in the same run.

### Kinetic metabolic model

#### Model structure

The kinetic model for *Synechocystis* central carbon metabolism was based on a previous model [16]. The final model contained 29 reactions connecting 36 metabolites (22 internal). Sink reactions were formulated as irreversible Michaelis-Menten-type equations. Phosphate supply followed mass action kinetics. Two model variants were created: One base model, including only the regulatory interactions in the previous version [16], and one model with interactions on F/SBPase (GAP and NADPH).

#### Metabolite concentrations and flux distribution

Due to the uncertainty associated with published metabolomics datasets, potential thermodynamically feasible metabolite concentrations describing the metabolic state were randomly sampled as performed previously [16]. Metabolite concentration ranges identified via NET analysis [25] were used as constraints for the sampling, resulting in ∼3000 feasible metabolite concentration sets (fMCSs) covering the entire thermodynamically allowable solution space.

The steady-state flux distribution was obtained using a genome-scale metabolic model (GEM) of *Synechocystis* (Knoop et al., 2013) as described in Janasch et al., 2019 [85]. All flux simulations were performed in Matlab R2020b using the *Gurobi Optimizer* version 9.1.1. Maximizing autotrophic growth was set as the objective function. Fluxes were manually curated to adjust the genome-scale flux distribution to the small-scale kinetic model structure and transformed into mM/min by multiplying with a cellular density of 434.78 g/L (*E. coli*, [86]).

#### Parameter Sampling

Rate equations were generally parameterized around the corresponding metabolite concentrations by sampling the range of 0.1x to 10x metabolite concentration in logarithmic space for K_M_ values. Inhibition constants K_i_ and activation constant K_a_ for the regulations identified by LiP-SMap were sampled in a narrower range of 0.5x to 2x around the metabolite concentrations used for the enzyme assays. For the activation of F/SBPase by GAP, K_m_ could maximally be reduced by 75%. Hill coefficients for F/SBPase were sampled between 1 and 1.5. V_max_ values were calculated back from metabolite concentrations, sampled kinetic constants and the steady-state flux distribution. For each of the 3151 fMCSs, 1000 parameter sets were generated, resulting in an ensemble of ∼3 million kinetic steady-state models to be analyzed for stability and metabolic control.

#### Metabolic control analysis

The dynamic behavior of the models was analyzed by linearizing them around their steady-state as performed previously, by forming the Jacobian matrix y [16, 87] [88]. The stability of each model in the ensemble was evaluated by calculating the eigenvalues of the Jacobian matrix, where positive eigenvalues cause instability. Flux control coefficients were calculated for all stable parameter sets based on elasticities and concentration control coefficients as described previously[89]. The models and all code required to perform the kinetic modeling analysis is available at https://github.com/MJanasch/KX_Kinetics.

## Supporting information

Supplementary Information Lip-SMAP

## Abbreviations

2OG: 2-oxoglutarate
2PG: 2-phosphoglycolate
3PGA: 3-phosphoglycerate
6PG: 6-phosphogluconate
AcCoA: Acetyl-CoA
ADP: Adenosine diphosphate
AGPase: ADP-glucose synthase (EC 2.7.7.27)
AMP: Adenosine monophosphate
ATP: Adenosine triphosphate
cAMP: Cyclic adenosine monophosphate
Cit: Citrate
DAHPS: 3-Deoxy-D-arabinoheptulosonate 7-phosphate synthase (EC 2.5.1.54)
dPGM: 2,3-diphosphoglycerate-dependent phosphoglycerate mutase (EC 5.4.2.11)
ED: Entner–Doudoroff pathway
EMP: Embden-Meyerhof-Parnas pathway
ENO: Enolase (EC 4.2.1.11)
FBA: Fructose-bisphosphate aldolase (EC 4.1.2.13)
FBP: Fructose-1,6-bisphosphate
FBPase: Fructose-1,6-bisphosphatase (EC 3.1.3.11)
F/SBPase: Bifunctional fructose-1,6/sedoheptulose-1,7-bisphosphatase (EC 3.1.3.11)
G6P: Glucose-6-phosphate
G6PDH: Glucose-6-phosphate dehydrogenase (EC 1.1.1.49)
GAP: Glyceraldehyde-3-phosphate
GAPDH: Glyceraldehyde-3-phosphate dehydrogenase (EC 1.2.1.12, 1.2.1.13, 1.2.1.59)
Glyc: Glycolate
Glyx: Glyoxylate
GPM: Phosphoglucomutase (EC 5.4.2.2)
GTP: Guanosine triphosphate
iPGM: 2,3-diphosphoglycerate-independent phosphoglycerate mutase (EC 5.4.2.12)
KDPG: 2-dehydro-3-deoxy-D-gluconate-6-phosphate
Mal: Malate
NADP: Nicotinamide adenine dinucleotide phosphate (oxidized)
NADPH: Nicotinamide adenine dinucleotide phosphate (reduced)
PEP: Phosphoenolpyruvate
PGI: Phosphoglucoisomerase (EC 5.3.1.9)
PGK: Phosphoglycerate kinase (EC 2.7.2.3)
Phe: Phenylalanine
PP: Pentose Phosphate pathway
PRK: Phosphoribulokinase (EC 2.7.1.19)
PYK: Pyruvate kinase (EC 2.7.1.40)
RPE: Ribulose-phosphate 3-epimerase (EC 5.1.3.1)
RPI: Ribose-5-phosphate isomerase (EC 5.3.1.6)
RPPK: Ribose-5-phosphate pyrophosphokinase (EC 2.7.6.1)
Ru5P: Ribulose–5-phosphate
Rubisco: Ribulose-bisphosphate carboxylase (EC 4.1.1.39)
RuBP: Ribulose-1,5-bisphosphate
SBPase: Sedoheptulose-1,7-bisphosphatase (EC 3.1.3.37)
SerA: Phosphoglycerate dehydrogenase (EC 1.1.1.95)
Suc: Sucrose
TAL: Transaldolase (EC 2.2.1.2)
TKT: Transketolase (EC 2.2.1.1)
TPI: Triose-phosphate isomerase (EC 5.3.1.1)

## Acknowledgements

We are grateful to Michael Jahn (KTH Stockholm) for helpful discussion on proteomics and cultivation of *Cupriavidus necator*. We thank Ralf Steuer (Humboldt University, Berlin) for discussions on kinetic modeling.

Funding for this work was from the Novo Nordisk Foundation (grant numbers NNF19OC0057652 and NNF20OC0061469), the Swedish Research Council Vetenskapsrådet (grant number 2016-06160), and the Swedish Foundation for Strategic Research SSF (ARC19-0051).

## Contributions

Conceptualization: JK, ES, PS, FE, IP, EPH

Experimental proteomics: JK, ES, DK, LS, FE

Proteomics data analysis: JAS, JK, ES, LS, FE

Enzyme kinetics and melting analyses: JK, KS, ES

Metabolic modeling: MJ, LZ

Writing initial draft: JK, ES, JAS, MJ, KS, EPH

Editing of final draft: JK, ES, JAS, MJ, KS, EPH

Funding acquisition: PS, EPH

