## Supplementary Information Lip-SMAP for "Metabolite interactions in the bacterial Calvin cycle and implications for flux regulation"

**List of Supplementary Figures and Tables**

**Figure S1.** Change in LiP digestion upon reduction and oxidation of *Synechocystis* proteome
**Figure S2:** Number of detected peptides in every LiP-SMap experiment
**Figure S3:** Number of peptides detected per metabolite-interacting protein compared to non-interacting proteins
**Figure S4:** Overlap and correlation of peptides from repeat LiP-SMap experiments
**Figure S5:** Persistence of interactions at low metabolite concentrations at high metabolite concentrations
**Figure S6:** Fraction interacting orthologs within functional groups in each organism
**Figure S7:** Similarity of ortholog interaction patterns (low concentration)
**Figure S8:** Log2(fold change) and significance of detected proteins in presence of 2 mM ATP at different $Mg^{2+}$ concentrations
**Figure S9:** Interactions of Calvin cycle enzymes and selected central carbon metabolism enzymes with metabolites (low concentration)
**Figure S10:** Schematic overview of the Calvin cycle
**Figure S11:** Effect of DHAP and GAP on the catalytic activity of syn-F/SBPase
**Figure S12:** Glyceraldehyde-3-phosphate (GAP) effect on thermal stability of *Synechocystis* F/SBPase at different $Mg^{2+}$ concentrations
**Figure S13:** Thermostability assays of *Cupriavidus* F/SBPase in the presence of various metabolites
**Figure S14:** Thermostability assays of *Synechocystis* F/SBPase in the presence of various metabolites
**Figure S15:** End-point *in vitro* assay of cnF/SBPase in presence of G6P
**Figure S16:** Light scattering assays of syn-F/SBPase under various conditions
**Figure S17:** Measured concentrations of product after 20 minutes of *in vitro* reaction as detected by malachite green assay and LC/MS
**Figure S18:** Effect of different metabolites (1 mM) on the kinetics of *Cupriavidus* transketolase
**Figure S19:** Effect of different metabolites (1 mM) on the kinetics of *Synechocystis* transketolase
**Figure S20:** Thermostability assays of *Synechocystis* and *Cupriavidus* transketolase in the presence of various metabolites
**Figure S21:** Kinetic analysis of the *Synechocystis* F/SBPase R194H mutant
**Figure S22:** Flux control coefficients for all reactions in the model
**Figure S23:** Difference between median FCCs between model variants

**Table S1:** Chosen concentrations for every used metabolite and their most extreme values found in literature
**Table S2:** All metabolite concentrations found across 7 metabolomics studies

**Table S3:** (separate file) Changes in fructose/sedoheptulose bisphosphatase kinetic parameters in the presence of metabolites
**Table S4:** Changes in transketolase kinetic parameters in the presence of various metabolites at 1 mM
**Table S5:** Transition list for mass spectrometry

**Supplemental Dataset S1** (separate data file) List of significantly changed proteins by reduction/oxidation through DTT/DTNB
**Supplemental Dataset S2:** (separate data file) List of all detected peptides across all LiP-SMap experiments
**Supplemental Dataset S3:** (separate data file) List of all proteins affected by at least one metabolite and their KEGG orthology groups (KOGs)
**Supplemental Dataset S4:** (separate data file) Phylogenetic trees of Calvin cycle enzymes labeled with detected protein-metabolite interactions

Supplemental datasets S1-S4 are available at
https://drive.google.com/drive/folders/1MJrtCzXYD-0mmGHh6Wt9lwP-hzko2zOh

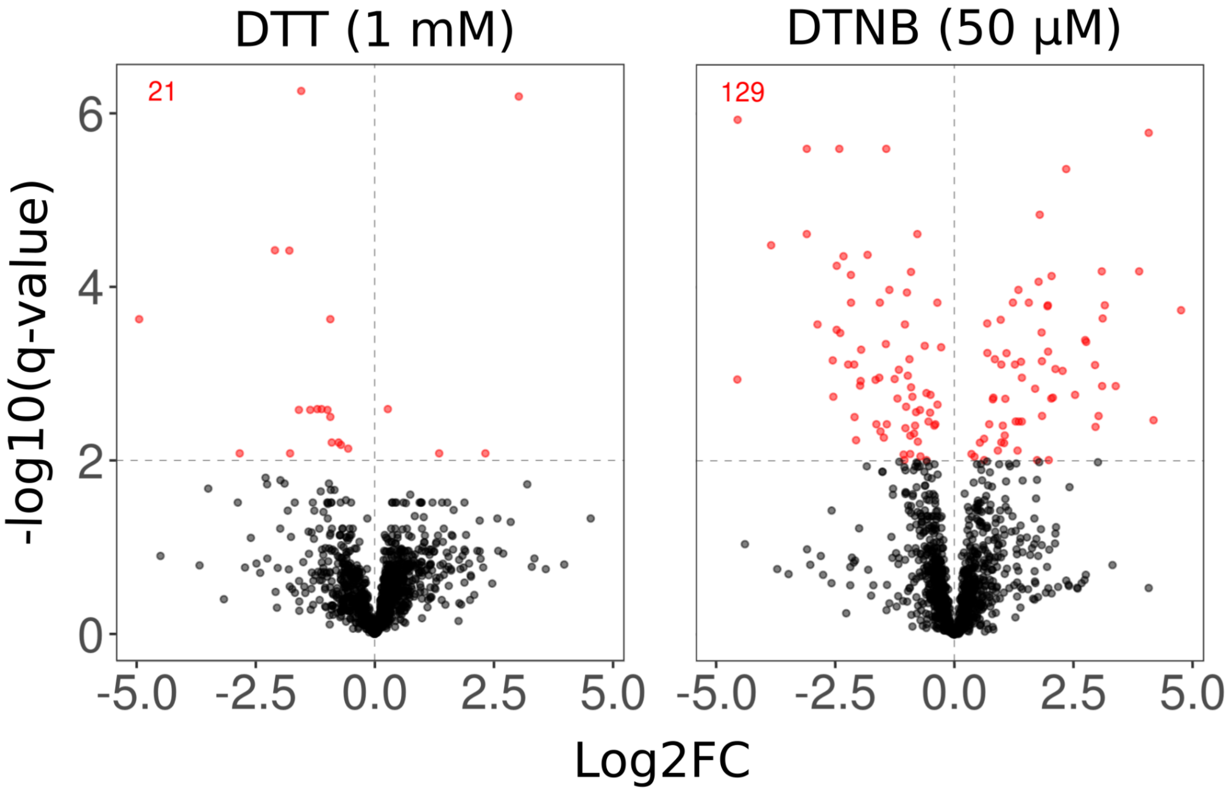


**Figure S1. Change in LiP digestion upon reduction and oxidation of *Synechocystis* proteome.** Synechocystis proteome extracts treated with either DTT (reductant) or DTNB (oxidant) were subjected to LiP-SMap analysis (untreated proteome as control). Each point represents a protein that was detected in both the treated and untreated condition. A point's coordinate on the x-axis corresponds to the peptide abundance fold change (FC), treated vs. untreated, of the peptide with the lowest q-value in that particular protein. Y-axes show the statistical significance of the fold change. Red points indicate significantly changed proteins ($q < 0.01$), and insets display the number of significantly changed proteins.

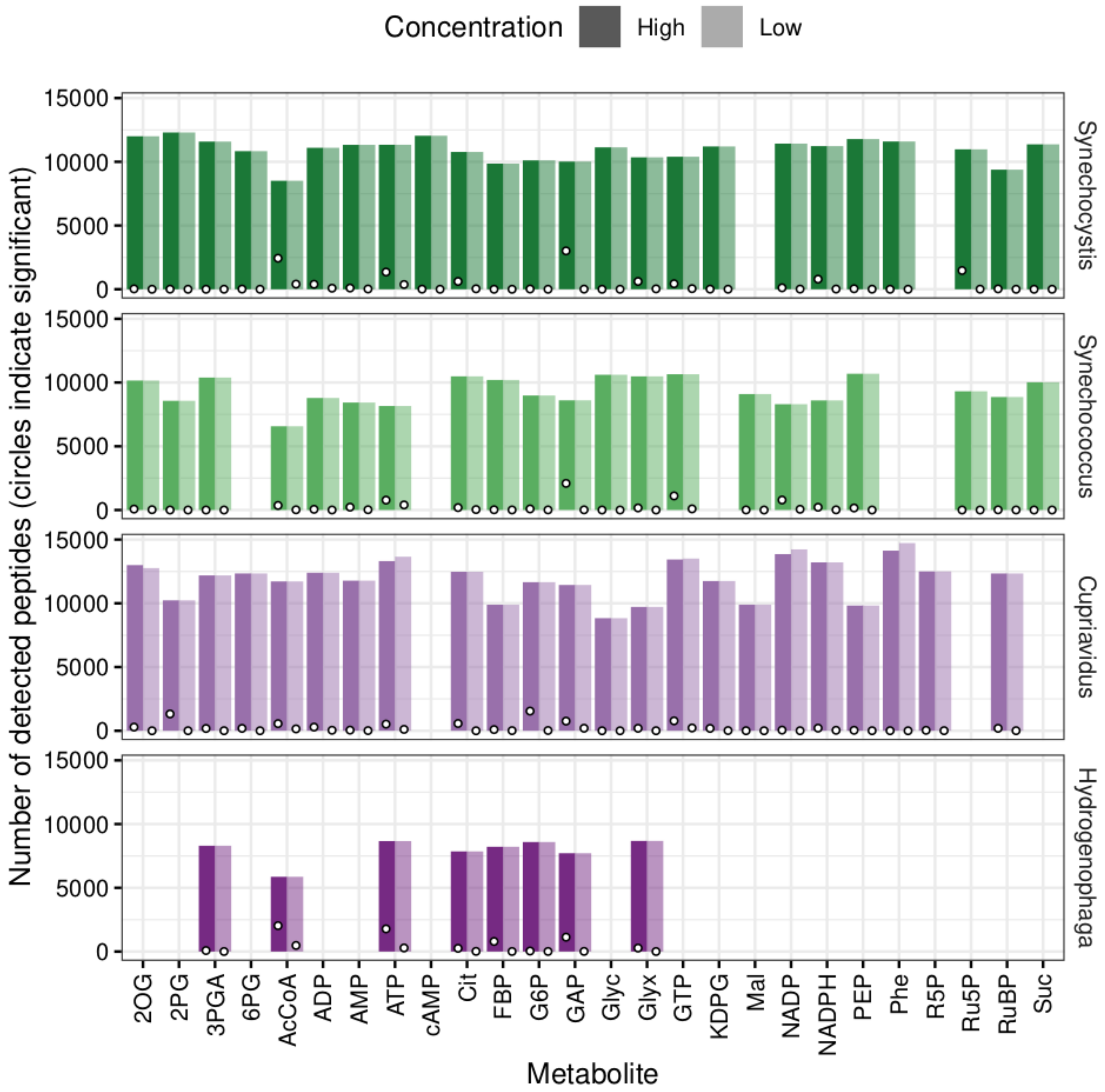


**Figure S2. Number of detected peptides in every LiP-SMap experiment.** The darker hue represents the high concentration of added metabolite and the lighter hue represents the low concentration. Circles indicate the number of significant peptides. When a metabolite was not tested against an organism, the space is blank.

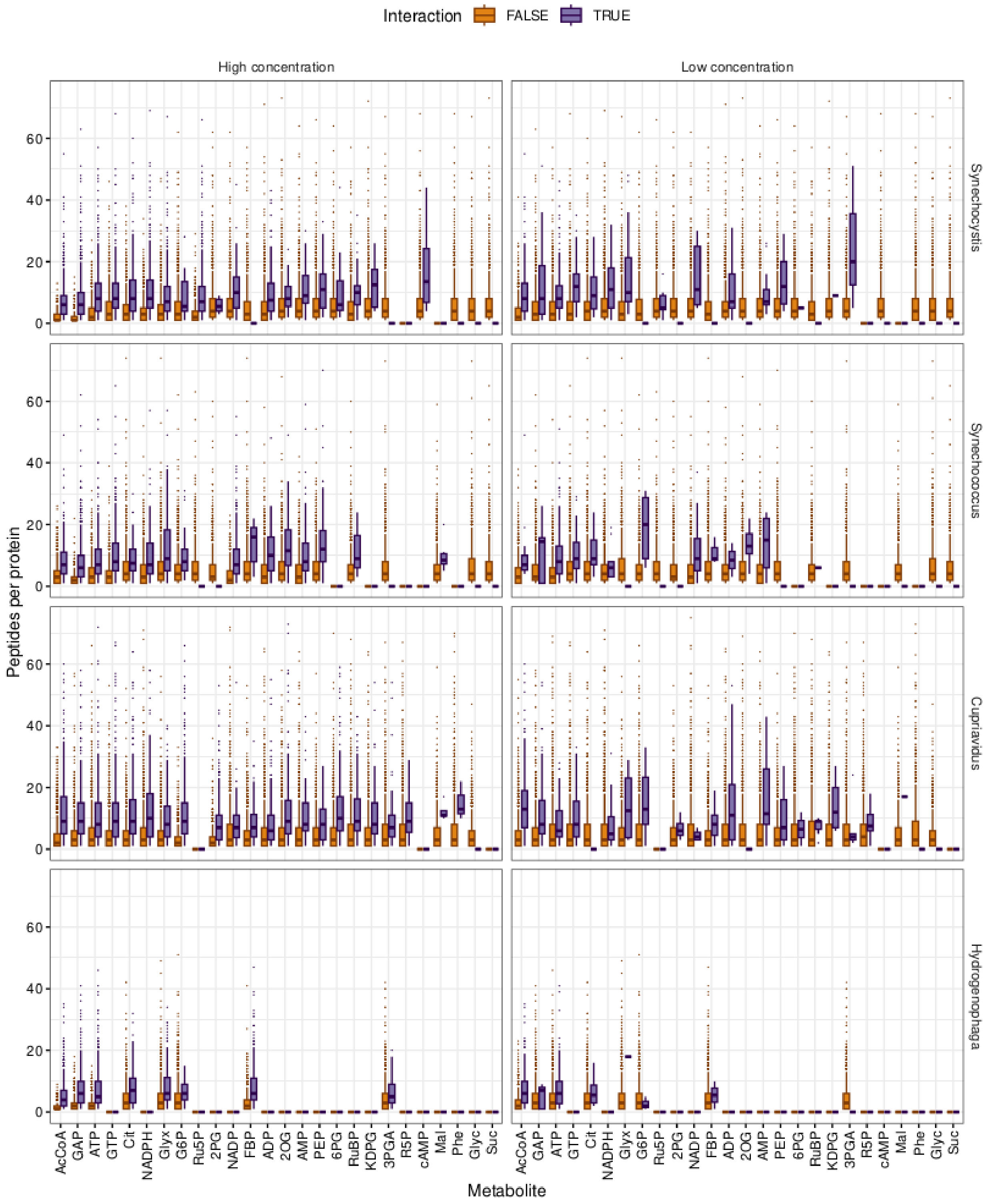


**Figure S3. Number of peptides detected per metabolite-interacting protein compared to non-interacting proteins.** Proteins were classified as not having interaction or having interaction with the tested metabolites, *i.e.* “Interaction FALSE” and “Interaction TRUE”. Interaction means that at least one peptide was significantly changed in abundance in presence of the metabolite ($q$ < 0.01). The y-axis indicates the number of peptides detected per protein summarized as box plots. The plots are split by high and low concentration of the interacting metabolite (columns) and by organism (rows).

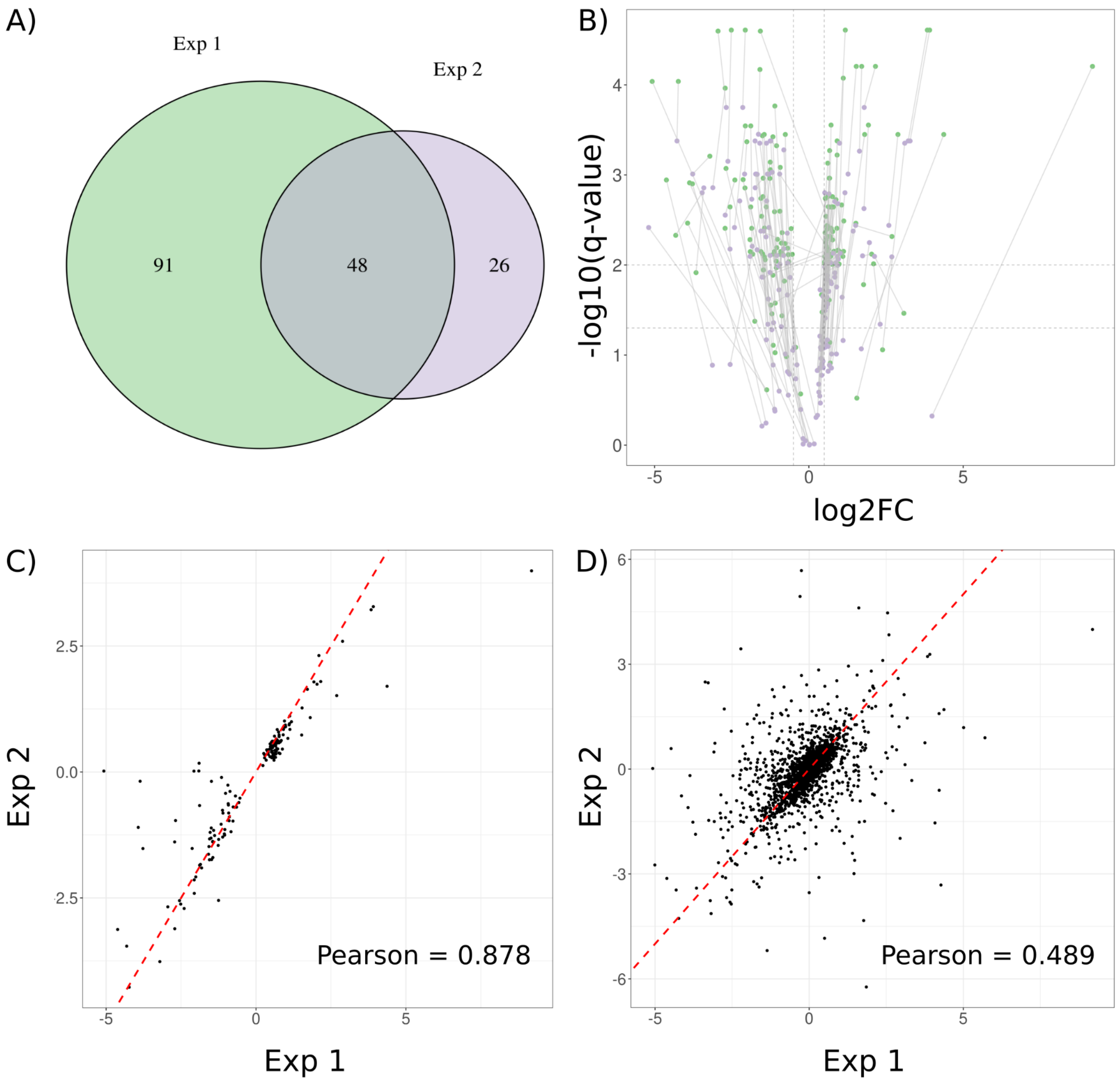


**Figure S4. Overlap and correlation of peptides from repeat *Synechococcus* PCC 7942 LiP-SMap experiments with 10 mM glyoxylate.** Two LiP-SMap experiments were performed in parallel, with the same reagent stocks and were run together on the LC-MS. A) Overlap of significant peptides between the two experiments. B) Volcano plot of all peptides significant in at least one experiment, with each peptide pair connected by a line. The lines being generally vertical indicate that the experiments differ mainly in significance rather than effect size. C) Correlation of the log2 fold changes of peptides with a q-value < 0.01 upon treatment with 10 mM glyoxylate D) Correlation of the log2 fold changes all peptides upon treatment with 10 mM glyoxylate

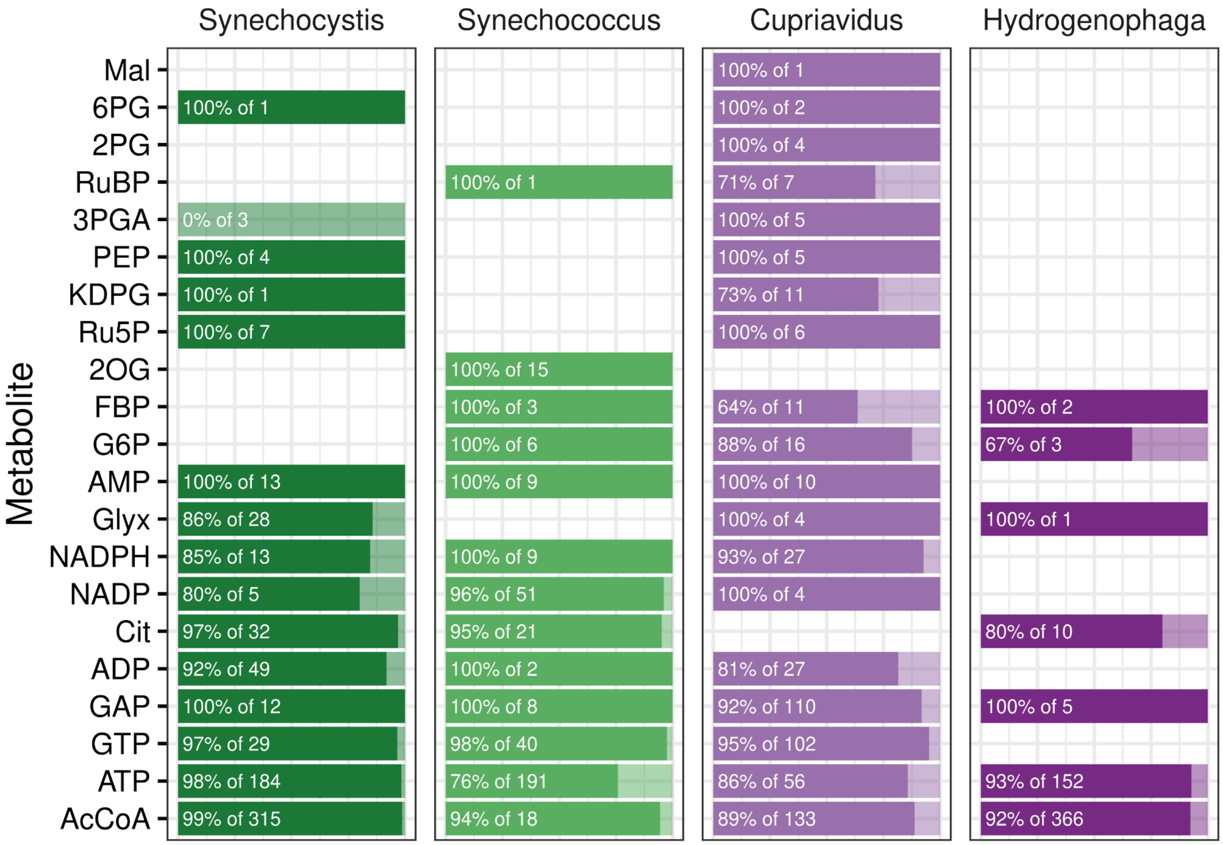


**Figure S5. Persistence of interactions at low metabolite concentrations at high metabolite concentrations.** Opaque bars indicate the fraction of low concentration interactions that were detected both in the low concentration and high concentration experiments, while transparent bars indicate interactions that were only detected in the low concentration experiments. Metabolites are ordered by the total number of interactions. Metabolites without low concentration interactions are excluded.

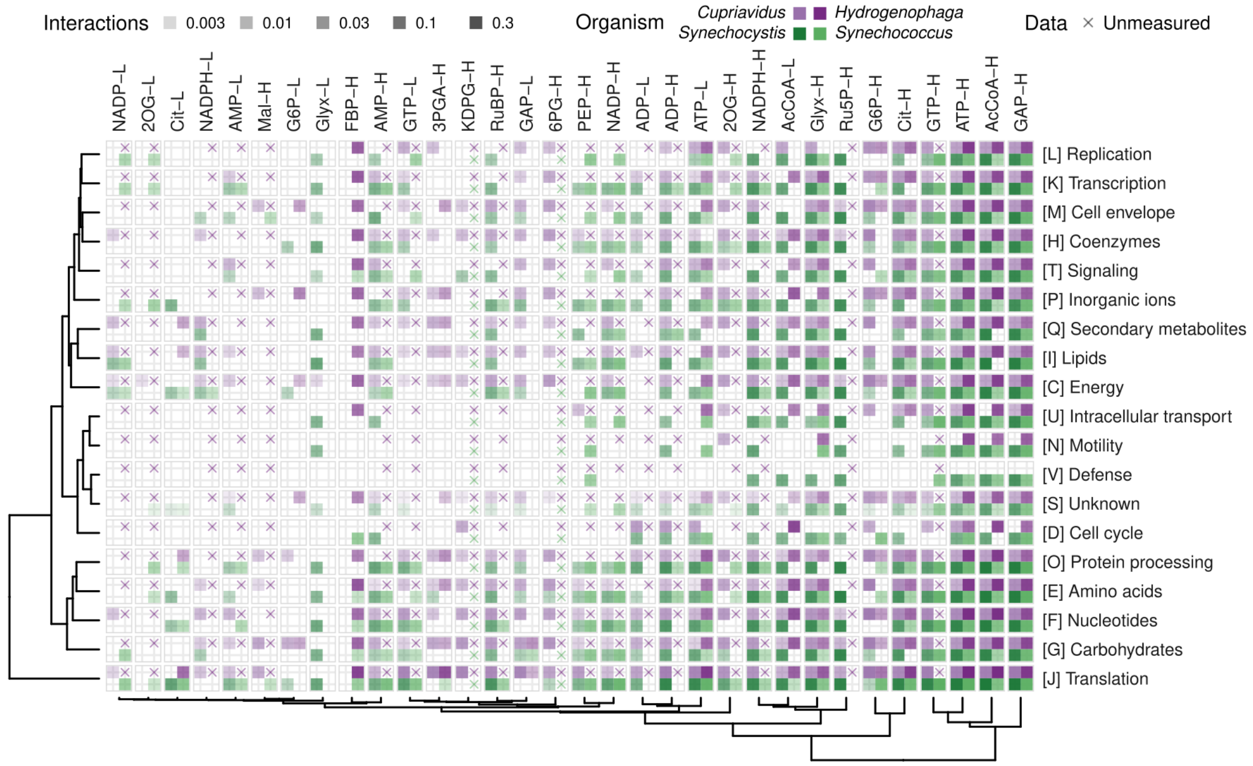


**Figure S6. Fraction interacting orthologs within functional groups in each organism.** If at least one sequence per ortholog family interacted with a metabolite at low (L) or high (H) concentration (heatmap columns), that ortholog was considered to be interacting. Interactions were then summarized per functional group (heatmap rows) and normalized by the total number of orthologs in that group. Dendrograms illustrate the clustering patterns of rows and columns based on Euclidean distance and the Ward.D2 algorithm. Interaction fractions in all four organisms contributed both to rows and columns. A cross indicates that the particular condition was not measured.

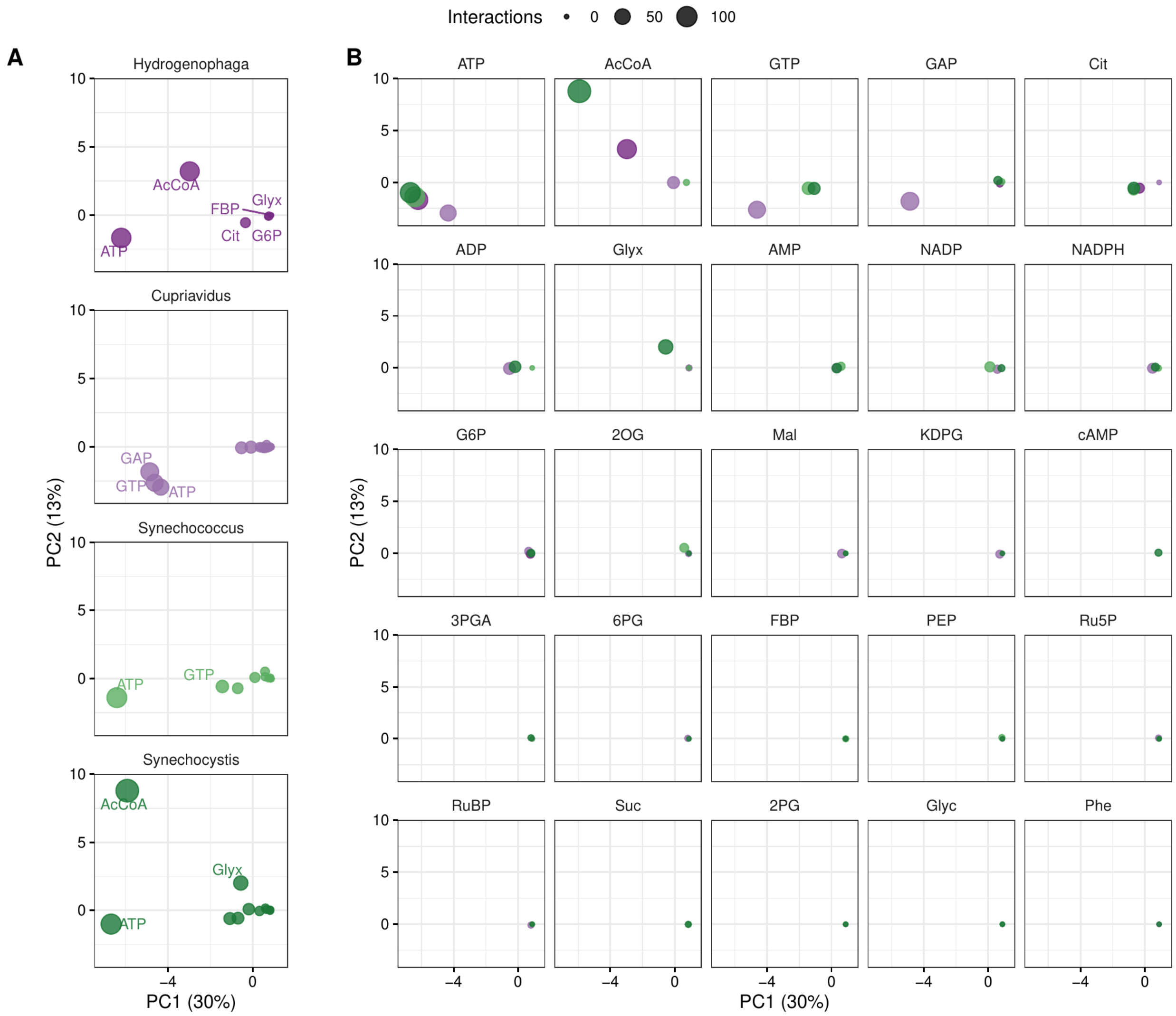


**Figure S7. Similarity of ortholog interaction patterns (low concentration).** Principal components were calculated from the presence or absence of interaction with each of 321 orthologs (see Materials and Methods). All data points shown here are from the same principal component analysis, but split per organism (A) or metabolite (B) to reduce overplotting. Percentages indicate the fraction of the total variance captured by the principal components.

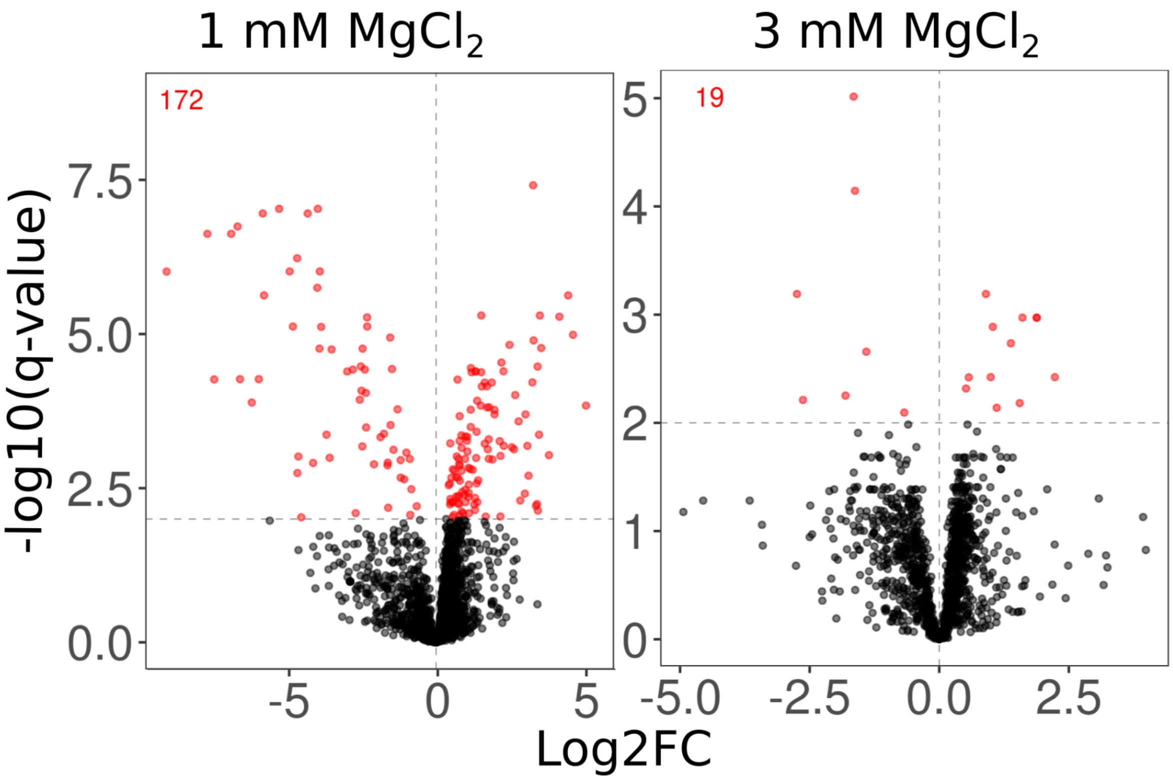


**Figure S8. Log2(fold change) and significance of detected proteins in presence of 2 mM ATP at different $Mg^{2+}$ concentrations.** The extracted proteome of *Synechocystis* was treated with 2 mM ATP and either 1 or 3 mM $MgCl_2$ and compared to a sample without ATP but with the same concentration $MgCl_2$. Each protein detected in both treated and untreated samples are represented by one dot with significantly ($q < 0.01$) changed proteins colored in red. The effect of ATP treatment is mitigated by an increased $MgCl_2$ concentration.

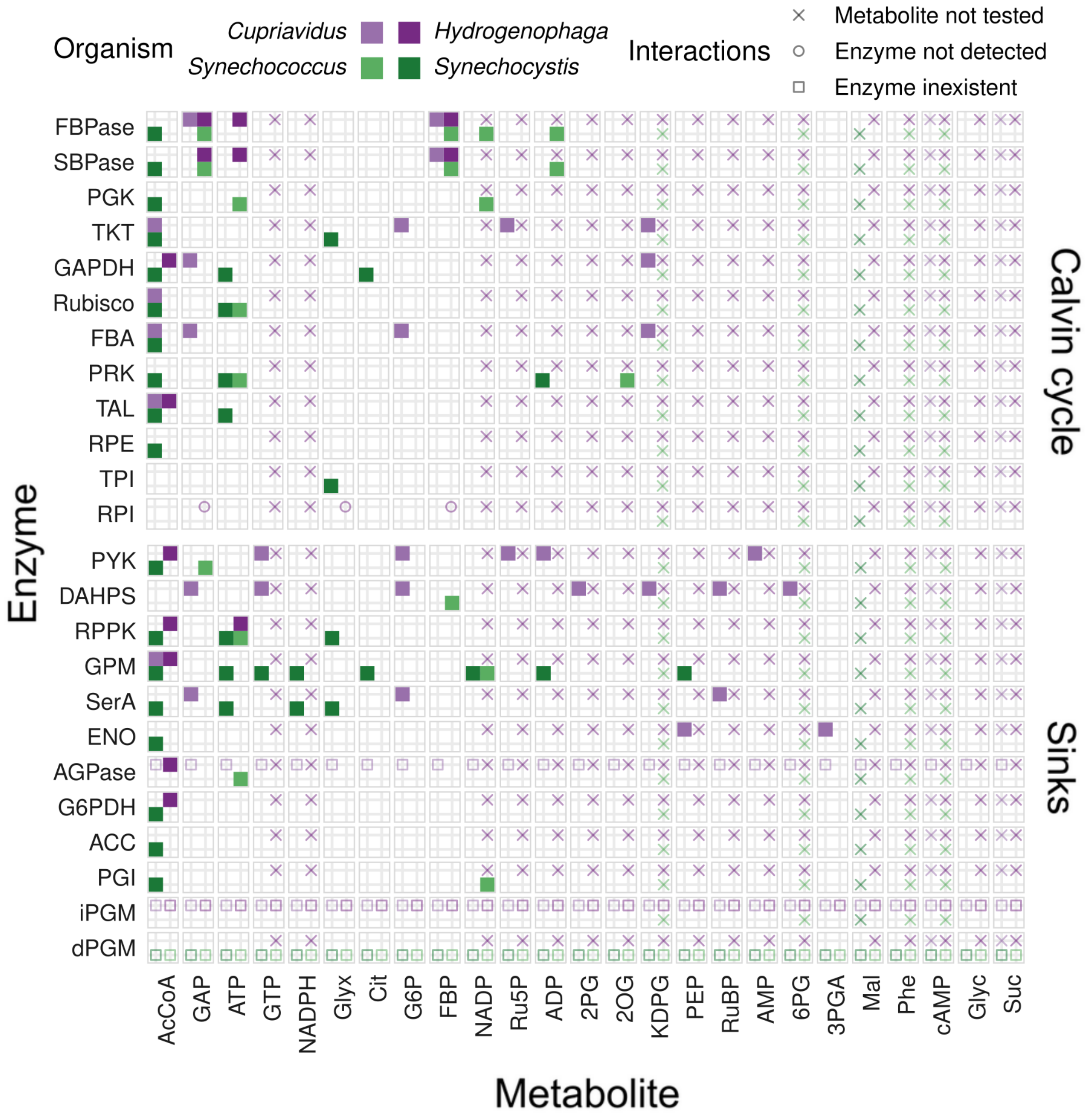


**Figure S9. Interactions of Calvin cycle enzymes and selected central carbon metabolism enzymes with metabolites (low concentration).** Interactions between metabolites (columns) at low concentration and enzymes (rows) identified by KEGG EC number annotation are shown for each organism by tiles filled with the corresponding color. A blank tile indicates that the interaction was not detected, while missing protein data is explained by a symbol. A cross indicates that the particular condition was not measured, a circle indicates that no proteins were detected, and a square indicates that there was no such enzyme in the corresponding genome. AGPase, ADP-glucose synthase (EC 2.7.7.27); DAHPS, DAHP synthase (EC 2.5.1.54); dPGM, 2,3-diphosphoglycerate-dependent phosphoglycerate mutase (EC 5.4.2.11); ENO, Enolase (EC 4.2.1.11); FBA, Fructose-bisphosphate aldolase (EC 4.1.2.13); FBPase, Fructose-1,6-bisphosphatase (EC 3.1.3.11); G6PDH, Zwf (EC 1.1.1.49); GAPDH, Glyceraldehyde 3-phosphate dehydrogenase (EC 1.2.1.12, 1.2.1.13, 1.2.1.59); GPM, Phosphoglucomutase (EC 5.4.2.2); iPGM, 2,3-diphosphoglycerate-independent phosphoglycerate mutase (EC 5.4.2.12); PGI,

Phosphoglucoisomerase (EC 5.3.1.9); PGK, Phosphoglycerate kinase (EC 2.7.2.3); PRK, Phosphoribulokinase (EC 2.7.1.19); PYK, Pyruvate kinase (EC 2.7.1.40); RPE, Ribulose-phosphate 3-epimerase (EC 5.1.3.1); RPI, Ribose 5-phosphate isomerase (EC 5.3.1.6); RPPK, Ribose-5-phosphate pyrophosphokinase (EC 2.7.6.1); Rubisco, Ribulose-bisphosphate carboxylase (EC 4.1.1.39); SBPase, Sedoheptulose-1,7-bisphosphatase (EC 3.1.3.37); SerA, Phosphoglycerate dehydrogenase (EC 1.1.1.95); TAL, Transaldolase (EC 2.2.1.2); TKT, Transketolase (EC 2.2.1.1); TPI, Triose-phosphate isomerase (EC 5.3.1.1).

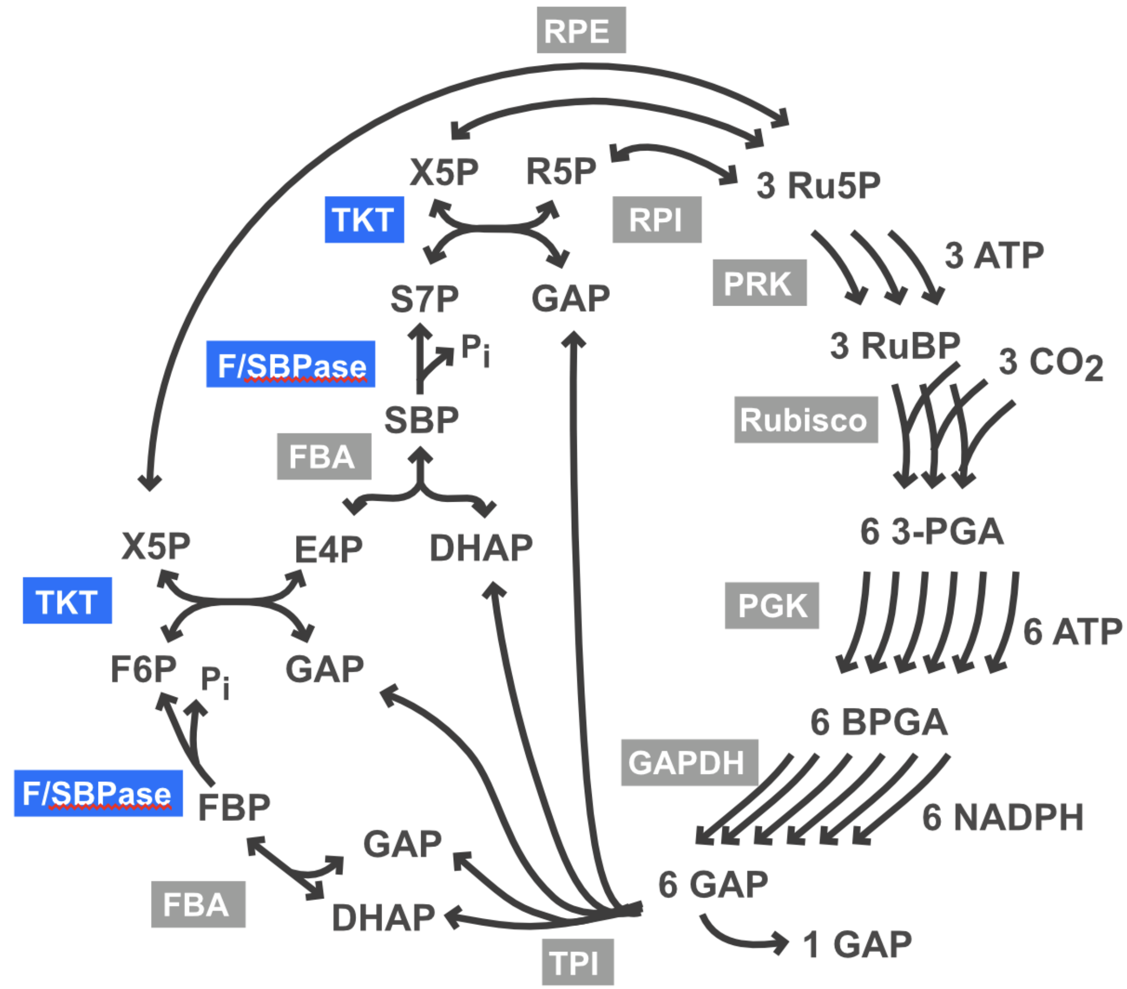


**Figure S10. Schematic overview of the Calvin cycle.** Enzymes that were characterized *in vitro* are outlined in blue.

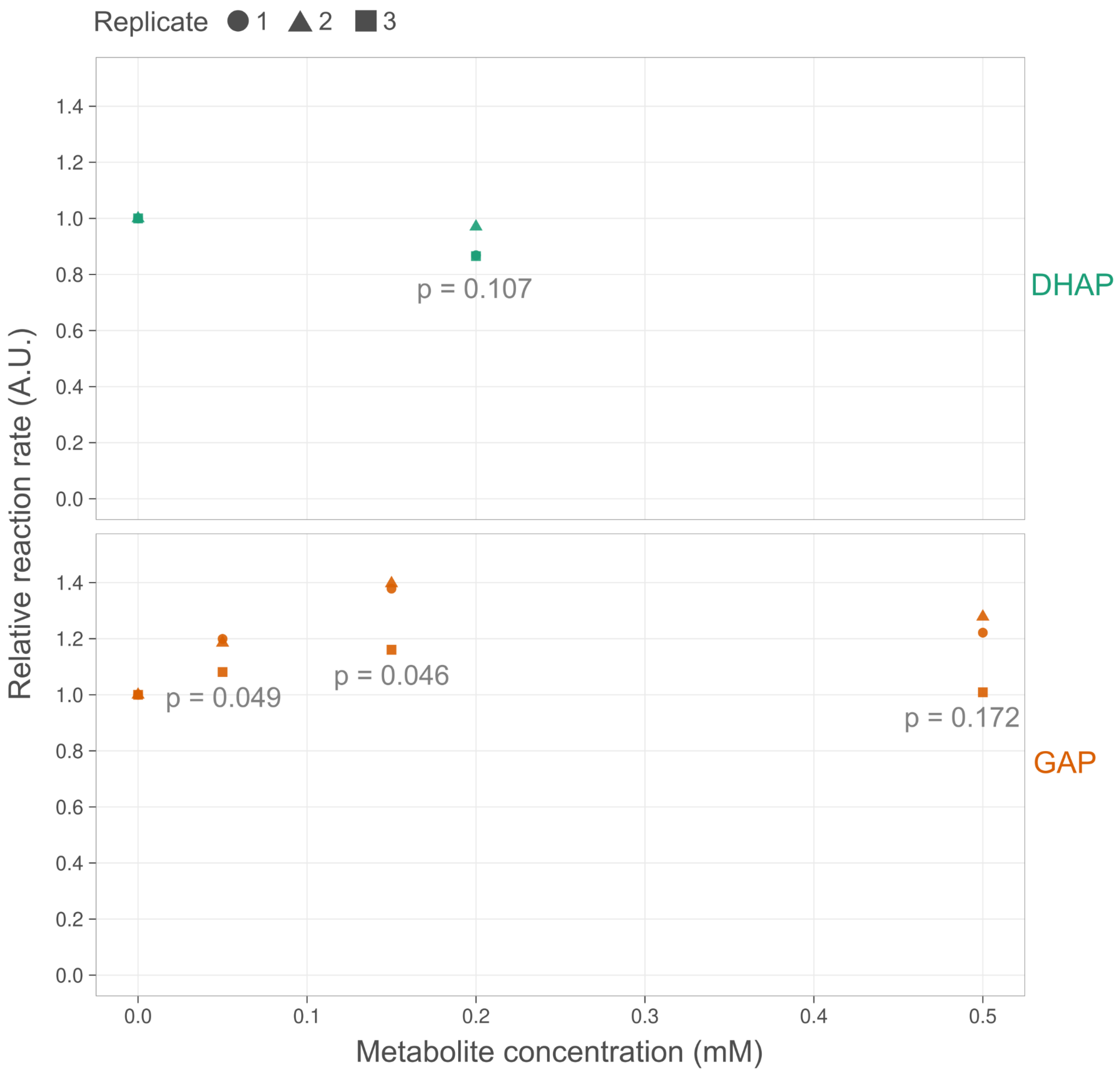


**Figure S11: Effect of DHAP and GAP on the catalytic activity of syn-F/SBPase.** The catalytic rate of *Synechocystis* F/SBPase (y-axis) was measured *in vitro* at different DHAP and GAP concentrations (x-axis) under half-saturating substrate concentration (60 µM FBP). Shown initial rates are relative to the rate measured in absence of metabolite e. P-values (p) show the statistical significance of the change in catalytic rate at each tested metabolite concentration relative to the rate at 0 µM metabolite (Student's t-test).

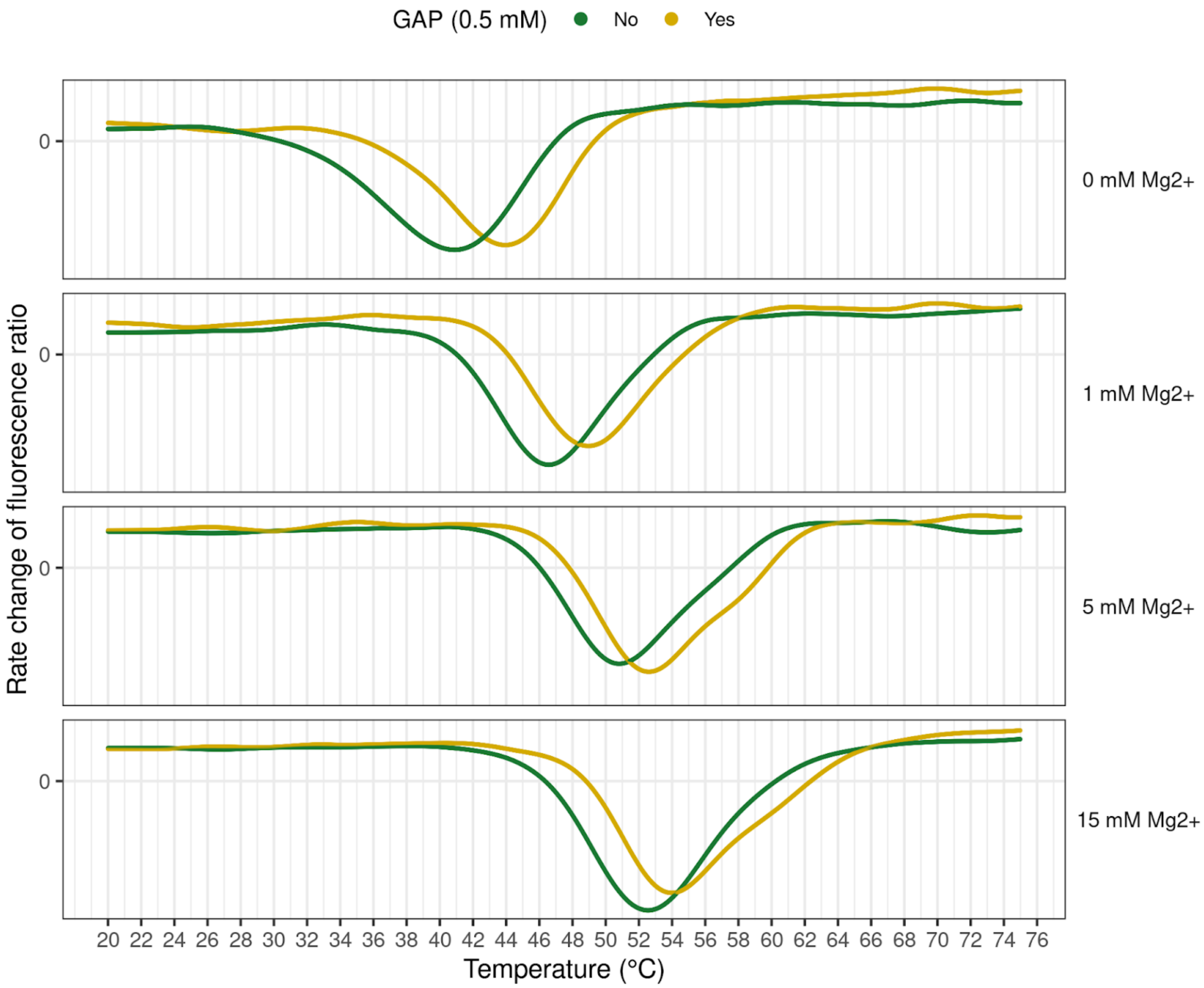


**Figure S12. Glyceraldehyde-3-phosphate (GAP) effect on thermal stability of *Synechocystis* F/SBPase at different $Mg^{2+}$ concentrations.** Curves indicate denaturation of F/SBPase over a temperature gradient of 1 °C/min. The Y-axis shows the change in the ratio of protein autofluorescence (350 nm/330 nm), and the temperature at minimum values indicate the melting temperature ($T_m$) at which half of the enzyme population is denatured.

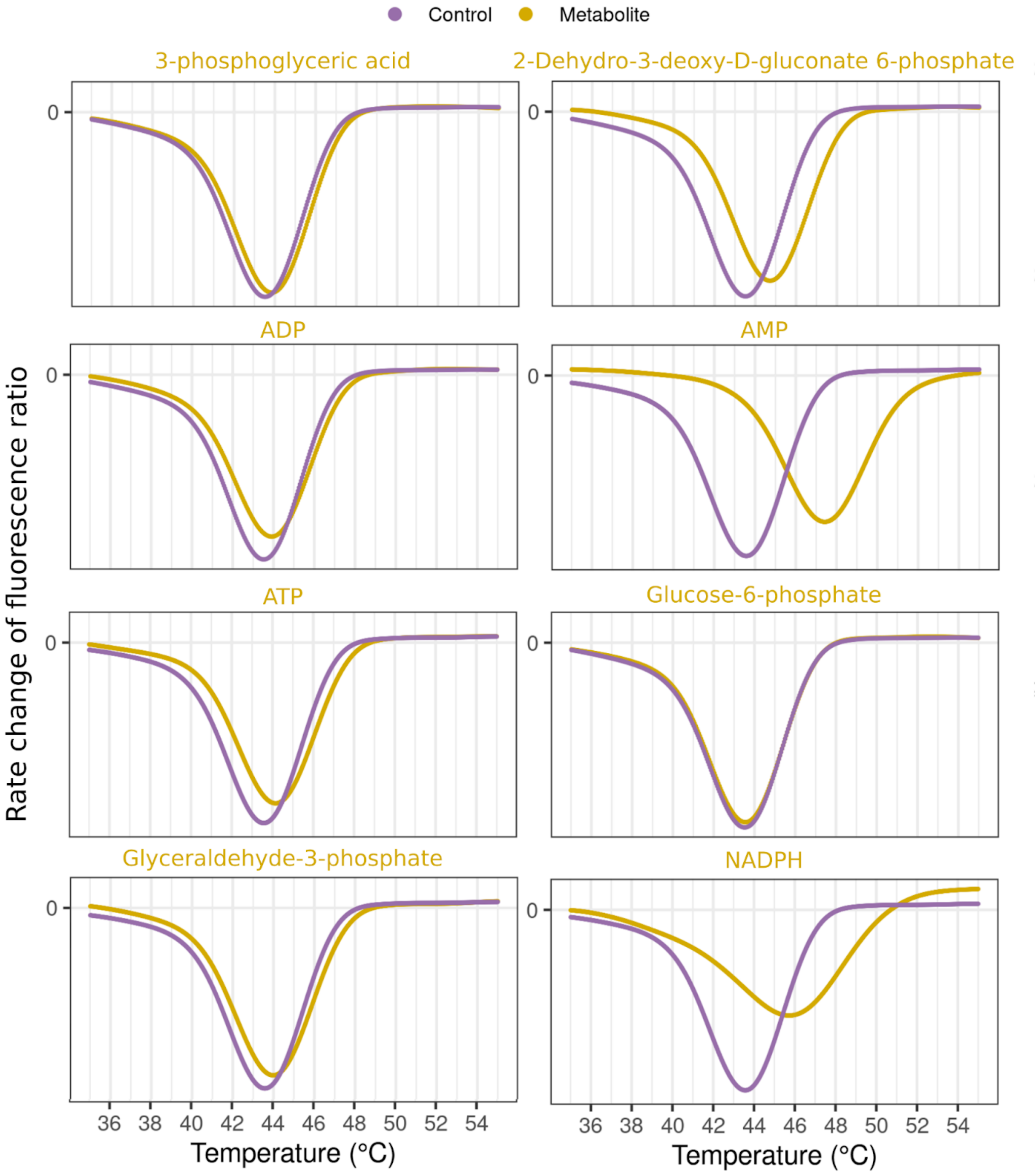


**Figure S13. Thermal shift assays of *Cupriavidus* F/SBPase in the presence of various metabolites.** All metabolites were tested at a concentration of 1 mM. Curves indicate denaturation of F/SBPase over a temperature gradient of 1 °C/min. Y-axis shows the change in the ratio of protein autofluorescence (350 nm/330 nm), and minimum values indicate the melting temperature ($T_m$) at which half of the enzymes are denatured.

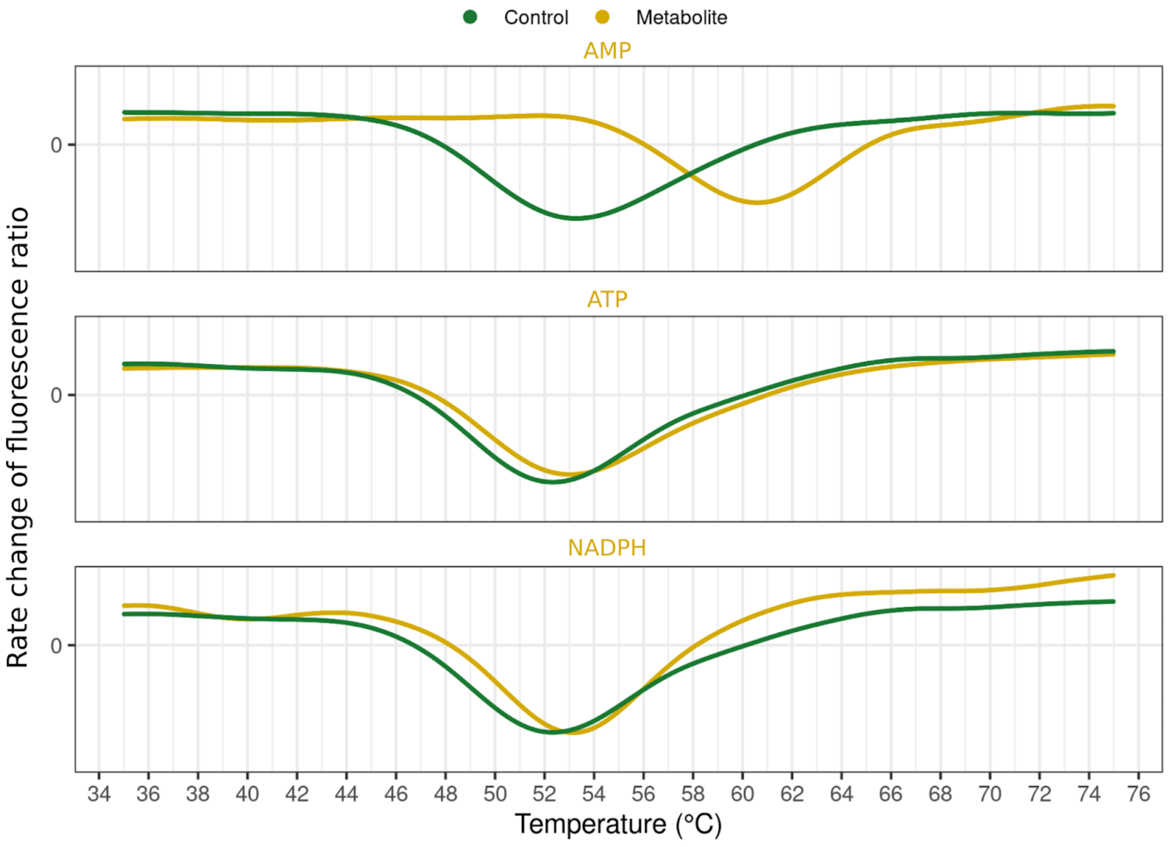


**Figure S14. Thermal shift assays of *Synechocystis* F/SBPase in the presence of various metabolites.** All metabolites were tested at a concentration of 1 mM. Curves indicate denaturation of F/SBPase over a temperature gradient of 1 °C/min. Y-axis shows the change in the ratio of protein autofluorescence (350 nm/330 nm), and minimum values indicate the melting temperature ($T_m$) at which half of the enzymes are denatured.

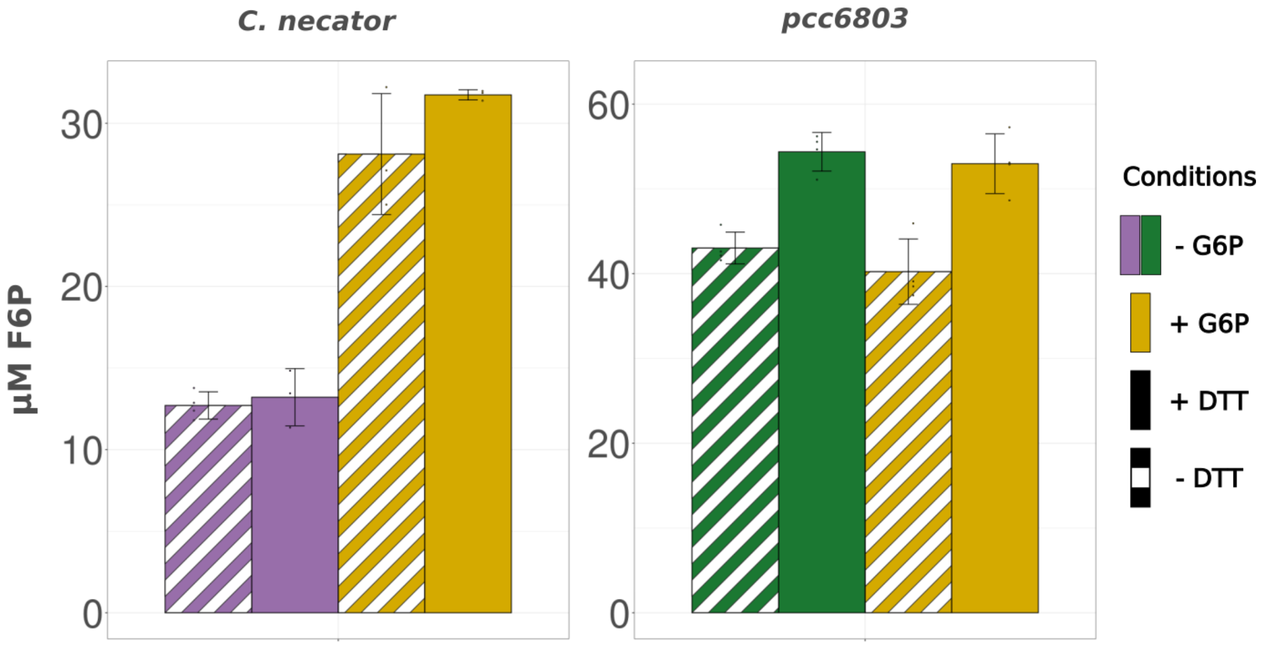


**Figure S15. End-point *in vitro* assay of F/SBPase in presence of G6P.** A substrate concentration of 150 µM was used and the enzyme concentration was 0.153 ng/µL for cnF/SBPase and 459 ng/µL for synF/SBPase. The method of detection was a malachite green assay after 20 minutes of reaction.

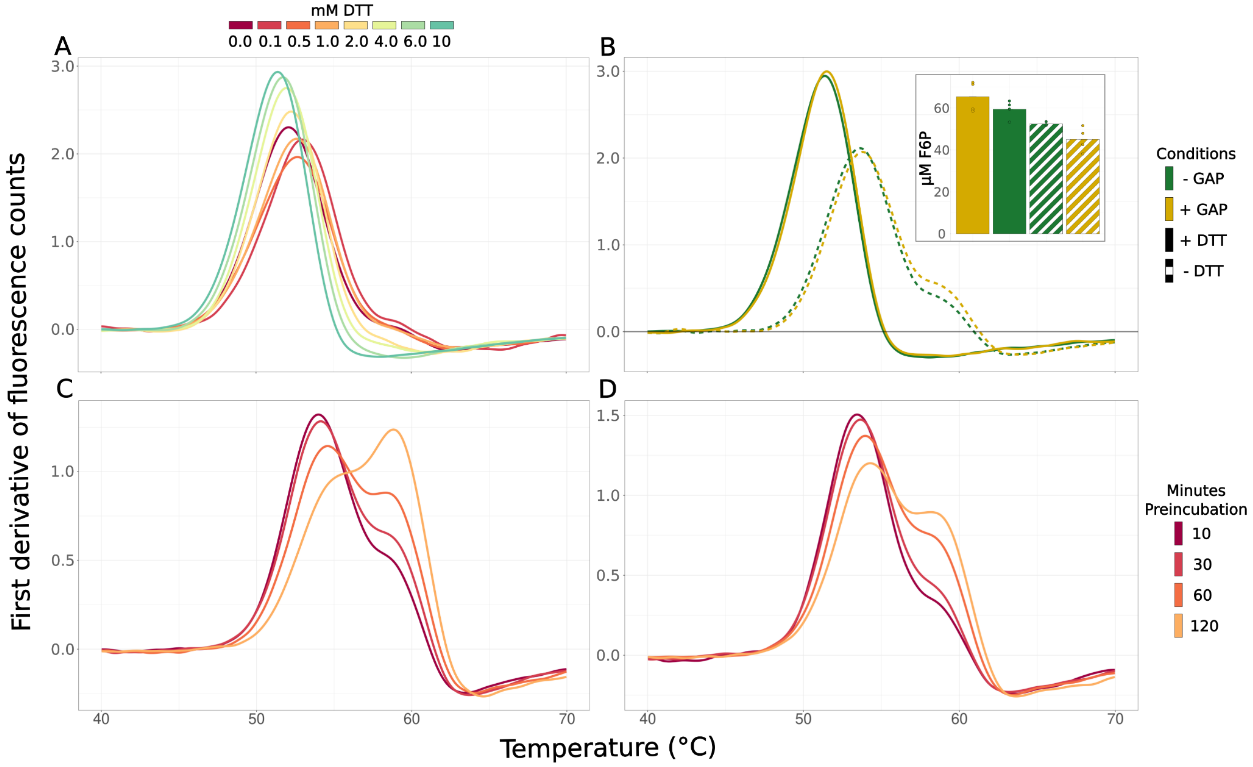


**Figure S16. Light scattering assays of syn-F/SBPase under various conditions. A)** Light scattering data of the enzyme at different concentrations of DTT ranging from 0 to 10 mM. **B)** Light scattering data for enzyme ± 10 mM DTT and ± 0.5 mM GAP. Also shown is the concentration of product measured by malachite green assay after 20 minutes of reaction as described under Methods **C)** Light scattering data of enzyme after different preincubation times at 30 degrees in the presence of 0.5 mM GAP. **D)** Light scattering data of enzyme after different preincubation times at 30 degrees in the absence of GAP.

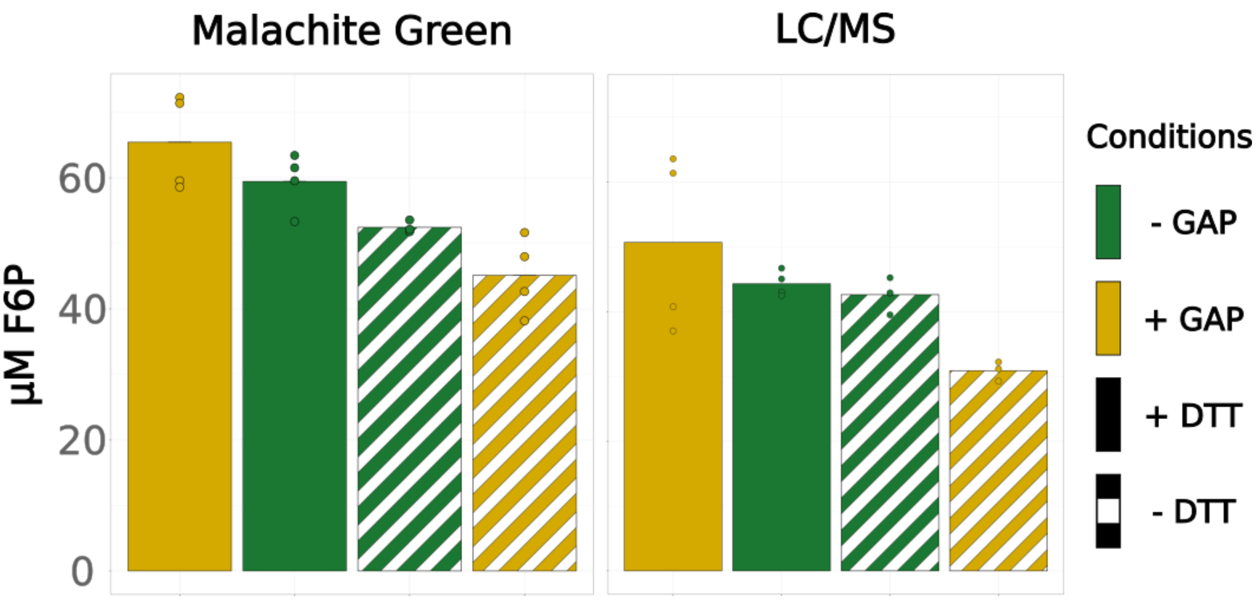


**Figure S17. Measured concentrations of product after 20 minutes of *in vitro* reaction as detected by malachite green assay and LC/MS.** A substrate concentration of 80 µM was used and the enzyme concentration was 0.23 ng/µL.

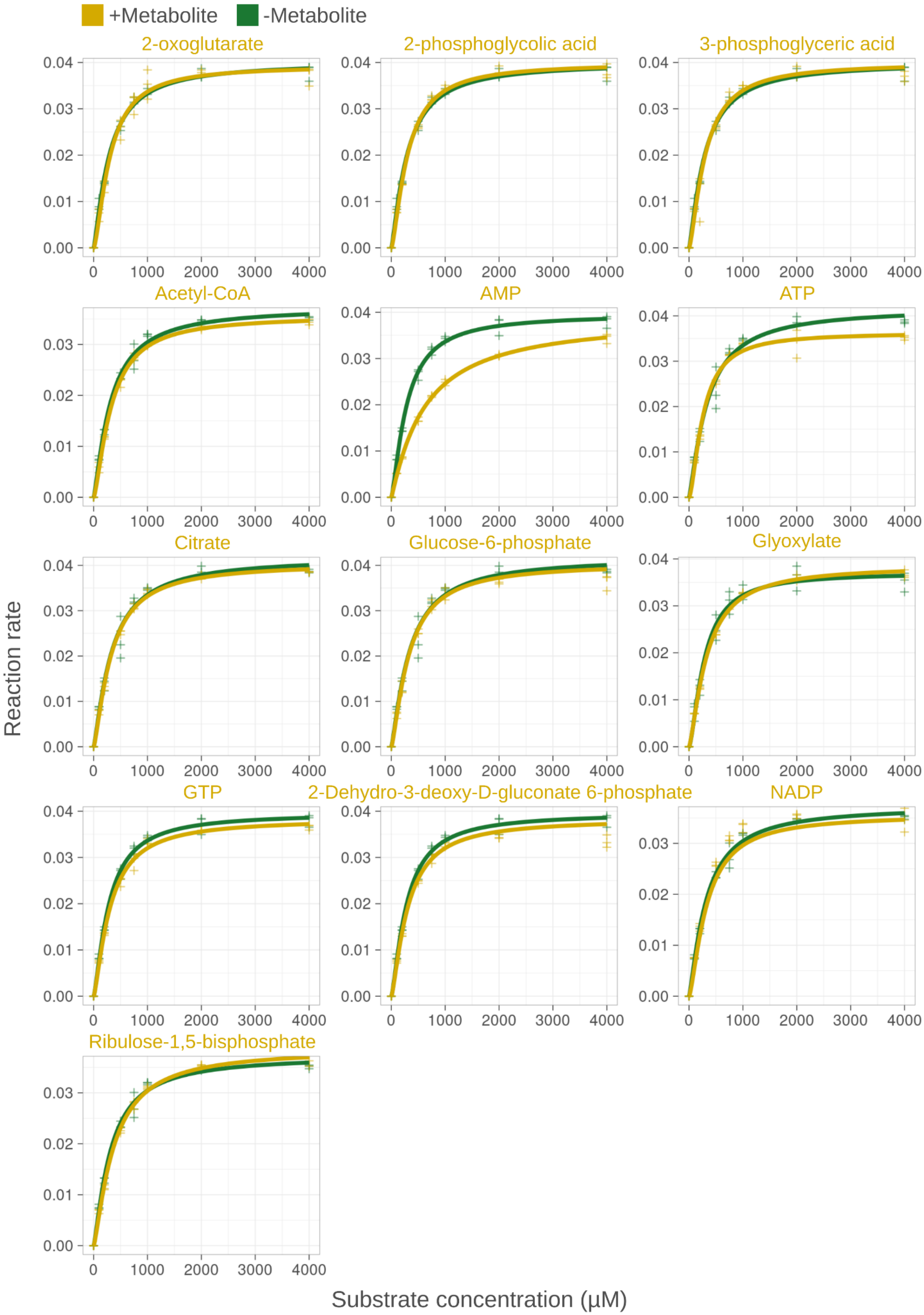

+Metabolite
-Metabolite
2-oxoglutarate
2-phosphoglycolic acid
3-phosphoglyceric acid
Acetyl-CoA
AMP
ATP
Citrate
Glucose-6-phosphate
Glyoxylate
GTP
2-Dehydro-3-deoxy-D-gluconate 6-phosphate
NADP
Ribulose-1,5-bisphosphate
Reaction rate
Substrate concentration (μM)

**Figure S18. Effect of different metabolites (1 mM) on the kinetics of *Synechocystis* transketolase.** The conversion of D-ribose-5-phosphate (substrate) and L-erythrulose to sedoheptulose-7-phosphate and glycolaldehyde was measured through the consumption of NADH by alcohol dehydrogenase when reducing glycolaldehyde to ethylene glycol. Each kinetic profile was characterized by measuring reaction rates for eight different substrate. Each kinetic profile was characterized by measuring reaction rates at eight different substrate concentrations in triplicates. Separate control reactions were run in parallel for each metabolite test.

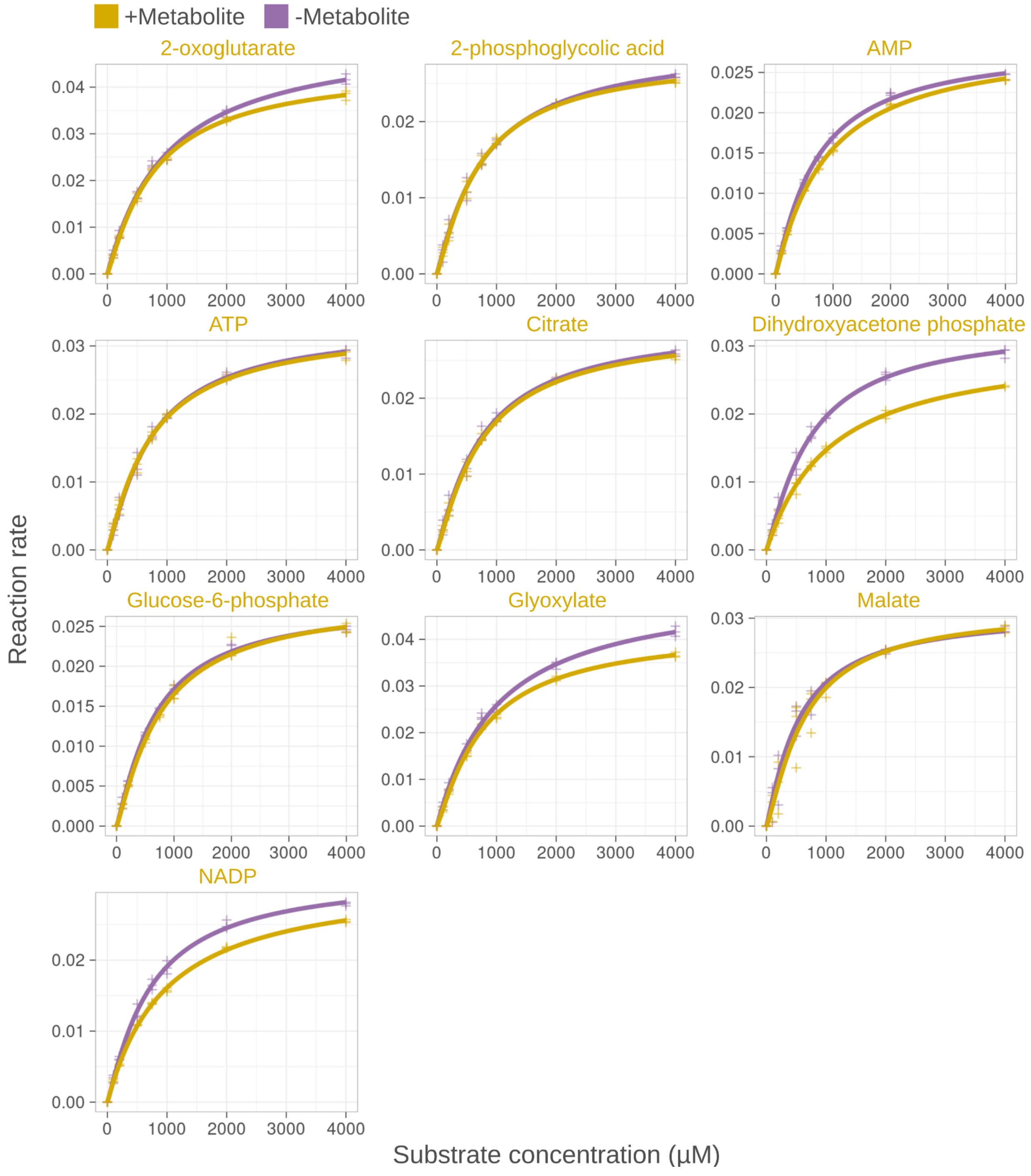


**Figure S19. Effect of different metabolites (1 mM) on the kinetics of *Cupriavidus* transketolase.** The conversion of D-ribose-5-phosphate (substrate) and L-erythrulose to sedoheptulose-7-phosphate and glycolaldehyde was measured through the consumption of NADH by alcohol dehydrogenase when reducing glycolaldehyde to ethylene glycol in a coupled spectrophotometric enzyme assay.g Initial rates were measured at eight different substrate concentrations in triplicates. Lines represent data fit to the enzymatic Hill equation.

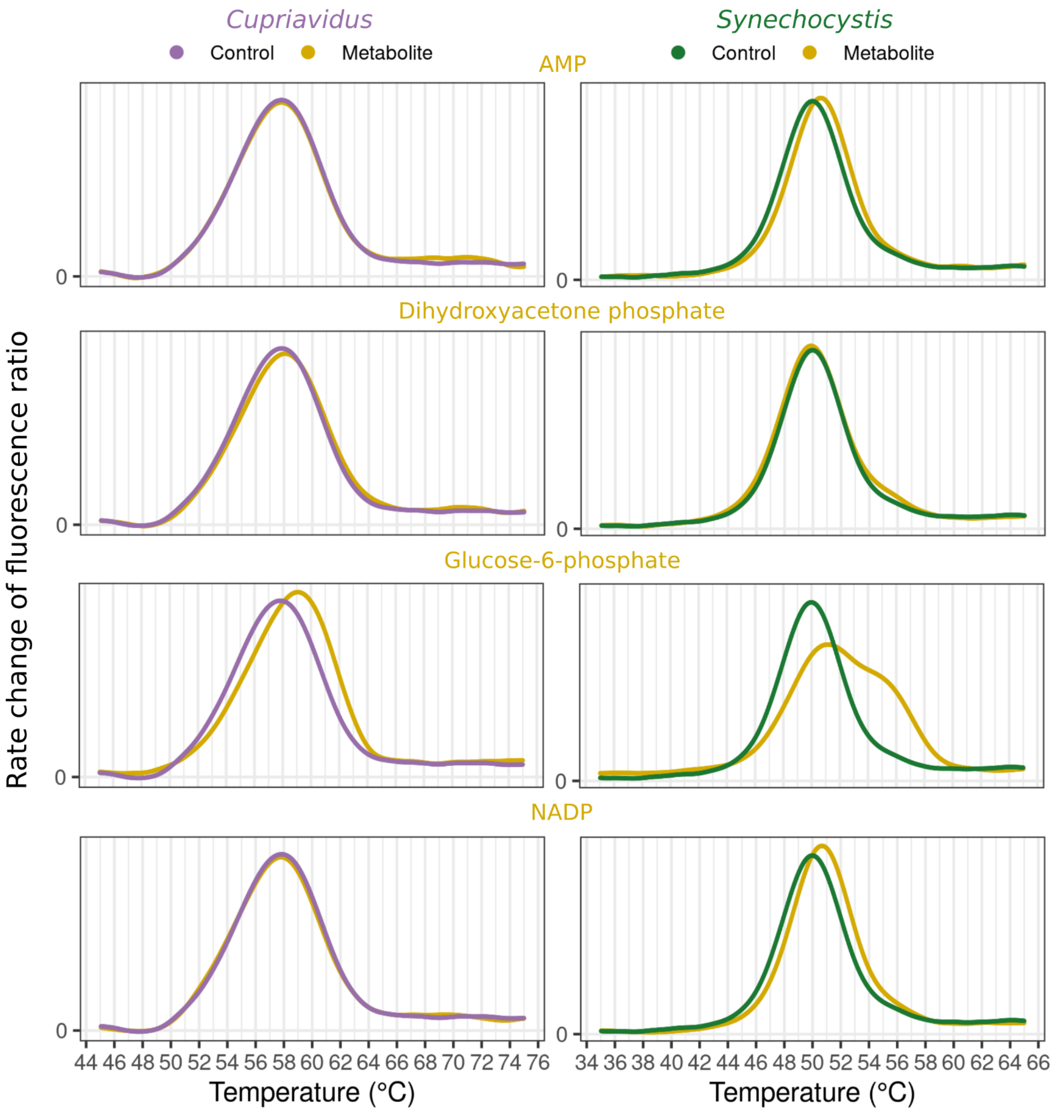


**Figure S20. Thermal shift assays of *Synechocystis* and *Cupriavidus* transketolase in the presence of various metabolites.** All metabolites were tested at a concentration of 1 mM. Curves indicate denaturation of transketolase over a temperature gradient of 1 °C/min. Y-axis shows the change in the ratio of protein autofluorescence (350 nm/330 nm), and maximum values indicate the melting temperature ($T_m$) at which half of the enzymes are denatured.

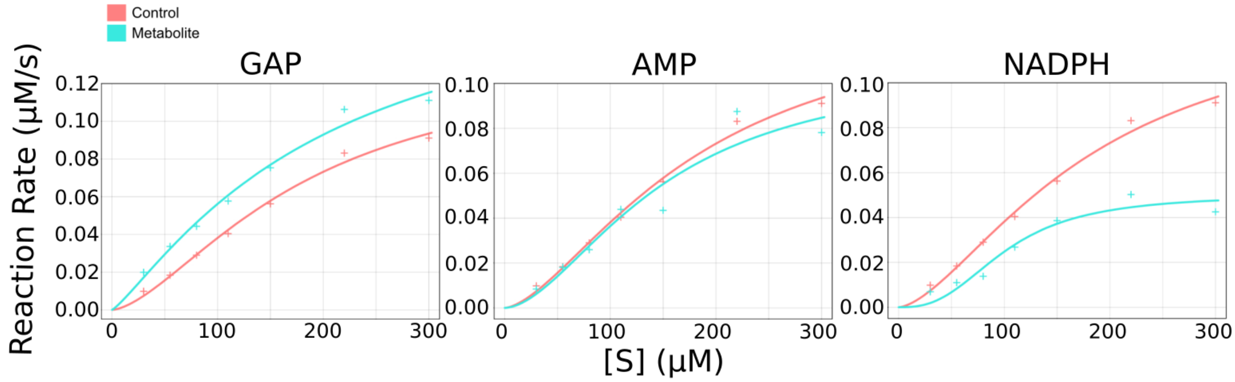


**Figure S21. Kinetic analysis of the *Synechocystis* F/SBPase R194H mutant.** The enzyme is AMP insensitive, but retains sensitivity to GAP and NADPH, consistent with binding sites of GAP and NADPH as detected by LiP being distinct from AMP binding site.

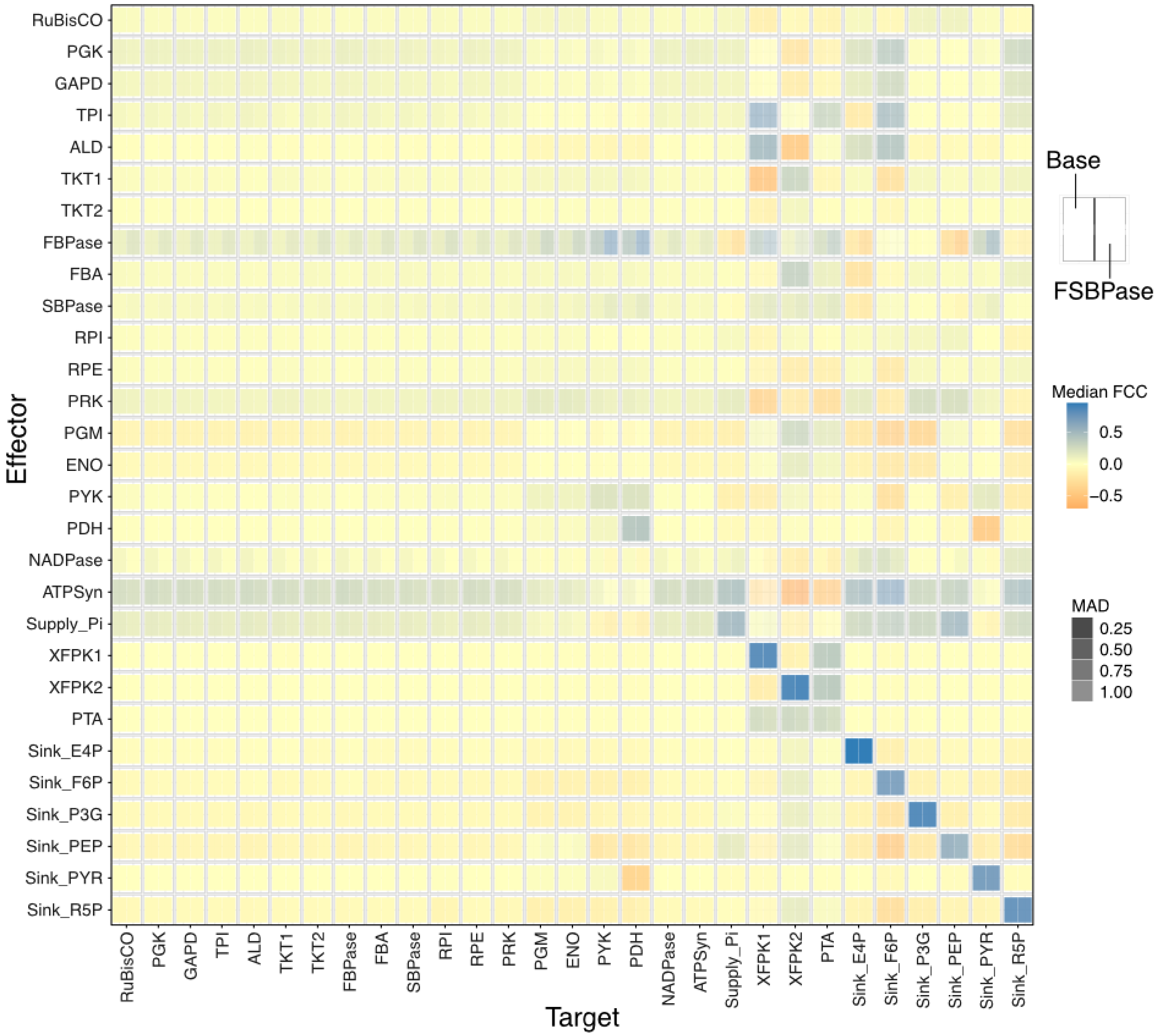


**Figure S22. Flux control coefficients for all reactions in the model.** Median FCCs and MAD values were calculated over all stable parameter sets for both variants as described in the methods section.

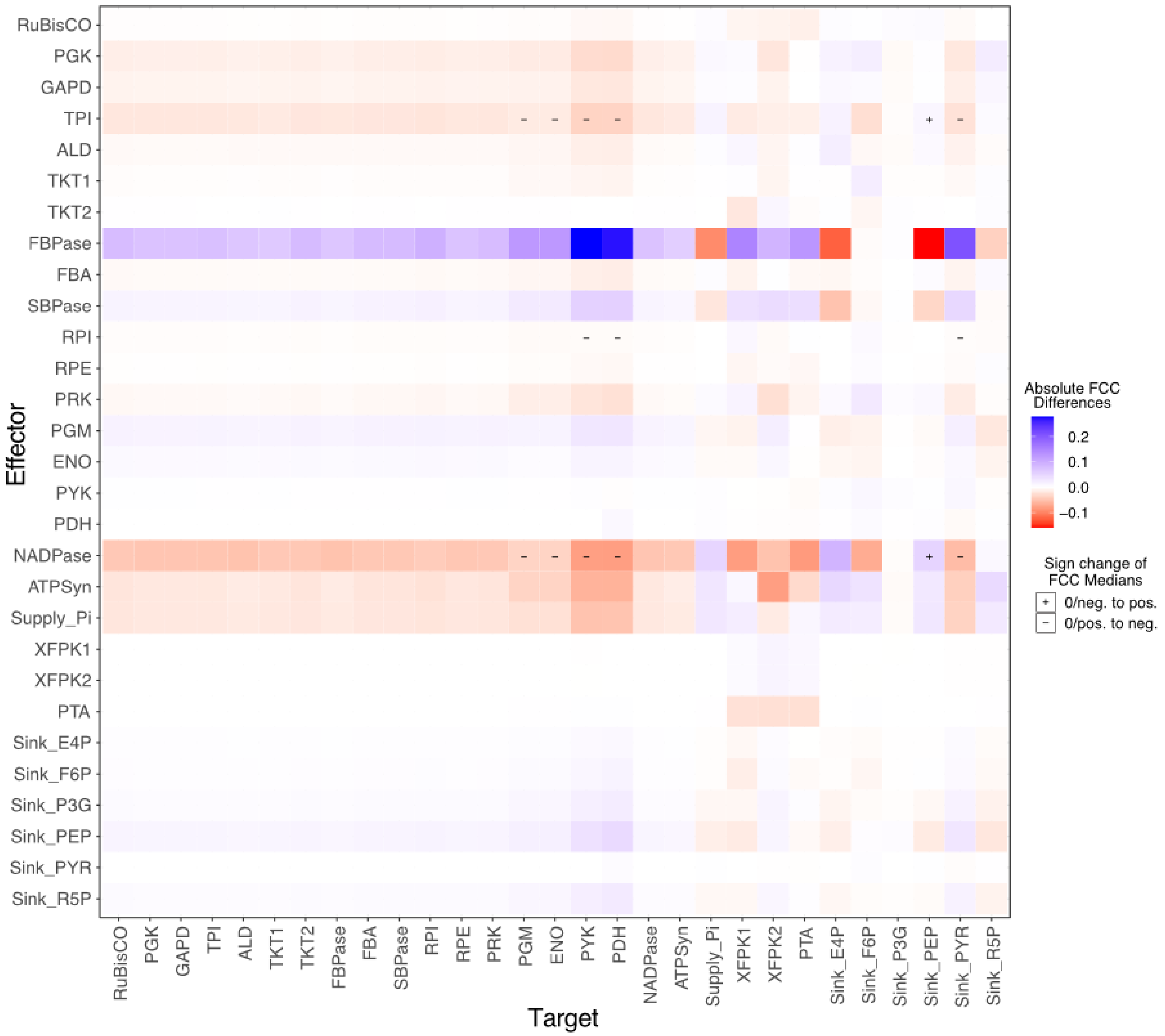


**Figure S23. Difference between median FCCs between model variants.** Median FCC of F/SBPase model subtracted by median FCC of base model for each Target/Effector pair. Plus signs indicate cases where the median FCC changed to a positive influence upon added F/SBPase regulation, whereas minus signs indicate cases where the FCC changed to a negative influence.

**Table S1. Chosen concentrations (mM) for every used metabolite and boundary values found in literature.**
The metabolite concentrations chosen for the LiP-SMap experiments in mM, the highest and lowest concentration found in literature, in mM (**Table S2**), and the highest and lowest concentrations allowed in the thermodynamically constrained model of Asplund-Samuelsson et al. 2018. In addition to the values shown below, several other reports have shown that metabolite concentrations can vary strongly across different conditions (Lempp et al. 2019; Marcus, Harel, and Kaplan 1983). As such, the "high" tested concentration was generally set higher than reported cellular concentration to capture metabolite accumulation that may occur during nutrient stress or metabolic perturbations and act as a regulatory signal.

| Metabolite | | Concentration ranges in literature | | | | Chosen concentrations | |
|---|---|---|---|---|---|---|---|
| Name | KEGG | Literature range lower limit | Literature range upper limit | Modeling range lower limit | Modeling range upper limit | Low | High |
| 2-phosphoglycolate | C00988 | 0.17 | 0.17 | 0.0001 | 100.0000 | 0.2 | 4 |
| 2-oxoglutarate | C00026 | 0.19 | 3.4 | 0.0031 | 2.1200 | 1 | 10 |
| 3-phosphoglycerate | C00597 | 1.54 | 19.02 | - | - | 2 | 20 |
| 6-phosphogluconate | C00345 | 0.07 | 9.58 | 0.0070 | 16.3800 | 1 | 10 |
| Acetyl-CoA | C00024 | 0.01 | 1.54 | 0.0001 | 0.9640 | 1 | 10 |
| ADP | C00008 | 0.58 | 5.26 | 0.0428 | 4.1867 | 1 | 10 |
| AMP | C00020 | 0.34 | 13.4 | 0.0470 | 11.2000 | 1 | 10 |
| ATP | C00002 | 0.2 | 3.06 | 0.0300 | 43.4300 | 2 | 32 |
| cAMP | C00575 | 0.34 | 0.34 | - | - | 0.5 | 5 |
| Citrate | C00158 | 0.17 | 5.45 | 0.0240 | 2.4800 | 2 | 20 |
| Fructose-1,6-bisphosphate | C00354 | 0.17 | 4.97 | 0.0163 | 7.0694 | 1 | 10 |
| Glucose-6-phosphate | C00668 | 0.17 | 3.4 | - | - | 1 | 10 |
| Glyceraldehyde-3-phosphate | C00118 | 0.17 | 0.65 | 0.0001 | 100.0000 | 0.5 | 5 |
| Glycolate | C00160 | 0.09 | 0.17 | - | - | 1 | 10 |
| Glyoxylate | C00048 | 0.17 | 0.17 | 0.0001 | 100.0000 | 1 | 10 |
| GTP | C00044 | 0.14 | 4.87 | 0.1595 | 1.0358 | 1 | 10 |
| KDPG | C04442 | - | - | - | - | 0.5 | 5 |
| Malate | C00149 | 0.17 | 6.79 | 0.0142 | 2.0602 | 1 | 10 |
| NADP | C00006 | 0.17 | 2.14 | 0.0055 | 1.3200 | 0.5 | 5 |
| NADPH | C00005 | 0.14 | 0.24 | 0.0001 | 49.4100 | 0.5 | 5 |
| Phosphoenolpyruvate | C00074 | 0.51 | 6.79 | 0.1700 | 2.9900 | 0.5 | 10 |
| Phenylalanine | C00079 | 0.12 | 0.25 | 0.0151 | 0.0955 | 0.5 | 5 |
| Ribulose-5-phosphate | C00199 | 0.03 | 5.09 | 0.0077 | 3.8900 | 1 | 10 |
| Ribulose-1,5-bisphosphate | C01182 | 0.05 | 17.15 | 0.0001 | 11.2311 | 1 | 10 |
| Sucrose | C00089 | - | - | - | - | 1 | 10 |

**Table S2. All metabolite concentrations found across 7 metabolomics studies in mM.**
Absolute metabolite concentrations found in literature. All values obtained from articles studying cyanobacteria were converted from μmol per gram cell dry weight to millimolar. This was done by calculating the amount of cell volume per gram dry weight from the values reported by Zavřel et al. for a growth rate of 0.05 $h^{-1}$ (Zavřel et al. 2019). The cell volume was calculated from the cell diameter and multiplied by the cell count per liter culture to obtain the total cell volume per liter culture. This was then divided by the dry weight per liter cell culture, giving the amount of cell volume per dry weight.

| Metabolite | | E.coli | PCC. 6803 | | | | | |
|---|---|---|---|---|---|---|---|---|
| Name | KEGG | Bennet (2009) | Nishiguchi (2019) | Yoshikawa (2013) | Takahashi (2008) | Shastri (2007) | Hasunuma (2013) | Dempo (2014) |
| 2-phosphoglycolate | C00988 | - | - | - | - | 0.17 | - | - |
| 2-oxoglutarate | C00026 | 0.44 | - | 0.71 | 0.19 | 3.40 | - | - |
| 3-phosphoglycerate | C00597 | 1.54 | 12.24 | 2.26 | 2.38 | 10.19 | 2.21 | 19.02 |
| 6-phosphogluconate | C00345 | 1.64 | 0.07 | 0.15 | 0.08 | 0.17 | - | 9.58 |
| Acetyl-CoA | C00024 | 0.73 | 1.54 | 0.27 | - | 0.17 | 0.01 | 0.70 |
| ADP | C00008 | 0.56 | 0.58 | 1.12 | 0.85 | - | 1.87 | 5.26 |
| AMP | C00020 | 0.28 | - | 0.76 | 0.34 | - | 1.51 | 13.40 |
| ATP | C00002 | 9.63 | 1.34 | 13.12 | 1.02 | - | 0.20 | 3.06 |
| cAMP | C00575 | 0.08 | - | - | - | - | 0.34 | - |
| Citrate | C00158 | 0.85 | 5.45 | 1.70 | 0.85 | 0.17 | 0.49 | 3.23 |
| Fructose-1,6-bisphosphate | C00354 | 15.20 | 0.51 | 4.97 | 0.17 | 0.17 | - | 0.42 |
| Glucose-6-phosphate | C00668 | - | 1.70 | 2.46 | 0.17 | 3.40 | 0.49 | 3.24 |
| Glyceraldehyde-3-phosphate | C00118 | - | - | - | 0.17 | 0.17 | - | 0.65 |
| Glycolate | C00160 | - | - | - | - | 0.17 | - | 0.09 |
| Glyoxylate | C00048 | - | - | - | - | 0.17 | - | - |
| GTP | C00044 | 4.87 | - | 0.90 | - | - | - | 0.14 |
| KDPG | C04442 | - | - | 0.00 | - | - | - | - |
| Malate | C00149 | - | 0.49 | 0.48 | 0.17 | 6.79 | 0.19 | 0.20 |
| NADP | C00006 | 0.00 | 2.14 | 1.17 | 0.17 | - | - | 0.97 |
| NADPH | C00005 | 0.12 | 0.14 | - | 0.24 | - | - | - |
| Phosphoenolpyruvate | C00074 | 0.18 | 6.79 | 1.49 | 0.51 | 6.79 | 1.09 | 2.90 |
| Phenylalanine | C00079 | 0.04 | - | 0.12 | - | - | 0.13 | 0.25 |
| Ribulose-5-phosphate | C00199 | - | 0.02 | 0.37 | 0.10 | 5.09 | 0.05 | 0.03 |
| Ribulose-1,5-bisphosphate | C01182 | - | 0.85 | 5.91 | 0.05 | 17.15 | - | 0.48 |
| Sucrose | C00089 | - | - | - | - | - | - | - |

**References:** (Bennett et al. 2009; Nishiguchi et al. 2019; Yoshikawa et al. 2013; Shastri and Morgan 2007; Hasunuma et al. 2013; Dempo et al. 2014; Takahashi, Uchimiya, and Hihara 2008)

**Table S3. Effect of metabolites on kinetic parameters for *Synechocystis* and *Cupriavidus* F/SBPase.** See separate file: TableS3_summary_stats_FSBPase_kinetics.xlsx. Table shows mean and standard deviation of kinetic parameters for syn-F/SBPase and cn-F/SBPase, with and without added metabolite (2-3 replicate assays). P-values were calculated by comparing kinetic parameters with versus without added metabolite, using Student's t-tests. The columns named "*Max rate change"* and "*2nd highest rate change"* show the maximum and 2nd highest change in catalytic rate when metabolite is added, across all tested substrate concentrations.

**Table S4. Transition list for mass spectrometry.**

| Compound | Retention Time (min) | RT window (min) | Precursor (m/z) | Product (m/z) | Collision Energy (V) | Min Dwell Time (ms) |
|---|---|---|---|---|---|---|
| F6P | 4.5 | 9 | 259.022 | 138.979 | 20 | 38.505 |
| F6P | 4.5 | 9 | 259.022 | 96.969 | 35 | 38.505 |
| F6P | 4.5 | 9 | 259.022 | 78.959 | 40 | 38.505 |

Takahashi, Hideyuki, Hirofumi Uchimiya, and Yukako Hihara. 2008. "Difference in Metabolite Levels between

Photoautotrophic and Photomixotrophic Cultures of Synechocystis Sp. PCC 6803 Examined by Capillary Electrophoresis Electrospray Ionization Mass Spectrometry.” *Journal of Experimental Botany* 59 (11): 3009–18.
Yang, Mingkun, Hui Huang, and Feng Ge. 2019. “Lysine Propionylation Is a Widespread Post-Translational Modification Involved in Regulation of Photosynthesis and Metabolism in Cyanobacteria.” *International Journal of Molecular Sciences* 20 (19). https://doi.org/10.3390/ijms20194792.
Yoshikawa, Katsunori, Takashi Hirasawa, Kenichi Ogawa, Yuki Hidaka, Tsubasa Nakajima, Chikara Furusawa, and Hiroshi Shimizu. 2013. “Integrated Transcriptomic and Metabolomic Analysis of the Central Metabolism of Synechocystis Sp. PCC 6803 under Different Trophic Conditions.” *Biotechnology Journal* 8 (5): 571–80.
Zavřel, Tomáš, Marjan Faizi, Cristina Loureiro, Gereon Poschmann, Kai Stühler, Maria Sinetova, Anna Zorina, Ralf Steuer, and Jan Červený. 2019. “Quantitative Insights into the Cyanobacterial Cell Economy.” *eLife* 8 (February). https://doi.org/10.7554/eLife.42508.
