## Supplementals for "Metabolite interactions in the bacterial Calvin cycle and implications for flux regulation": Dataset S4. Phylogenetic trees of Calvin cycle enzymes labeled with detected protein-metabolite interactions.pdf

Each tree is based on fewer than 1 000 representative sequences and all sequences from the four organisms in this study, which belong to KEGG ortholog (KO) families catalyzing steps in the Calvin cycle (see Materials and Methods). The ring indicates organism group; Kingdom for eukaryotes, and phylum for bacteria and archaea, except for Proteobacteria, which are divided into classes. Organism groups that were too small to warrant their own color were grouped into the “Other” categories. Each tree is titled with the KO ID, gene names, enzyme names, and Enzyme Commission number. Sequences from the four organisms in this study are indicated by symbols and text with an organism-specific color. These sequences show additional information in the text boxes; The top text line indicates the UniProt ID, the gene name, and the locus ID (in parentheses). The bottom text line indicates all metabolites with which the enzyme has at least one significantly interacting peptide at low or high concentration. Scale bars indicate substitutions per site.

ALD

K01623 ALDO; fructose−bisphosphate aldolase, class I [EC:4.1.2.13]

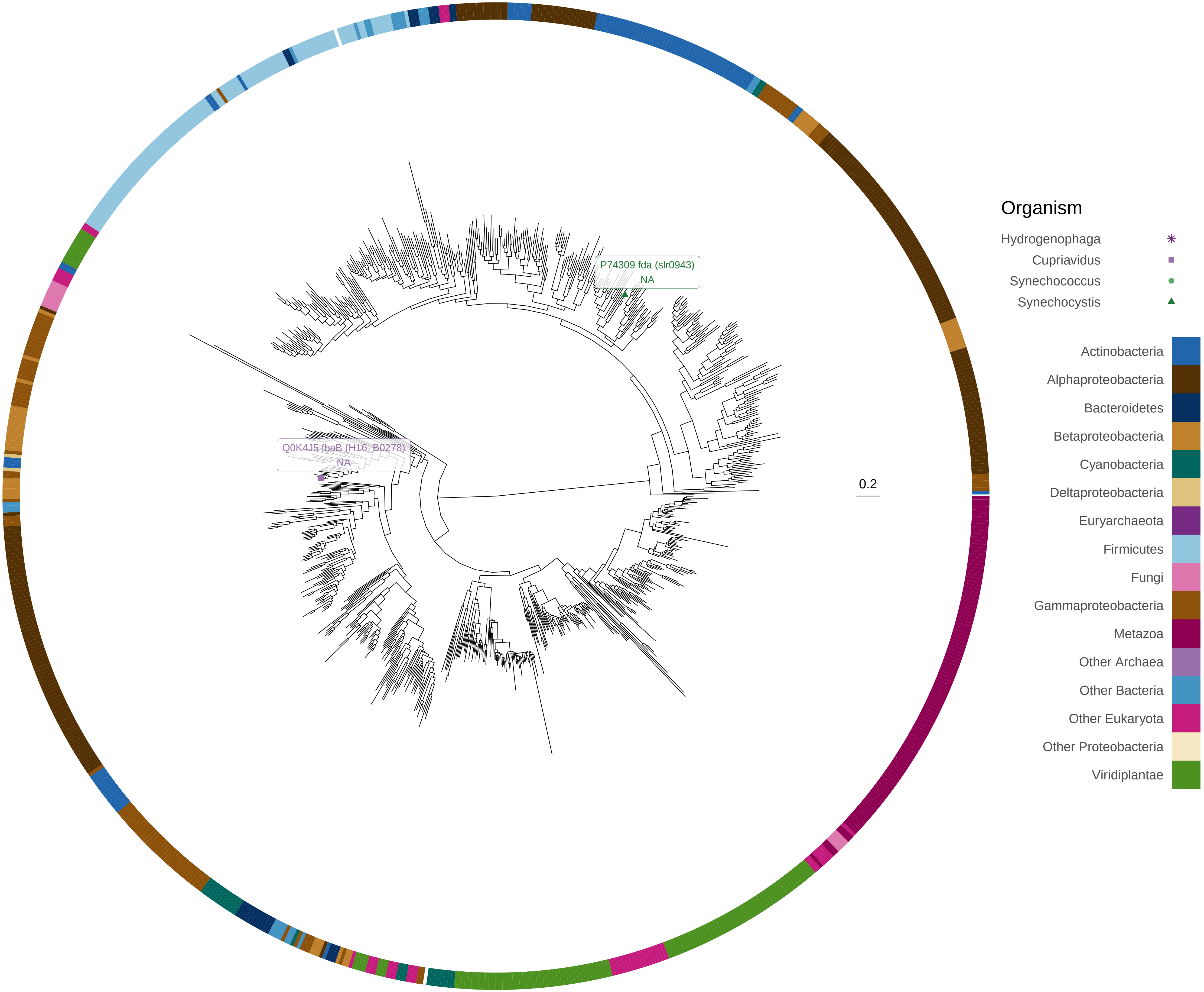

FBA

K01624 FBA, fbaA; fructose−bisphosphate aldolase, class II [EC:4.1.2.13]

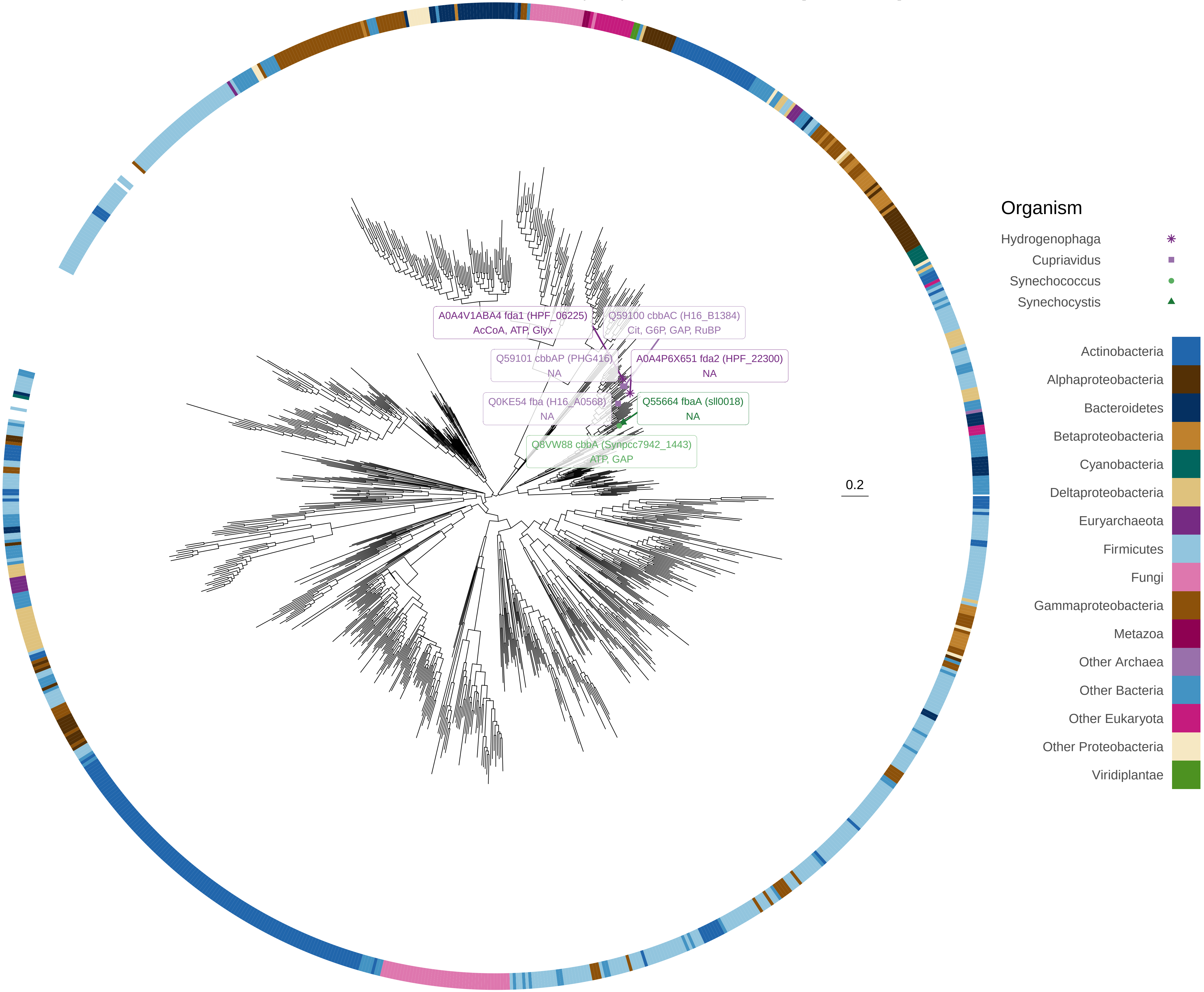

### FBP−SBPase_I

K01086 fbp−SEBP; fructose−1,6−bisphosphatase I / sedoheptulose−1,7−bisphosphatase [EC:3.1.3.11 3.1.3.37]

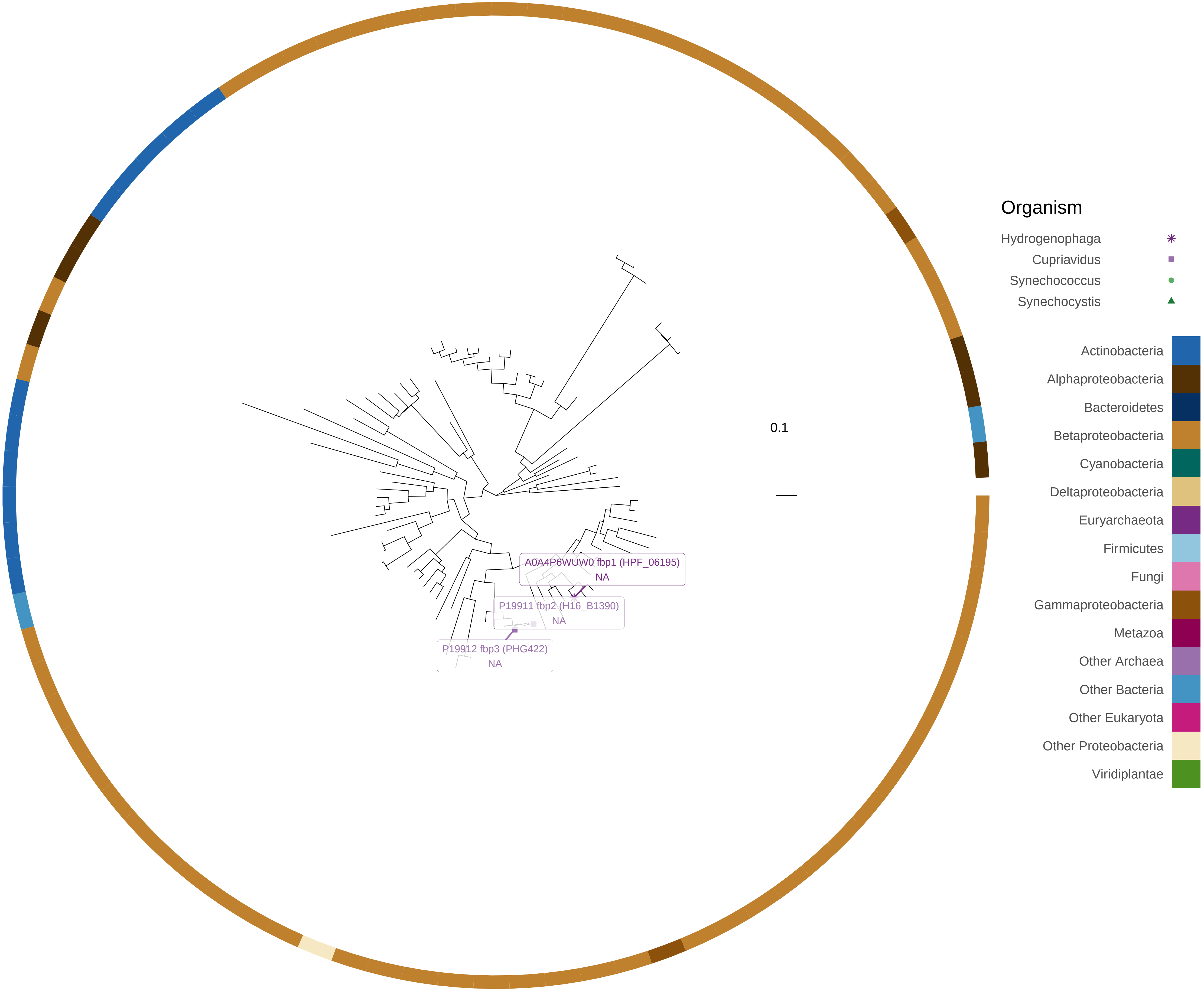

FBP−SBPase_II_glpX

K11532 glpX−SEBP; fructose−1,6−bisphosphatase II / sedoheptulose−1,7−bisphosphatase [EC:3.1.3.11 3.1.3.37]

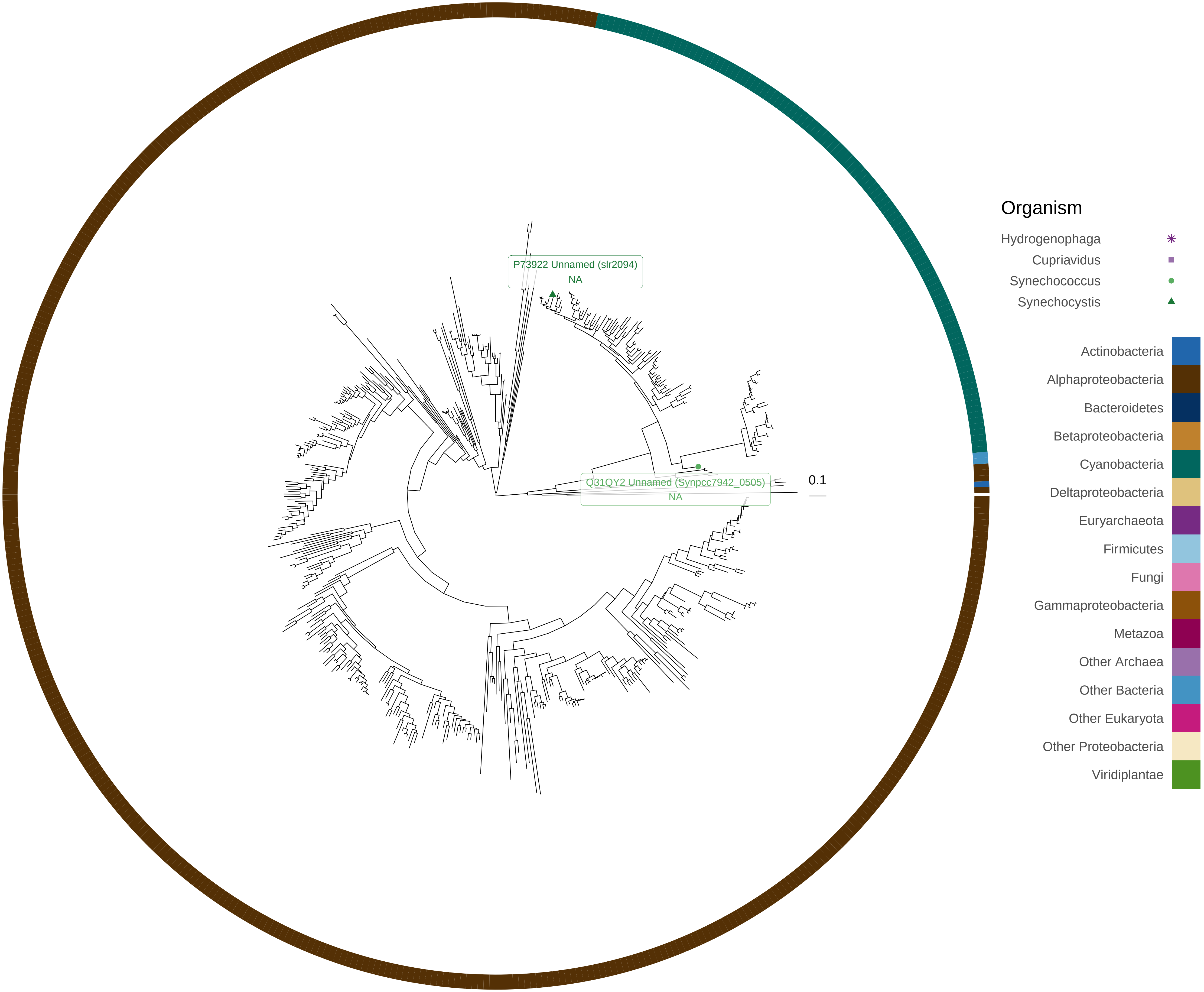

FBPase_I

K03841 FBP, fbp; fructose−1,6−bisphosphatase I [EC:3.1.3.11]

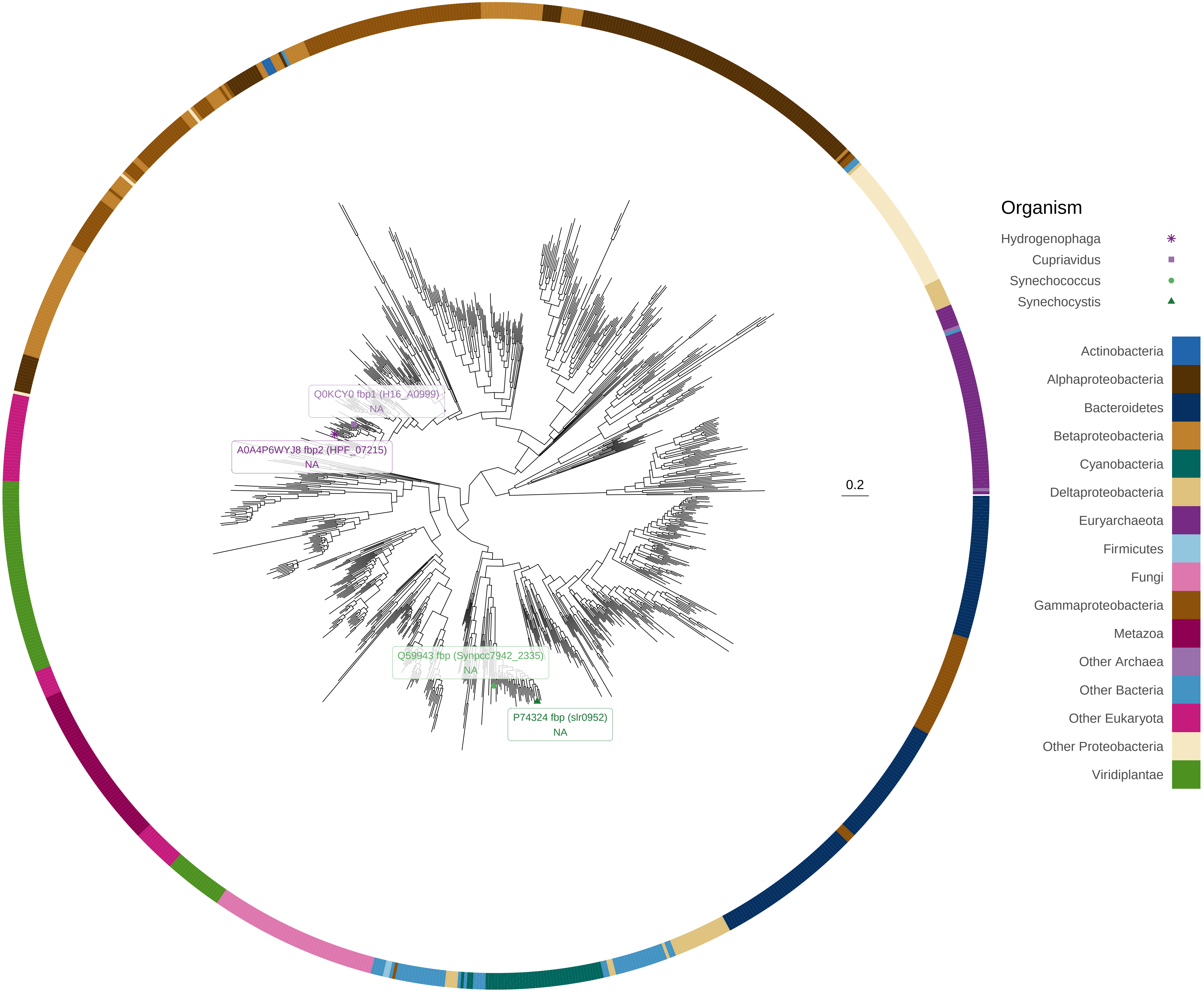

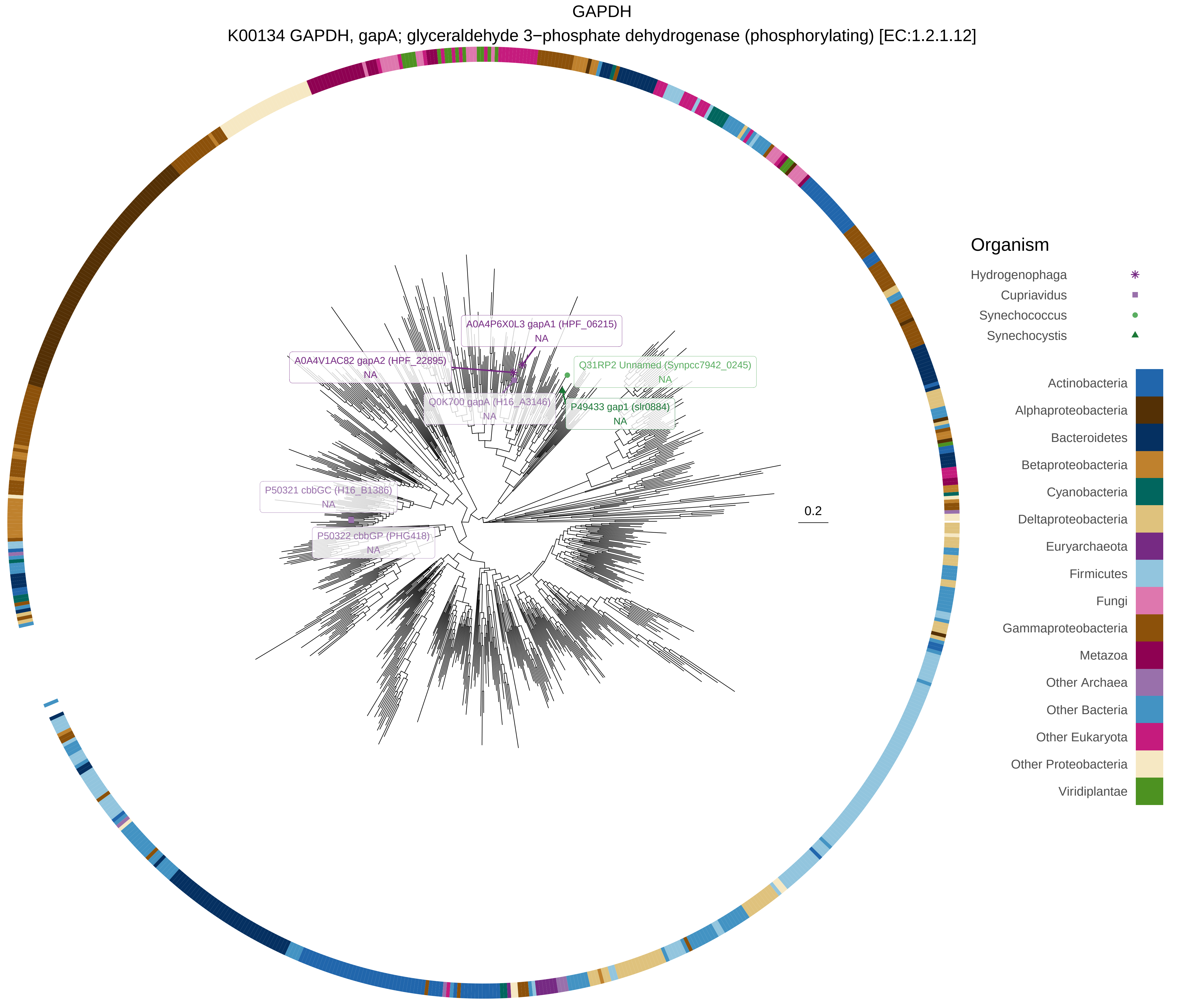
GAPDH
K00134 GAPDH, gapA; glyceraldehyde 3−phosphate dehydrogenase (phosphorylating) [EC:1.2.1.12]
A0A4P6X0L3 gapA1 (HPF_06215)
NA
A0A4V1AC82 gapA2 (HPF_22895)
NA
Q31RP2 Unnamed (Synpcc7942_0245)
NA
Q0K700 gapA (H16_A3146)
NA
P49433 gap1 (slr0884)
NA
P50321 cbbGC (H16_B1386)
NA
P50322 cbbGP (PHG418)
NA
0.2
Organism
Hydrogenophaga
Cupriavidus
Synechococcus
Synechocystis
Actinobacteria
Alphaproteobacteria
Bacteroidetes
Betaproteobacteria
Cyanobacteria
Deltaproteobacteria
Euryarchaeota
Firmicutes
Fungi
Gammaproteobacteria
Metazoa
Other Archaea
Other Bacteria
Other Eukaryota
Other Proteobacteria
Viridiplantae

GAPDH2

K00150 gap2; glyceraldehyde−3−phosphate dehydrogenase (NAD(P)) [EC:1.2.1.59]

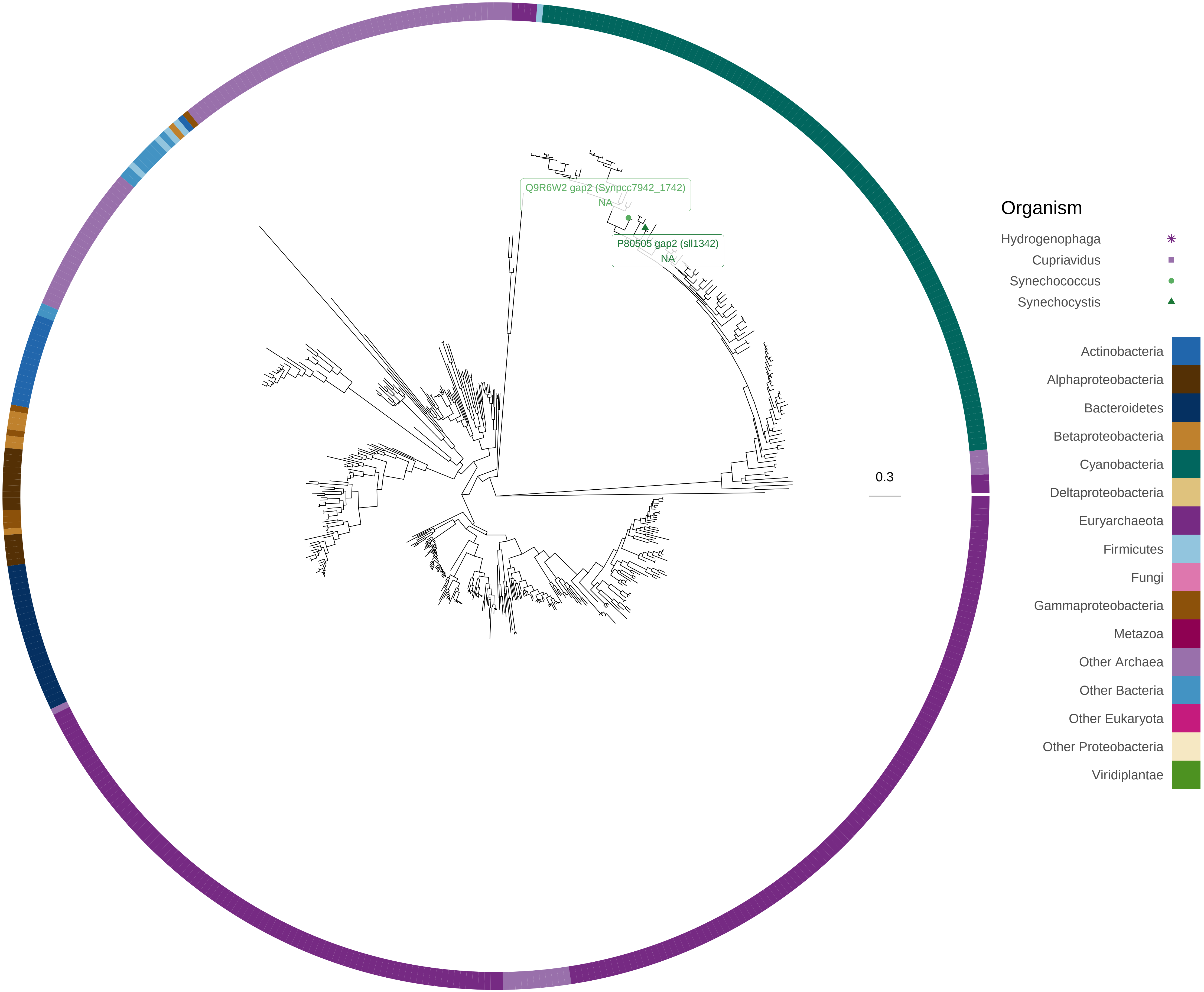

PGK

K00927 PGK, pgk; phosphoglycerate kinase [EC:2.7.2.3]

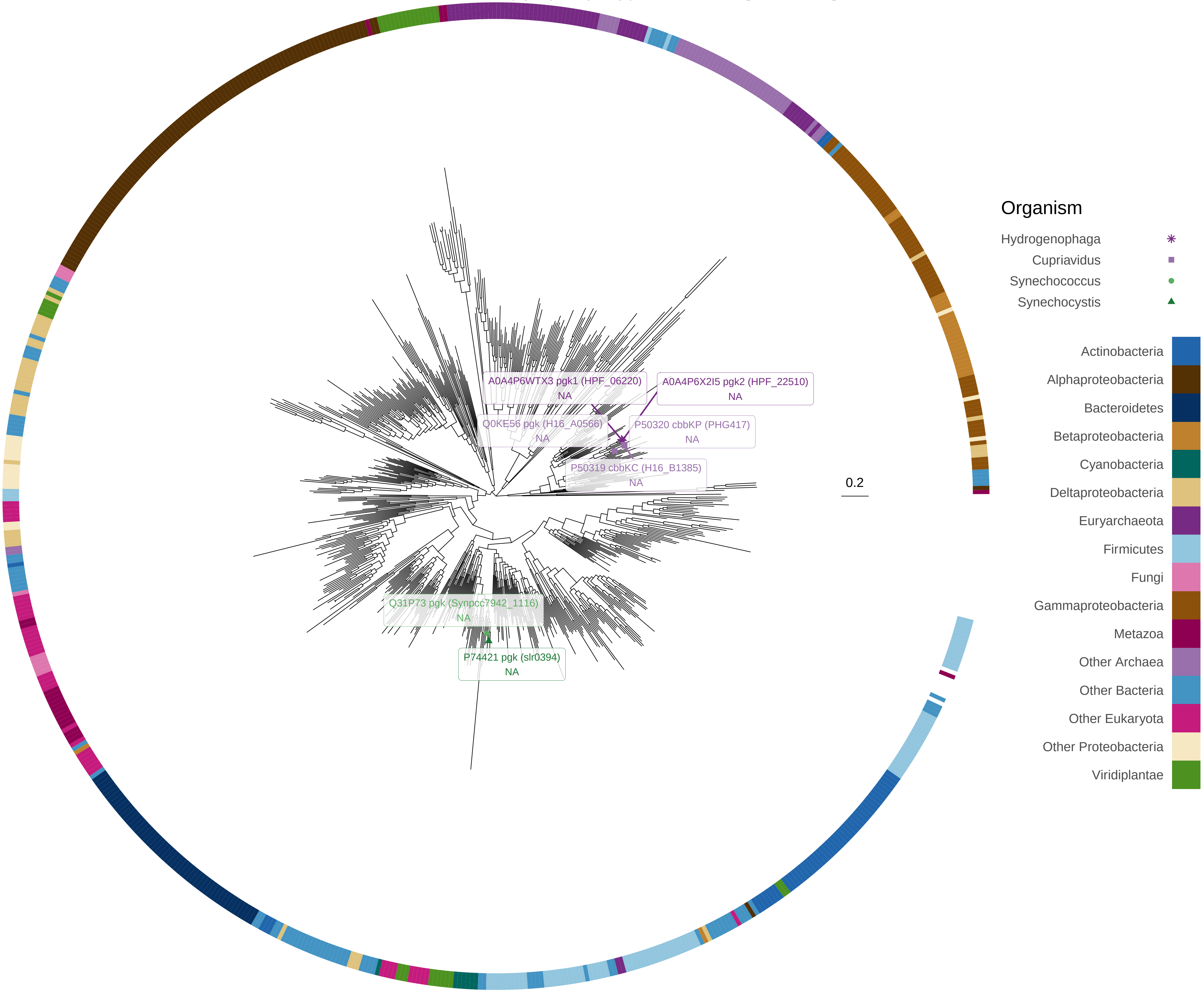

### PRK

K00855 PRK, prkB; phosphoribulokinase [EC:2.7.1.19]

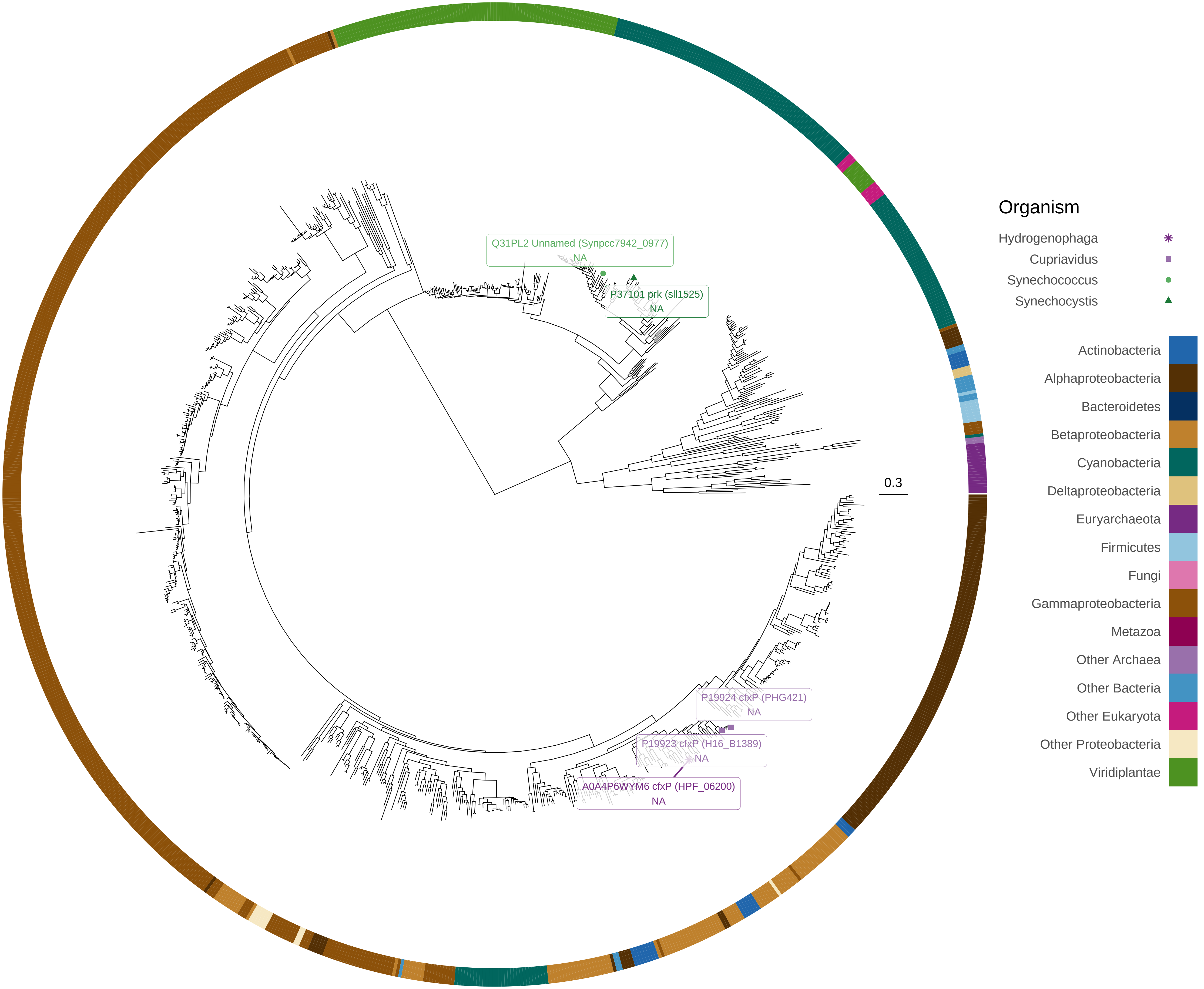

rbcL

K01601 rbcL, cbbL; ribulose−bisphosphate carboxylase large chain [EC:4.1.1.39]

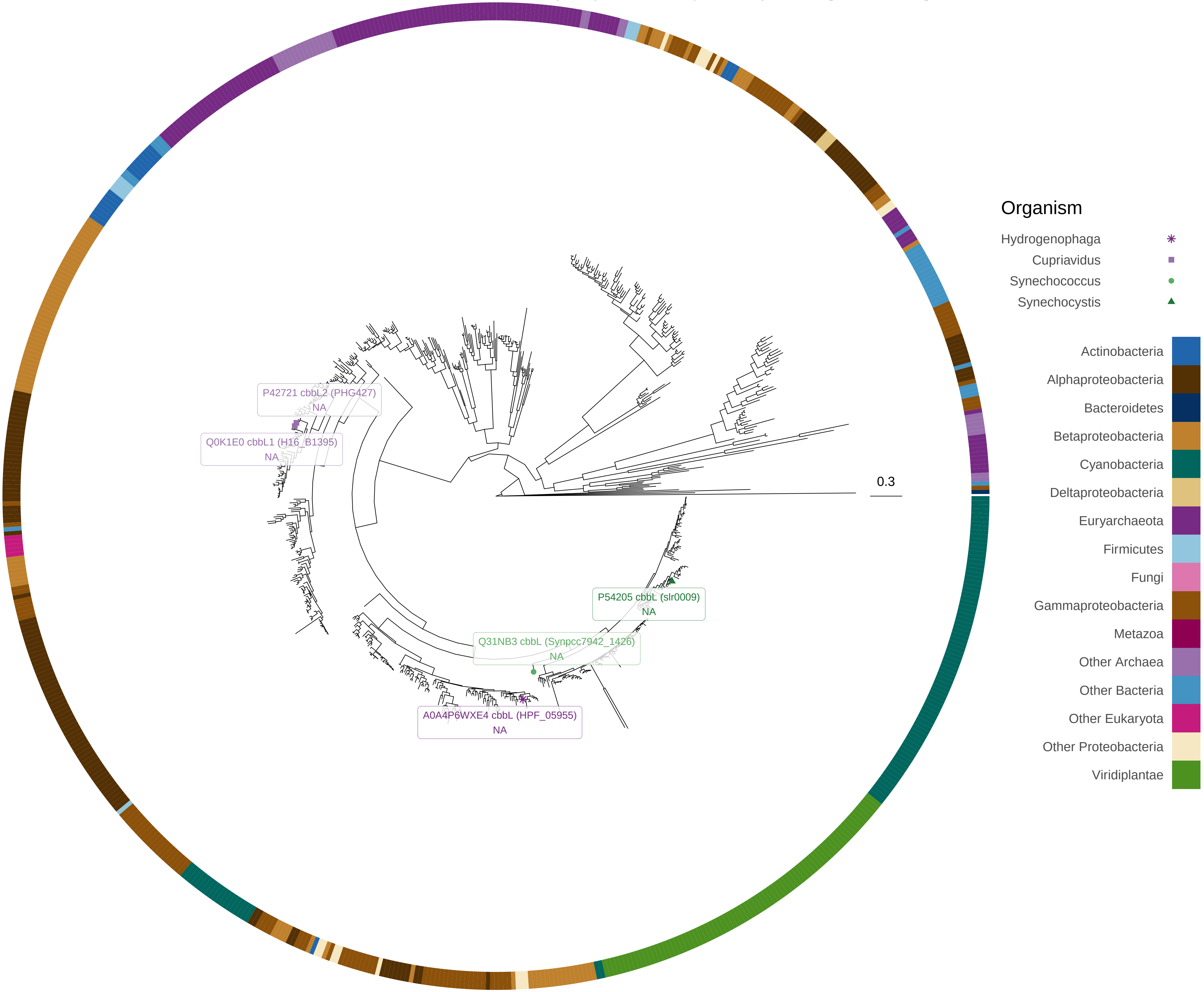

rbcS

K01602 rbcS, cbbS; ribulose−bisphosphate carboxylase small chain [EC:4.1.1.39]

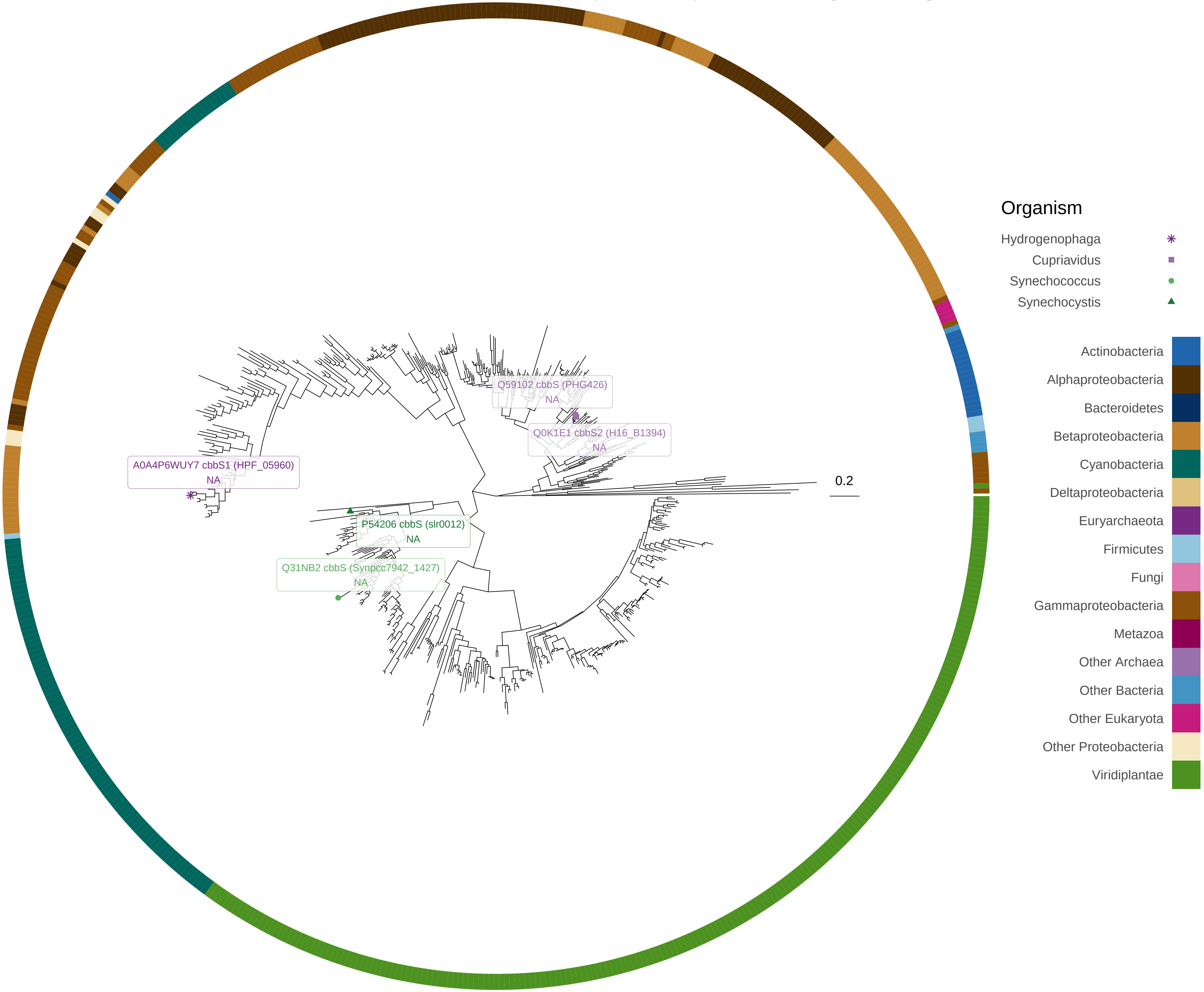

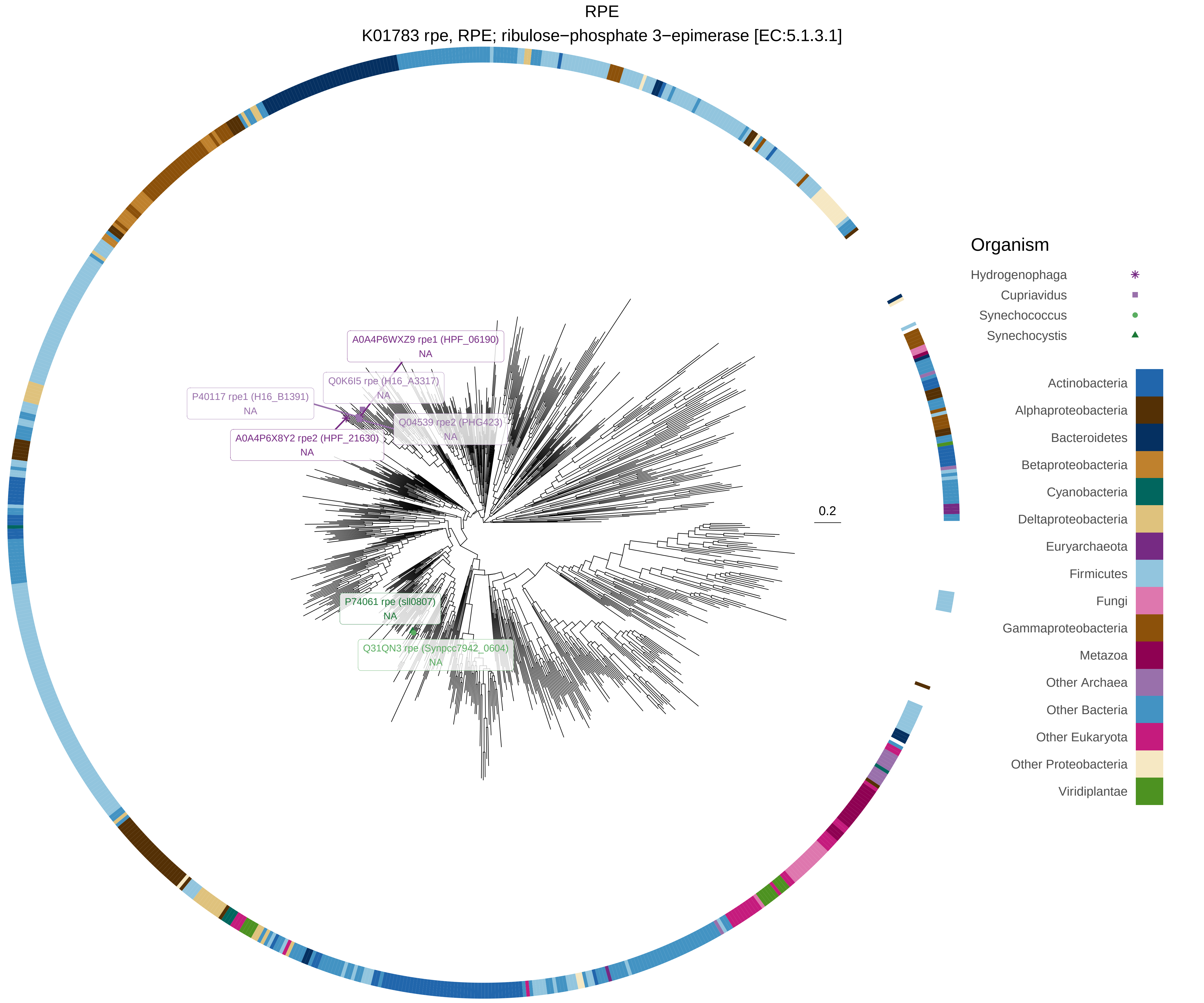
RPE
K01783 rpe, RPE; ribulose−phosphate 3−epimerase [EC:5.1.3.1]
A0A4P6WXZ9 rpe1 (HPF_06190)
NA
Q0K6I5 rpe (H16_A3317)
NA
P40117 rpe1 (H16_B1391)
NA
Q04539 rpe2 (PHG423)
NA
A0A4P6X8Y2 rpe2 (HPF_21630)
NA
P74061 rpe (sll0807)
NA
Q31QN3 rpe (Synpcc7942_0604)
NA
0.2
Organism
Hydrogenophaga
Cupriavidus
Synechococcus
Synechocystis
Actinobacteria
Alphaproteobacteria
Bacteroidetes
Betaproteobacteria
Cyanobacteria
Deltaproteobacteria
Euryarchaeota
Firmicutes
Fungi
Gammaproteobacteria
Metazoa
Other Archaea
Other Bacteria
Other Eukaryota
Other Proteobacteria
Viridiplantae

rpiA

K01807 rpiA; ribose 5−phosphate isomerase A [EC:5.3.1.6]

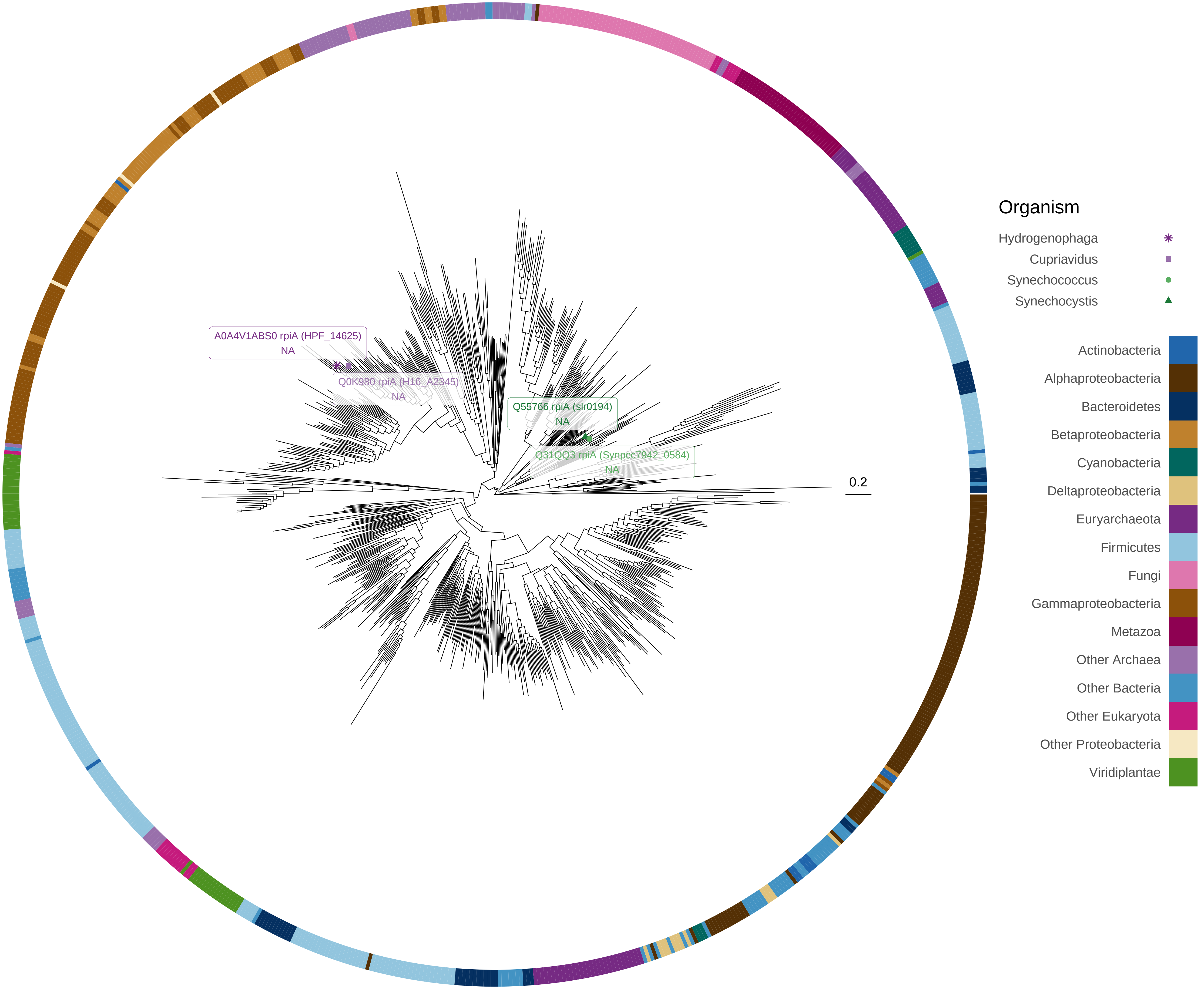

rpiB

K01808 rpiB; ribose 5−phosphate isomerase B [EC:5.3.1.6]

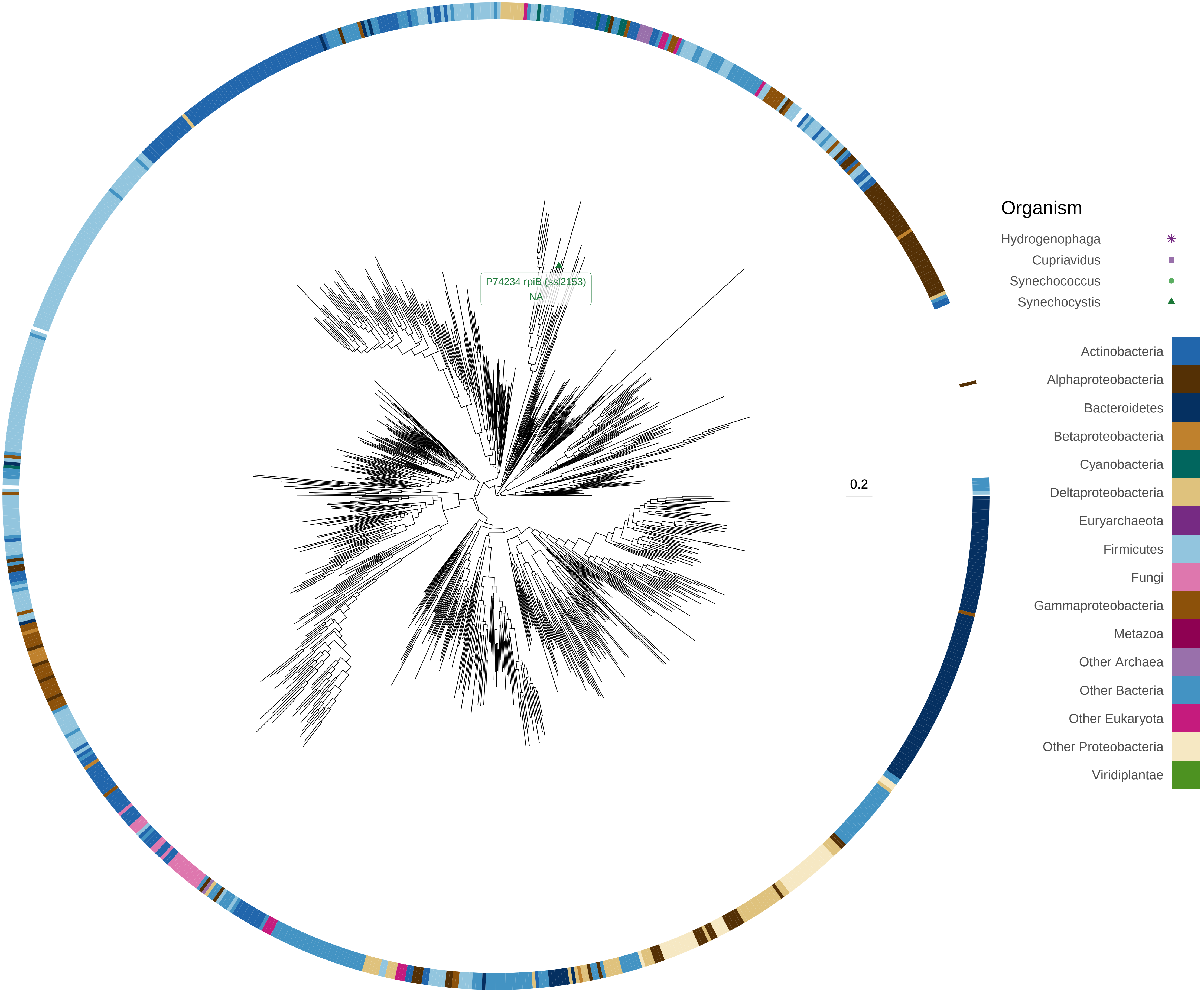

TAL

K00616 E2.2.1.2, talA, talB; transaldolase [EC:2.2.1.2]

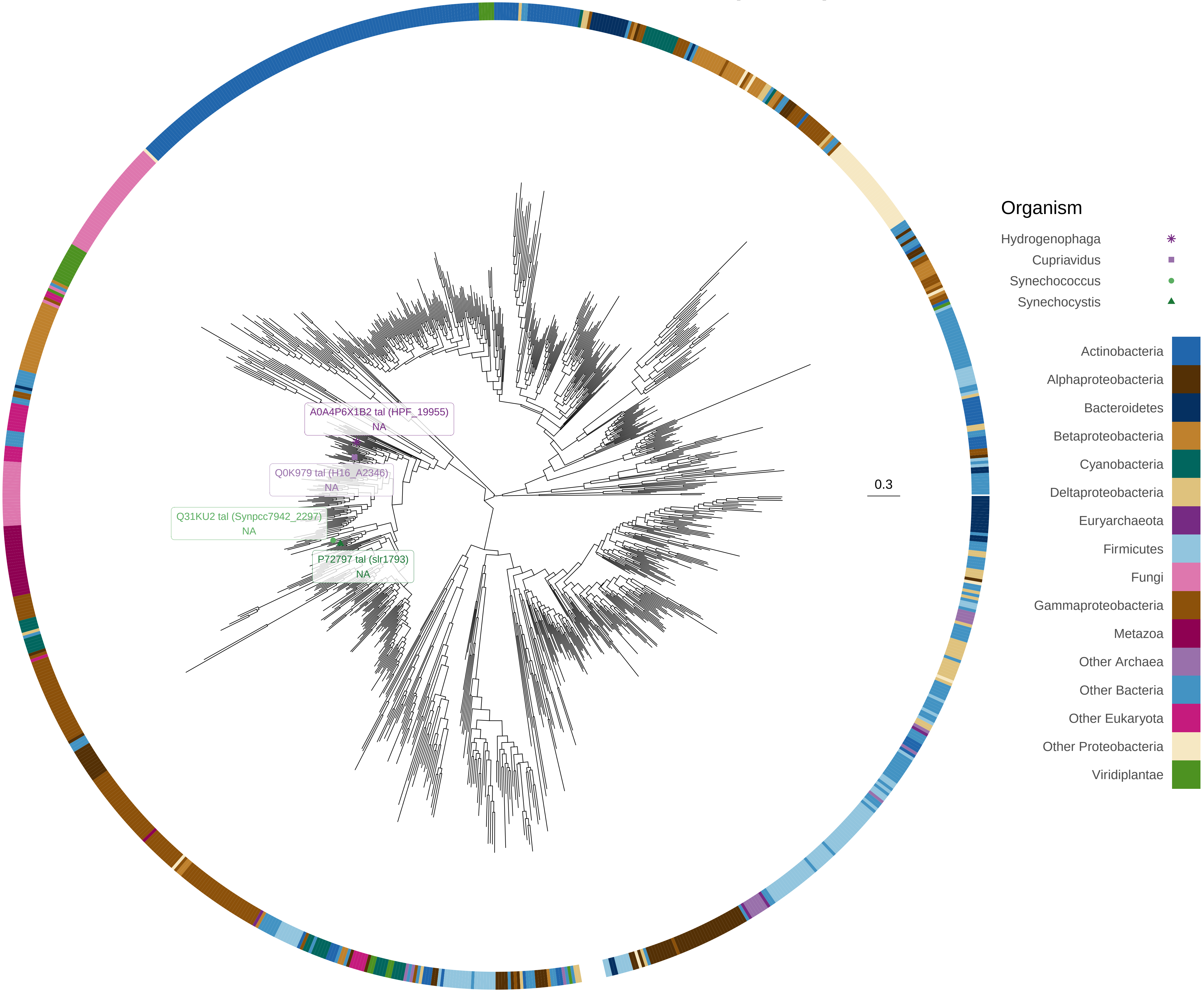

TKT

K00615 E2.2.1.1, tktA, tktB; transketolase [EC:2.2.1.1]

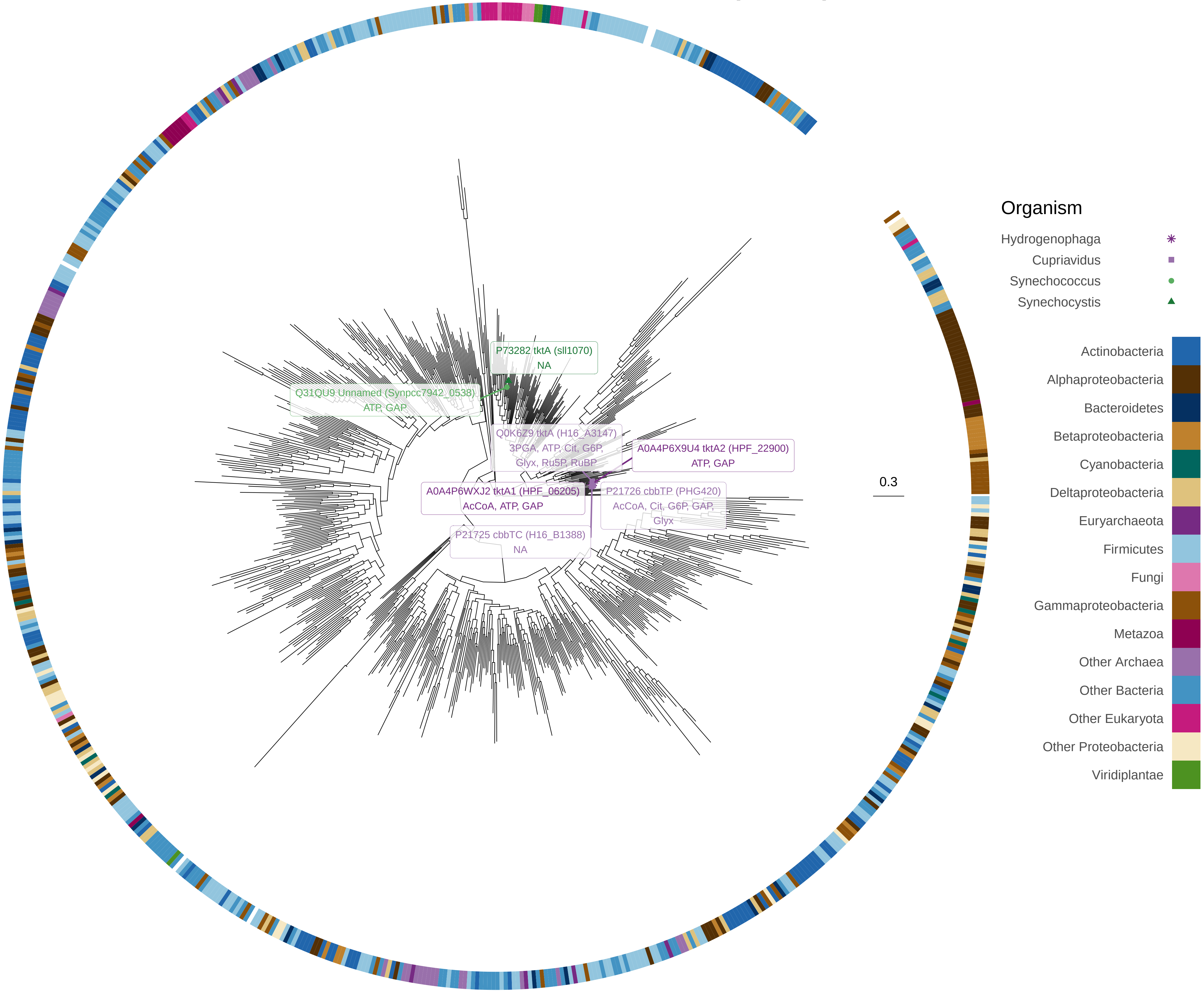

TPI

K01803 TPI, tpiA; triosephosphate isomerase (TIM) [EC:5.3.1.1]

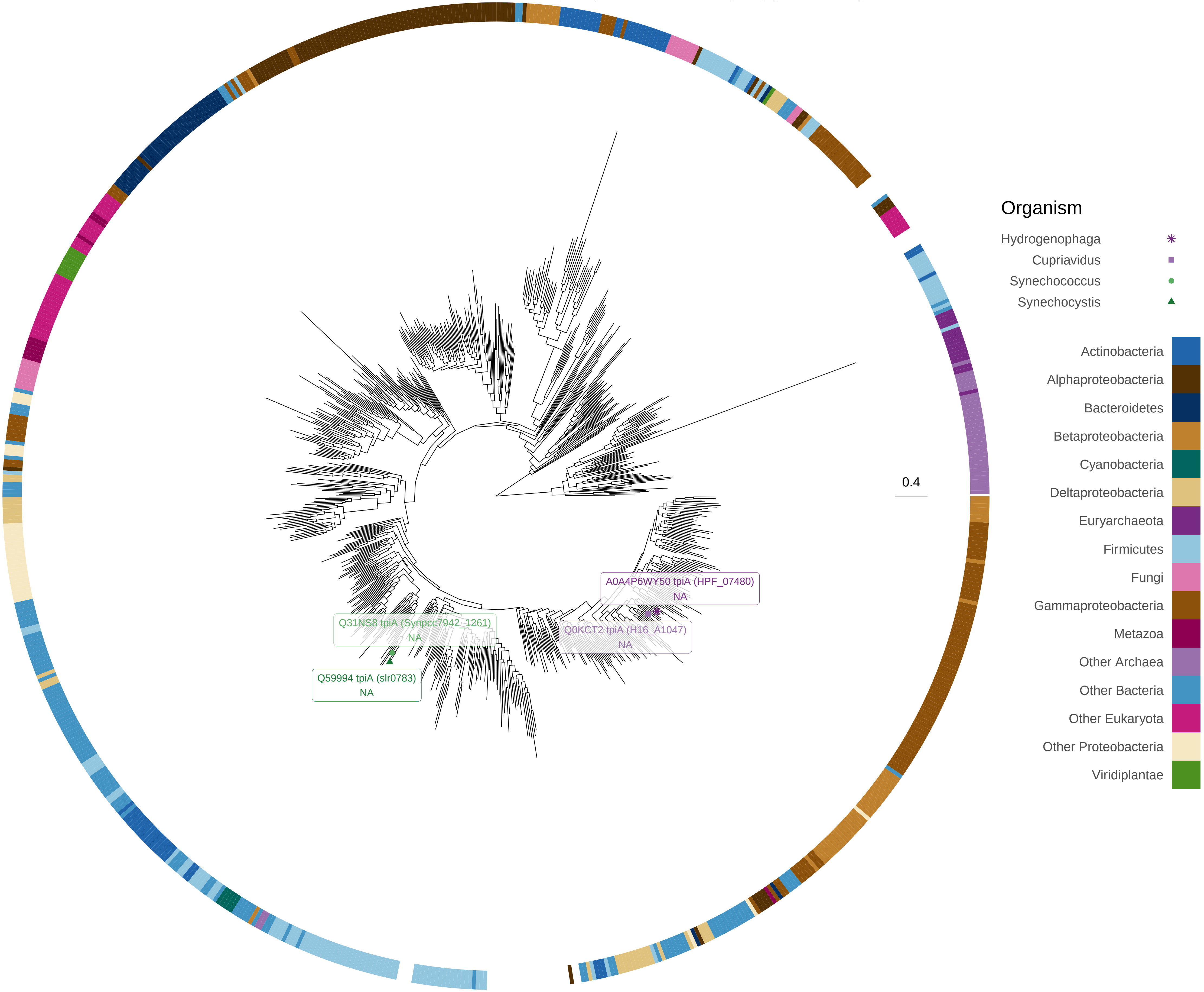
